# Rules, hypergraphs, and probabilities: the three-level analysis of chemical reaction systems and other stochastic stoichiometric population processes

**DOI:** 10.1101/2023.12.11.571120

**Authors:** Eric Smith, Harrison B. Smith, Jakob Lykke Andersen

## Abstract

We consider problems in the functional analysis and evolution of combinatorial chemical reaction networks as rule-based, or three-level systems. The first level consists of rules, realized here as graph-grammar representations of reaction mechanisms. The second level consists of stoichiometric networks of molecules and reactions, modeled as hypergraphs. At the third level is the stochastic population process on molecule counts, solved for dynamics of population trajectories or probability distributions. Earlier levels in the hierarchy generate later levels combinatorially, and as a result constraints imposed in earlier and smaller layers can propagate to impose order in the architecture or dynamics in later and larger layers. We develop general methods to study rule algebras, emphasizing system consequences of symmetry; decomposition methods of flows on hypergraphs including the stoichiometric counterpart to Kirchhoff’s current decomposition and work/dissipation relations studied in [1]; and the large-deviation theory for currents in a stoichiometric stochastic population process, deriving additive decompositions of the large-deviation function that relate a certain Kirchhoff flow decomposition to the extended Pythagorean theorem from information geometry. The latter result allows us to assign a natural probabilistic cost to topological changes in a reaction network of the kind produced by selection for catalyst-substrate specificity. We develop as an example a model of biological sugar-phosphate chemistry from a rule system published in [2]. It is one of the most potentially combinatorial reaction systems used by biochemistry, yet one in which two ancient, widespread and nearly unique pathways have evolved in the Calvin-Benson cycle and the Pentose Phosphate pathway, which are additionally nearly reverses of one another. We propose a probabilistic accounting in which physiological costs can be traded off against the fitness advantages that select them, and which suggests criteria under which these pathways may be optimal.

## I. INTRODUCTION: FROM RULES AND MICRO-STATISTICS TO MACRO-PHENOMENA IN COMPLEX COMBINATORIAL SYSTEMS

### The three-level architecture of many combinatorial and complex stochastic systems

Systems that may be defined by quite complicated and heterogeneous components and interactions may nonetheless be tractable to analysis if their state-spaces and event-spaces do not grow too large to organize and navigate [3, 4]. At a different extreme, systems with very large state- and event-spaces produced by combinatorial composition may be impractical to represent explicitly yet nonetheless remain tractable if these spaces have high symmetry and not-too-much structural diversification [5– 7]. It is at the interaction of these two forms of “largeness” – where heterogeneous mechanisms and interactions act combinatorially to generate large state- and event-spaces, that we encounter many frontiers of complexity [8].

One class of complex systems for which a broad approach has come forth are the so-called *rule-based systems* [9–15]. For many of these, the phenomena of interest are realized as stochastic processes, in which case we also refer to them as “three-level” systems. The first level consists of “rules” that abstract the classification and transformation of objects of some kind. Formal models of rules, together with prescriptions for applying them to particular objects while preserving the contexts in which they act [13, 14, 16], enable the inductive construction of object and event types by recursive rule application, which then furnish a second level of description. For stochastic phenomena, the models of literal objects and events, together with choices about the possible states in which they can occur, can take on the interpretation of generators of stochastic processes [17] producing trajectories and probability dynamics in the state spaces that constitute a third level of description.

In this paper we demonstrate a program of analysis for the phenomena generated by a three-level system, with an emphasis on the interaction of heterogeneity in the generating structures and combinatorial interactions among them, both from rules to graphs and from graphs to probability dynamics. We are interested in the way specification of the system – by us as model-builders or by constraints imposed physically or emerging from selection in nature – may be introduced at one level in the hierarchy and then propagate across levels to create order in the output at other levels. This propagation may occur by restricting combinatoriality, in which case we may be able to derive its consequences by direct arguments, or it may guide combinatorial expansion that drives emergence and robustness of collective phenomena through laws of large numbers. We will exhibit cases of each, and study how jointly they enable the design and control of function in complex combinatorial systems.

### The stochastic stoichiometric population processes among the three-level systems

For simplicity of approach, our treatment will apply to what we term *stochastic stoichiometric population processes* (SSPPs): discrete-state stochastic processes in which objects are counted as individuals grouped into types, with states corresponding to their joint counts in populations, and in which each elementary event removes some set of individuals from a population and replaces it with some other set. The SSPPs are an expressive abstraction in that they are known to include phenomena with computationally complex search and optimization problems [18], yet they are defined from a small collection of primitives [19, 20] and admit numerous topological [21] and other [15, 20, 22] modes of analysis.

The SSPPs are well known and developed as models of chemical reaction networks (CRNs) [1, 23–28], but we refer to them in general form to emphasize that they have much wider applications including Darwinian populations and genetic lifecycles [29–31], and to many of these the narrow and restrictive assumptions about dynamics that are natural for elementary CRNs do not apply [32].

The generative relation of rules in the form of graph grammars to stoichiometric chemical networks has been studied extensively by Andersen *et al*. [2, 13, 16, 33–35]. Separately, the generative relation between the topology of stoichiometric systems and their physical thermodynamics and more general large-deviation behavior has been studied by several authors [1, 23, 24, 26–28, 32, 36–40]. Therefore we will focus here on phenomena that draw jointly on properties at all three levels, and on the combinatorics both of rules and of stochastic events.^1^

### What questions arise for dynamics that is both combinatorial and complex?

The exemplar for the interaction of system-defining structure with large-numbers combinatorics, for both phenomenology and theory, has been the relation between internal-energy landscapes and entropy in equilibrium thermodynamics [6, 42–44]. With suitable generalizations to non-equilibrium contexts, that relation will arise in the presentation here, between the stoichiometric graph and its large-deviation statistics [32, 45, 46]. For rule-based systems, a second interplay of generating structure with combinatoriality arises between the rule algebras and the graph with its associated spaces of states and transitions [11, 13, 14].

We will apply our general constructions to an example problem from the biochemistry of sugar-phosphates, which is unusual for the high combinatoriality achievable from a small set of generating mechanisms [2], and for the degree to which repeated rule use has been retained in biology and has been essential to achieving networks that recombine sugars without material loss and with modest dissipation.

In terrestrial evolution we find the emergence of a single, ancient, universal pathway for sugar rearrangement in the anabolic Calvin-Benson-Bassham cycle [47] through substrate specificity of catalysts, from a formally-infinite chemical network that could diffusely perform the same conversion. Moreover, the Calvin-cycle pathway is nearly reversed in direction to form a second ancient, nearly-universal pathway in the catabolic Pentose-Phosphate pathway [48–53]. We want to account both for how the two functions of catalysts – realizing reaction mechanisms and specifically restricting substrates [54, 55] – relate in the process of pathway evolution, and also why these particular pathways are the outcomes.

Three questions we wish to answer are: 1) Which aspects of pathway specificity derive from rule properties and which from ancillary network restriction? 2) How are macroscopic transport capabilities and force/flux response that might be targets of selection attributable to lower-level and distributed catalyst specificities that can be modified to select them? and 3) What cost functions might be assignable that bridge the functional properties of the network as a transport mechanism and the informational attributes of that network as a product of selection?

### Technical and general-method results

A variety of more specific methods and results are derived that contribute to the foregoing integrated, biologically-motivated questions, but support the analysis of SSPPs as three-level systems more generally:

For rules we study the presence of discrete symmetries and resulting conserved quantities that constrain all conversions that can be performed by composition of rule actions. We show that certain optimality properties such as shortest path length and a topological characteristic related to minimization of dissipation can be computed from the rule algebra alone without the intermediate step of explicit construction of stoichiometric networks through network expansion and subsequent exhaustive enumeration of integer-flow solutions on the resulting networks.

For stoichiometric networks we study the generalization of Kirchhoff’s current decomposition laws from electric circuit theory [56]. The systematic construction of bases of null flows of stoichiometry has been shown before to be useful for analysis of dissipation [37]. In our graph-grammar model it will enable a complete construction of the null space in terms of a fixed class of basis types for networks of indefinite size.

We situate our analysis of work and dissipation in a somewhat more general study of nesting hierarchies for graphs than is done by Wachtel *et al*. [1], because our concern is less with defining and computing transduction and efficiency (already fully performed by them), than with deriving all concepts within the representational abstraction of the hypergraph. Where Wachtel *et al*. invoke several *ad hoc* criteria specific to chemistry (and only for “elementary” reactions with all species kept explicit) – such as their class of admissible *chemical transformations* defined in terms of element conservation, and dissipation derived inherently in terms of energy and the local detailed balance commonly assumed in stochastic thermodynamics [57, 58] – we obtain all notions of system/environment relations and their jointly performed transformations solely from stoichiometry and graph embedding, and entropy decompositions from large-deviation functions. We do this to emphasize that the SSPPs are a much wider class to which these abstractions can be applied than to CRNs with energy conservation [32], and that even within chemistry a variety of different coarse-grainings [59] may entail different conservation laws and functional forms for likelihood within a standard modeling framework.

In the dynamics of probability under the master equation, we re-present the work-dissipation identities derived in [1, 24], and derive the large-deviation function for currents. In particular we show that the LDF possesses an additive decomposition in the form of the extended Pythagorean theorem of information geometry [60, 61], for current solutions under the mass-action rate law in hierarchies of nested graphs.

#### A. Organization of the presentation

Sec. II introduces SSPPs as a model class and their topological representation by hypergraphs. The stoichiometric counterparts to Kirchhoff current laws are derived for steady states, along with related identities for chemical work delivery and dissipation noted in [1]. Definitions of system/environment decomposition for graphs, and reducibility or irreducibility of flows on graphs, are given in terms of graph topology and stoichiometry alone. These will serve as a basis to define boundary conditions for driven networks and to formally distinguish work transduction through stoichiometric and non-stoichiometric coupling (termed “tight” and “loose” coupling in [1], and treated there only through examples).

Sec. III introduces the second generative relationship making a CRN a three-level system: from an underlying rule algebra representing chemical reaction mechanisms by graph grammars [13, 16] to the set of chemical species and the hypergraph of reactions that defines the SSPP. Here we introduce a graph-grammar model of sugar-phosphate chemistry from [2], and a thermochemical landscape produced from high-throughput computational methods [62]. From these we can cast the emergence of a universal Calvin cycle and Pentose-Phosphate pathway as a problem of selection from a combinatorially large network faced historically by biological evolution.

We frame the problem of realizing a feasible chemical conversion on a large network in terms of integer linear programming, solved by exhaustive enumeration of all stoichiometrically independent integer flows performing that conversion on the graph. We then show how rules can be used to decompose the space of null flows for graphs of any size, and how conservation laws at the rule level propagate to constrain solutions at any order, leading to derivations of solution minimality that circumvent a need for exhaustive enumeration.

Sec. IV derives the large-deviation theory for concentrations and currents on general SSPPs, and applies it to flows in the null space from the example of Sec. III. We show first how the large-deviation function (LDF) for steady-state currents is a time integral in which the integrand relates to Liouville flows [63] and their associated Hamilton-Jacobi theory for large deviations [64–66], and recapitulates a mathematical structure seen in the simpler LDF for concentrations in a single-time distribution. We then show how the information geometry induced by the Kullback-Leibler divergences of mass-action currents on a graph and any of its subgraphs leads to the additive decomposition by the extended Pythagorean theorem, following the same partition used to decompose Kirchhoff components of null flows in Sec. II.

Although only a simplifying case and thus not a general result, we study the limit of linear response near a thermodynamic equilibrium, and show how a measure of network resistance to a feasible conversion can be defined for stoichiometric systems, and how this is dominated by contributions from graph topology shown in Sec. III, for realistic thermochemical landscapes governing the actual Calvin cycle and Pentose Phosphate pathway. Finally we make the argument that for nested graphs as a model representation for biological selection for catalyst specificity, the LDF for currents with its Pythagorean decomposition is a natural cost function, nesting in the manner of partition functions, and compatible as a summand in a cost function for selection in biological populations also modeled as SSPPs.

## II. THE STOCHASTIC STOICHIOMETRIC POPULATION PROCESSES AS A MODEL CLASS

We begin with the definition of stochastic stoichiometric population processes (SSPPs) as the modeling abtraction. In *population processes*, objects are regarded as individuals grouped into *P* (finite or countable) types, and the states refer to populations indexed by the counts of their members by type. States are then lattice points in the non-negative orthant of ℤ^*P*^. Change events are *stoichiometric*, meaning that in each elementary event some set of individuals, specified by type-counts, is removed from the population and some other set is added to produce the next state of the population.

### A. Declaring a model (as far as the generator)

We intend that all features of the system be made explicit within the limited terms provided by the model abstraction. Thus, for instance, if a graphical model is used for a chemical reaction network, and we wish to consider “environments” of the reaction system that extract some molecules and supply others, we do not wish to appeal to *ad hoc* criteria outside the declared model for what combinations of sources and sinks are admissible or inadmissible; these should be derivable from conservation laws made explicit in the model declaration.

While the examples in this paper will be derived from chemical reactions with microscopic reversibility, a class with many physical constraints that limit their complexity, we wish to derive relations that extend within the same modeling abstraction to very differently constrained problems such as population genetics [30], strictly irreversible processes [32], or coarse-grained effective theories that may absorb buffered species [59] from the medium within effective rate constants and thereby break some microscopic conservation laws while still preserving others. Conversely, biochemical models [67] may lack key conversions and thereby preserve larger-scale motifs, with vitamins in organism physiology serving as an important example. This subsection presents the commitments needed to declare an explicit model.

#### a. Populations

The species types in a population are given an index *p* ∈ 1, …, *P*. In reaction schemata, species themselves (molecule types) are written as a (column) vector of formal labels *X* ≡ [*X*_*p*_]. The state of a system corresponds to the counts of all its member species; we write these as a (column) vector of non-negative integer coefficients n ≡ [n_*p*_].

Probabilities over population states n are written as column vectors *ρ* ≡ [*ρ*_n_] with ∑_n_ *ρ*_n_ = 1. We will represent stochastic processes on the population by the flow of probability under a continuous-time, discrete state master equation [63] of the form

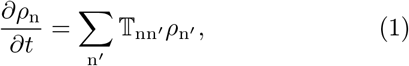

in which 𝕋 ≡ [𝕋_nn′_ ] is a representation of the generator of the process called the *transition matrix*. 𝕋 is a stochastic matrix, meaning 1^*T*^ 𝕋 ≡ 0.

#### b. Stoichiometry

The key property of stoichiometric population processes that gives transitions a structure peer to, and independently specifiable from, the type structure of species is the set-wise removal and replacement of population members in elementary events. To capture this structure topologically we introduce an independent class of objects from species, termed *complexes* following Horn and Jackson [19] and Feinberg [20], and denoted with an index *i*.

Whereas species types *p* admit arbitrary counts n_*p*_ that carry the system state through time, complexes are fixed multisets of species that correspond to samples from the population state. The type membership in complexes is represented in a matrix *Y* with columns 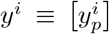, giving the count of each type *p* in complex *i*, known as the *stoichiometric coefficients*.

*Reactions* are written directionally, and every reaction connects an ordered pair of complexes. To simplify the notation for indexing in this paper, we will suppose that there is at most one reaction in either direction between a unique pair of complexes, so that reactions can be labeled with ordered pairs (*ji*), for a reaction that removes complex *i* and adds complex *j* in its place.

A collection of reaction schemata specify a stoichiometric model that we write as a *hypergraph* [21] that we designate 𝒢 and identify with the set of all its reactions:^2^

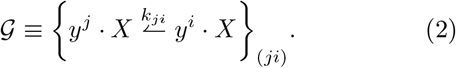

A dot product of two column vectors denotes the formal sum 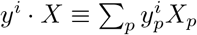.

#### c. Rates

Reaction rates correspond in the stochastic process to probabilities per unit time for each transition event to occur, which are functions of both the samples defined by complexes and rate parameters. Here we adopt the simplest case of proportional sampling of complexes from the population without replacement. The number of possible samples of complex *i* from a population in state n defines a complex *activity* of the form

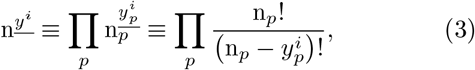

where we use the underbar notation for the (component-wise) falling factorial [25]. The *half-reaction rate* for reaction (*ji*) is a product of the activity (3) of the input complex *i* with a rate constant *k*_*ji*_, used to label the reaction transition in schema (2).

Since the elements of the transition matrix 𝕋 ≡ [𝕋_nn_′ ] have a closed form in terms of the much smaller collection of activities (3) and rate constants, it is convenient to write the transition matrix as a projection operator that performs index shifts as part of the definition of the matrix elements. As a notation for this representation, we use the exponential form 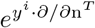 for the shift operator n → n + *y*^*i*^ acting to the right.^3^ Incorporating the shift operator representation, the transition matrix for the stochastic process on the graph (2) requires only a sum over reactions:

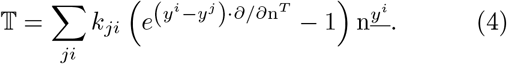

The schema (2) defines an adjacency/rate matrix on complexes that we designate 𝔸 ≡ [𝔸_*ji*_] with 𝔸_*ji*_ = *k*_*ji*_ ; *j* ≠ *i*. 𝔸_*ii*_ = −∑ _*j*≠*i*_ *k*_*ji*_, and in terms of this matrix the generator (4) can be written in a form that expresses a certain formal symmetry between the removed and emplaced complexes as

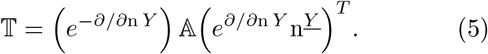

##### 1. First-moment dynamics and mean-field approximation

The probability vector *ρ* is (in general) infinite-dimensional, and solution of its dynamics will generally be carried out only in approximation. The lowest-order approximation comes from the first moment which we denote with angle brackets:

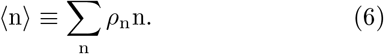

First-moment dynamics underlie the deterministic rate laws for average behavior but not only that: In the construction of cumulant-generating functions and functionals, and their Legendre transforms, first-moment dynamics arise also as rays (also called “eikonals”) [69, 70] for the leading contributions to the probabilities of rare large deviations, computed within a Hamilton-Jacobi theory [32, 66] derived from the generator (5).

For now we note only the deterministic rate equation from the generator (4), which in the large-n approximation gives the *mass-action rate law* :^4^

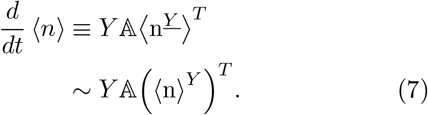

In the product-form steady states produced by complex-balanced networks [22], which we term “A CK” states, the factorial moments evaluate to powers 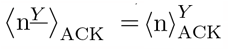, and the mass-action law (7) is exact.^5^

We introduce a notation *v* ≡ [*v*_(*ji*)]_ for a column vector of net currents through reactions (signed according to the index order *ji*). The general form and mean-field approximation for reaction (*ji*) under the rate law (7) are given by

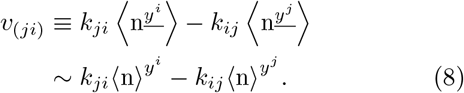

Like the transition matrix 𝕋 on states, the adjacency matrix 𝔸 is stochastic on complexes, meaning 1^*T*^ 𝔸 ≡ 0. This is so because the complex network is a simple network, and each current *v*_(*ji*)_ multiplies a difference of column vectors *y*^*j*^ − *y*^*i*^. We denote the *stoichiometric matrix* between currents and species as 𝕊 ≡ [𝕊_*p*(*ji*)_] with column

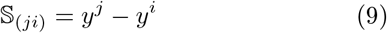

corresponding to reaction (*ji*). The first-moment equation (7) is then expressed directly in terms of currents as

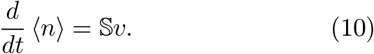

Eq. (10) remains an identity whether or not currents come from the mass-action rate law, a relation that will be important later when computing large-deviation functions for currents.

##### 2. Subcase: rate parameters from state variables in some thermal-equilibrium context

For the special case of networks with detailed-balance steady states, the half-reaction rate constants can be expressed in terms of a pair of potentials, one defined for species and the other for reactions. For applications to chemistry, we introduce a vector of one-particle chemical potentials for species, 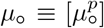.^6^ From single-molecule chemical potentials we construct a chemical potential for any complex *i* from its corresponding stoichiometric vector as 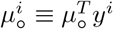. Finally, we introduce a vector of chemical potentials 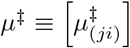 that we may think of as “one-complex” chemical potentials at the transition state. In terms of these, the half-reaction rate constants are given by

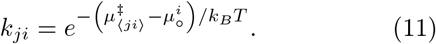

where *k*_*B*_ is Boltzmann’s constant and *T* temperature. The corresponding equilibrium constants are

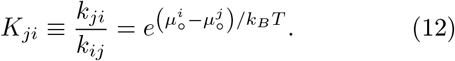

Denote by *<u>n</u>* ≡ *<u>n</u>*_*p*_ the mean number vector in the steady state (which is an ACK distribution for which *<u>n</u>* are the Poisson parameters [22]). *<u>n</u>* depends on the equilibrium constants (12) and on the values of any quantities conserved under the stoichiometry from 𝕊, but not on the transition-state potentials *μ*^‡^.

We may introduce an adjacency matrix with rate constants descaled by the equilibrium complex activities, 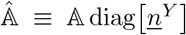, in terms of which the generator (5) becomes

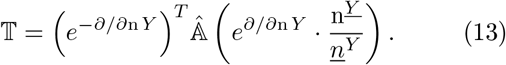

The full 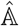 equals a sum of terms that on each pair of complexes {*j, i*} connected by a reaction have the forms

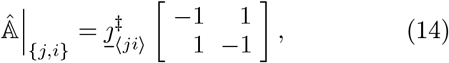

in which

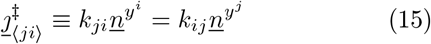

is the magnitude of the equal and opposite half-reaction currents through the transition state which cancel at equilibrium.

##### 3. The near-equilibrium linear-response regime

For systems displaced weakly from a detailed-balance equilibrium, the currents (8) may be expanded to linear order in mean-field approximation, as

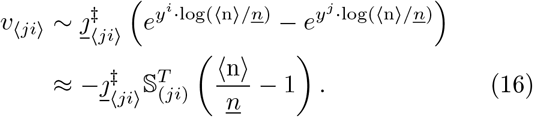

We collect the components of the vector *v*, and write their resulting aggregated source and sink currents for columns of *Y* in Eq. (7), as

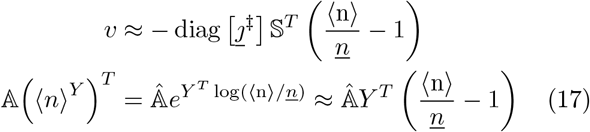

Comparing the forms (7,10), and using the expansions (17), we see that the first-moment equation in linear response is relaxation under a symmetric matrix

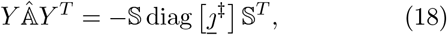

which depends only on the stoichiometry and the transition-state currents at equilibrium.

### B. The generator as the middle level in 3-level rule-based systems

The foregoing construction applies to any stochastic stoichiometric population process, providing, for example, a model of populations of literal molecular species and literal reactions among them. For many phenomena, however, the literal reaction network is neither the most compact nor the most complete reflection of our knowledge of the process at work. Chemical reactions are grouped into equivalence classes in terms of common reaction *mechanisms* [72], so named because they perform specific conversions of chemical bonds among conserved atomic centers, which together are generally only fragments of whole molecules.

Here we will be interested in stoichiometric systems that are not merely declared *ad hoc* but are themselves generated by specified mechanisms. The generalizations from the particular chemical notion of reaction mechanism will be termed *rules*, and the approach to creating state spaces, networks, and dynamics from rules is called *rule-based* modeling [12]. We will also term phenomena described in this way **three-level systems**, for the hierarchical levels with generative relations between them: those of *rules*, stochastic process *generators* specified as in Eq. (4) from the topology of the hypergraph, and *ensemble dynamics* on the state space or its bundle of histories.

For chemistry a natural intermediate level of abstraction, known as a *graph grammar* [16] represents molecules as ordinary graphs in which atoms are (typed) nodes and bonds are (typed) links. Reaction mechanisms correspond to rewriting rules for graph fragments, which retain atomic centers and reconfigure bonds. The mathematical relation bridging the (lower) level of rules and the (higher) level of stoichiometric reaction networks is the structure-preserving embedding, of the graph fragment altered under the reaction mechanism within the full graph of a molecule. The parts of the input molecules not altered by the reaction mechanism constitute a context for the atomic centers acted on by the rule, which is propagated through the rewrite to produce new output molecules. Structure-preserving embedding is realized in a formal rule-based language as a double-pushout from category theory [13, 16].

By recursive application of all rules from some set to a starting set of seed molecules, a stoichiometric graph can be formed, in which each reaction is an image of the generating rule. Chemical species have no such simple mapping, as each species (except starting inputs) is created through the joint action of the rule as a creator of bond patterns, and the pushout that embeds the reacting centers in molecular contexts. The notion of an algebra of rules arises because both the embedding step and the rewriting step may create or destroy instances of graph patterns to which other rules apply, so that the application of rules can generally be non-commutative [73].

Each level-crossing in a three-level system produces a one-to-many map, which may map finite domains to either finite or indefinitely large ranges. Thus each rule in a finite set of mechanisms may be realized in indefinitely many literal reactions in the generated hypergraph, and and each reaction in a hypergraph may govern indefinitely many explicit transitions in the lattice of states.

### C. Stoichiometry and other aspects of hypergraph topology

From the stoichiometric matrix 𝕊 defined in Eq. (9), three important subspaces are defined that govern the conservation and dynamics of matter, currents, and probability, under any stochastic model and at any rate parameters and particle content. From properties of the adjacency matrix 𝔸 from Eq. (5) that defines the network of reactions between complexes, results from [19, 20, 22] provide a surprisingly strong and restrictive classification of steady states. This section reviews the needed constructions of those two results.

#### 1. Null spaces of the stoichiometry: conserved quantities and balanced flows

The first decomposition concerns the rank and nullity of the stoichiometric matrix 𝕊.

##### a. Right null space

Let *v* denote a (column) vector of reaction fluxes on the graph 𝔾, and *μ* a (row) vector of chemical potentials. Then the case

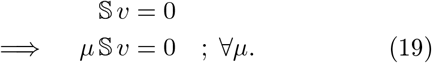

Such *v* are in right null space of 𝕊. They cannot affect *d*⟨*n*⟩ */dt*, and for any potential *μ* ≡ [*μ*_*p*_], the second line of Eq. (19) means that *v* as a whole is non-dissipating. Denote the space spanned by solutions *v* to Eq. (19) as ker 𝕊.

##### b. Left null space

For any (row) vector *c* ≡ [*c*_*p*_] of real numbers with the property

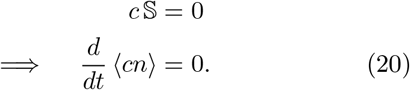

The component *c*_*p*_ is the measure of the *conserved quantity c* ascribed to species *p* by the stoichiometry. The second line of Eq. (20) will be true for any vector of fluxes *v* causing change of *n*. Because the coefficients of 𝕊 are integers, a basis spanning solutions (20) (for finite 𝕊) can always be found in vectors over the integers. Denote the space spanned by the solutions *c* of Eq. (20) as (ker 𝕊 ^*T*^)^*T*^.

The image of 𝕊, denoted im 𝕊 = im (*Y* 𝔸), is called the *stoichiometric space*. We denote its dimension by *s* ≡ dim im 𝕊, which is the rank of 𝕊. It follows that *s* + dim (ker 𝕊 ^*T*^)^*T*^ = |{*p*}|, the number of species, and that *s* + dim ker 𝕊 = |{⟨ *ji*⟩ | *k*_*ji*_ or *k*_*ij*_ ≠ 0}|, the number of reactions.

#### 2. The complex network and model classes with respect to balance

The relation between the left null spaces of 𝔸, and of *Y* 𝔸, along with the symmetry or non-symmetry of 𝔸, separate steady states under stochastic process generators of the form (4) into three distinct classes according to whether their flows balance within the complex network, between each pair of complexes within that network, or only on species but not on complexes. For two of these, uniqueness results and remarkably restrictive functional forms are also known [20, 22] for steady states along with the existence of Lyapunov functions for the first-moment parameters.

##### a. Complex balance

the property of the steady states in this class is ∑_*j*_ *v*_(*ji*)_ = ∑_*j*_ *v*_(*ij*)_; ∀*i*. The topological condition on the graph ensuring this property is ker (*Y* 𝔸) = ker 𝔸.

**Consequences:** There is a unique interior fixed point (all components positive) of the mean-field equations (7) [20] at any values of the generating parameters {*k*_*ji*_}, which we denote *n*, and the associated steady-state solution to the master equation is either a product of Poisson distributions for *n*_*i*_ or the restriction of such a product to a linear subspace [22]. For these distributions the mean-field approximation of the rate law (7) at large *n* is also exact. For ⟨*n*⟩ = *e*^*θ*^*n* the mean-value parameter for any other non-steady-state distribution under the same large-*n* approximation, the *f* -divergence [74]

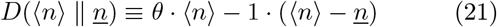

is the large-deviation function (see Sec. IV for constructions) and the Lyapunov function under the stationary-path equations of the Hamilton-Jacobi theory (see [32] for more extended discussion in relation to large deviations).

##### b. Detailed balance

the property of the steady states in this class is that *v*_(*ji*)_ = *v*_(*ij*)_; ∀ reactions (*ji*). The requirement on rate constants in addition to the topological condition for complex balance, that they be expressible as in Eq. (11), implies that 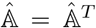. The fact that 𝔸 and thus 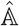 are stochastic on complexes then means 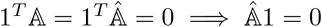.

**Consequences:** In addition to having the *f* - divergence (21) as the Lyapunov function common to all complex-balance cases, for detailed balance the eikonals [69, 70] that carry the dominant probability for the tails of the large-deviation function, which are constructed as *escape paths* in the Hamilton-Jacobi construction [32], are time-reverses of the classical relaxation trajectories in the Hamilton-Jacobi theory.

The general case with ker (*Y* 𝔸) ≠ ker 𝔸 is subject to neither of the foregoing restrictions of uniqueness or functional form, and can require non-local solution of all moments throughout the state space.

#### 3. Complex balance and deficiency

Complex balance entails very strong restrictions and simplifications on steady states relative to the complexity of the general case, so not surprisingly, it is not a generic condition for most stoichiometric graphs. Apart from achieving complex balance by fine tuning of rate constants (a subset of measure zero even when it is possible), one strong topological condition is known that implies complex balance much more generally.

The topological index termed *deficiency* is denoted and defined as *δ* ≡ *C* − *l* − *s*, where *C* is the number of complexes, *l* is the number of connected components (called “linkage classes” in the CRN literature) of the complex network defined by 𝔸, and *s* as above is the dimension of the stoichiometric space. For *δ* = 0 (with other technical conditions that all linkage classes are “strong terminal” linkage classes [20, 25, 75], ensuring that every complex can be recurrently visited) the steady states will be complex-balanced at all values of the rate constants.

### D. Graphs and their subgraphs

We want the stoichiometry of a model to contain all our commitments to the “kinematics” of the system modeled: the types of its species and its transitions and the relations between them; prescriptions for decomposing a large system into a focal subsystem and its environment within the larger system; and specifications of those transformations considered feasible both within the sub-system and in its environment.

From a graph’s topology we may perform a Kirchhoff-like decomposition of the linearly independent flows on and through it. Subsequently specifying a dynamical model as in Sec. II provides constructions for chemical potentials dual to species, and from these and the Kirchhoff decomposition we can derive accounting identities between chemical work and dissipation, and then transduction and efficiency following [1]. From the particular study of null flows on graphs, we may derive criteria for flow reducibility or irreducibility that determine the response of thermodynamic flows to thermodynamic forces, and we may define a systematic algorithm for flow reduction that also decomposes the dissipation and the large-deviation rate functions for currents defined in Sec. IV.

Here in a series of small subsections, we introduce the necessary operations on graphs and on the flows that they support, making particular use of the null spaces of the stoichiometry on an overall graph that will be the context for all subsystems of interest. Work-dissipation identities are noted, and transduction through both stoichiometric and non-stoichiometric coupling defined in terms of null flows. The subsection closes with special forms in the limit of linear response, where many properties of interest for resistance or efficiency are defined only from graph topology and thermodynamic equilibrium states, without reference to the magnitude of non-equilibrium forcing.

#### 1. Internally-balanced flows and through-flows of a subgraph

A *flow* on a graph is any assignment of currents to reactions in the graph. In this section we will denote flows *v*, but they are not restricted to be solutions to the mass-action law (or any other law) as in Eq. (1) above. *Balanced flows* [16]^7^ are those that do not result in changes of species counts. They are arrived at in general as solutions to an integer linear program, possibly in combination with currents of species onto or off the graph. By construction the flows in the right null space of the stoichiometric matrix of a graph are balanced in the absence of any external source or sink currents, and so we will term these *internally balanced flows*, as well as *null flows*. Rather than construct balanced flows in relation to arbitrary source or sink currents for species, we will construct all such flows from the null space in some outer graph, so that their span is derived from the graph stoichiometry.

Starting from a graph 𝒢, the following containment and restriction relations may be defined, and a few small accounting lemmas checked:

**Subgraph of a graph:** denoted 𝒢′ ⊂ 𝒢, it is defined to be a subset of the reactions in the set (2) defining 𝒢. We include all species present in any complexes in the reactions in 𝒢′ and exclude any species in 𝒢 not included in any complexes in 𝒢′.

**Complement to a subgraph:** denoted (𝒢′)^⊥^ ⊂ 𝒢, it is defined to be the graph generated by the reactions in 𝒢 and not in 𝒢′. Species in any complexes in the reactions of the complement are included in the complement. In applications we will often refer to the complement of a subgraph as its *environment* within 𝒢.

**Boundary of a subgraph:** The species in 𝒢′ ∩ (𝒢′)^⊥^ ≡ *∂*_𝒢_ 𝒢′ define the *boundary* of 𝒢′ in 𝒢.

**Restriction of a stoichiometric matrix to a sub-graph:** Let 𝕊′ ≡ 𝕊|_𝒢′_ denote the restriction of 𝕊 to the species and reactions in 𝒢′. Then (ker 𝕊^*T*^) |_𝒢_′ ⊂ ker (𝕊′)^*T*^ : 𝒢′ conserves all quantities conserved in 𝒢, and may conserve more. Conversely ker 𝕊′ ⊂ (ker 𝕊)|_𝒢_′ : null flows within 𝒢′ are null within 𝒢, but there can be null flows in 𝒢 that are contained outside, or that pass through, 𝒢′.

**Conservative flows through a subgraph:** The sub-space (ker 𝕊′)^⊥^∩(ker 𝕊)|_𝒢_′ – flows that are non-null within 𝒢′ but are the restriction of null flows in 𝒢 – will take the place in the constructions below of what [1, 24] term *emergent cycles*. By construction they do not change concentrations of species within the interiors of both 𝒢′ and (𝒢′)^⊥^, meaning that all fluxes are balanced within either subgraph except on its boundary nodes, and the fluxes at those nodes from the subgraph and the complement are likewise balanced.

**Subgraph-specific conservation laws:** Note that (im 𝕊)| _𝒢_′ ∩ ker (𝕊′)^*T*^ are quantities conserved by all reactions in 𝒢′ but changed by some reaction within 𝒢. We will generally not make use of them, because they are simply part of the conserved quantities determining stoichiometric equivalence classes within the subgraph, and internally-balanced flows within 𝒢 cannot change them in (𝒢′)^⊥^.

#### 2. Supporting graph of a flow and definition of flow irreducibility

Let *v* be a flow on 𝒢.

##### a. Species current and boundary

Denote by *J*(*v*) ≡ 𝕊*v* the vector of net currents to all species nodes resulting from flow *v*. Define the *boundary* of *v* as *∂*_𝒢_ *v* ≡ {*p* | *J*_*p*_(*v*) ≠ 0}. If *v* is a null flow then *∂* _𝒢_ *v* = ∅.

##### b. Support and irreducibility

Define the support of *v*, denoted supp *v* ⊂ 𝒢, to be the subgraph of 𝒢 consisting of the reactions on which *v* is nonzero. We will call *v irreducible* if dim ker 𝕊|_supp *v*_ = 0: removal of any reaction from supp *v* results in a graph that cannot support the net conversion (if any) performed by *v*.

If dim ker 𝕊 |_supp *v*_ *>* 0 then *v* is *reducible*. For any reaction (*ji*) in the support of a null flow in ker 𝕊|_supp *v*_, the magnitude of that null flow may be chosen to set the flux through (*ji*) to zero, resulting in a flow on supp *v* − (*ji*). In this way the supporting graphs for one or more irreducible flows may be extracted from the supporting graph of a reducible flow. We will call any sequence of such removals of reactions from the supporting graph of a reducible flow *v* to reach the supporting graph of an irreducible flow a *reduction* of *v*.

**Remark:** An irreducible flow *v* is fully determined by the supporting graph supp *v* and the net conversion *J* = 𝕊*v* on *∂* _𝒢_*v*, so we may write *v* = *v*(*J*). There are no current degrees of freedom left undetermined within the stoichiometric space of species conversions im 𝕊 |_supp *v*_, and under the master equation (1), all species degrees of freedom in the stoichiometric space are determined as functions of the fluxes in *v*.^8^ A reducible flow includes one or more null current components that do not change species numbers and that are left undetermined by *J* = 𝕊*v*. These components may vary across different solutions to the mass-action rate equations, and their large-deviation values will allow us to associate improbabilities with the reaction eliminations that produce equal deviations during a flow reduction sequence.

**Corollary:** Since stoichiometry is discrete, the flux components in an irreducible flow are all integer multiples of some common denominator.

#### 3. The dimension of a reaction in a null flow

Next let *v* be a null flow in 𝒢. Consider some reaction (*ji*) ∈ supp *v*. Define the **dimension** of (*ji*) within *v* to be dim_*v*_ (*ji*) ≡ dim ker 𝕊|_supp *v*_− dim ker 𝕊|_supp *v*−(*ji*)_. This dimension is always defined, because if (*ji*) ∈ supp *v* then *v* is nonzero on (*ji*) and so the removal of (*ji*) must eliminate some null flow, if only *v* in its entirety. dim_*v*_ (*ji*) counts the number of independent null flows on supp *v* that cannot exist without passing through (*ji*).

#### 4. Null flows that circulate through the boundary of a subgraph

Suppose that *v* is a null flow within 𝒢 for which supp *v*∩ 𝒢′ ≠ ∅ but supp *v* ⊈ 𝒢′. This is conservative flow passing through 𝒢′ by the definition of Sec. II D 1.

Because graphs and their complements are defined by disjoint sets of reactions, any such *v* is a sum of the restrictions of *v* respectively to 𝒢′ and its complement: *v* = *v*| _𝒢_′ + *v*|_(𝒢_′)^⊥^. By the stipulation that *v* is null, the boundaries of the two restrictions of *v* are the same and lie in the boundary of 𝒢′: *∂* _𝒢_ *v*| _𝒢′_ = *∂* _𝒢_ *v*|(_𝒢′_)^⊥^ ⊆ *∂* _𝒢_ 𝒢′. The currents from the two projections are equal and opposite: *J (v*| _𝒢_′) + *J(v*|_(𝒢′)_^⊥^)= 0.

#### 5. Work-dissipation identity for a flow through a subgraph

The conservation of energy, written as a relation between chemical work delivered and free energy dissipated to heat, takes a particularly simple form as an accounting identity between row and column sums in the stoichiometric matrix.

Let the (row) vector *μ* ≡ [*μ*_*p*_] be any chemical potential vector defined on species in 𝒢, and *v* a null flow in 𝒢 for which *v*|_𝒢_′ is a conservative flow passing through a subgraph 𝒢′ ⊂ 𝒢. Then the chemical work flow outward through the boundary of 𝒢′ is computed on species as

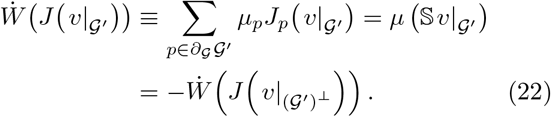

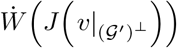 is the chemical work delivered by the environment-projection of the flow to the species in *∂*_𝒢_ 𝒢′.

The free energy dissipated to heat within 𝒢′ is computed on the reactions in the subgraph as

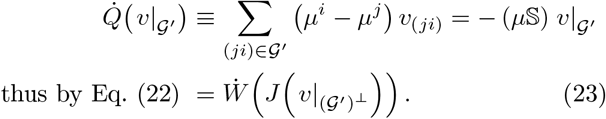

thus by Eq. (22) = 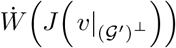

Since Eq. (23) relates the interior and boundary of a graph, it makes no reference to the complement except implicitly through any restrictions that complement may place on allowed boundary forms *J*. It will often be useful to simplify the notation below and write that for any conservative flow *v* passing through a graph 𝒢,

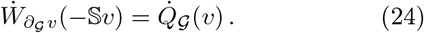

In diagrams, dissipation occurs through hyper-edges, while chemical work is transduced between subsets of stoichiometric lines to species in a graph boundary (see Fig.3 below once a specific model has been introduced).

#### 6. Redistribution of dissipation and chemical work by null cycles, and transduction by reactions

The foregoing definitions of the supporting graph of a flow, and the dimension of each reaction in that graph, provide a language in which to characterize chemical work transduction between flows, and to distinguish the stoichiometric coupling among reactions in an irreducible flow, from coupling between stoichiometrically independent components in a reducible flow by work transduction. Wachtel *et al*. [1] refer to these respectively as “tightly” or “loosely” coupled cycles.

##### a. Work and dissipation at a single reaction

Let *v* be a null flow in 𝒢. Then take (*ji*) ∈ supp *v* a reaction with dim_*v*_ (*ji*) = *d*. When a non-equilibrium chemical potential *μ* is imposed on the species in 𝒢 (for example, by external sources and sinks of species, which need not even be in supp *v*), we wish to understand the role of the null flow in redistributing chemical work and dissipation around supp *v*, and of the reaction (*ji*) in transducing chemical work among species in *∂*_𝒢_ (*ji*) (the collection of species in the complexes *j* and *i*).

Call the restriction *v*|_(*ji*)_^⊥^ *feasible* because it is a conversion within the stoichiometric subspace of the complement to reaction (*ji*) in supp *v* ⊆ 𝒢.^9^ From the definition (22), the chemical work delivered by *v*|_(*ji*)_^⊥^ to the species in *∂*_G_ (*ji*), 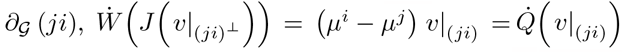, the dissipation within the reaction (*ji*) due to *v*.

##### b. Stoichiometric versus potential coupling of flows to boundary conditions

Let {*v*_1_, …, *v*_*d*_} be the elements in a basis for ker 𝕊|_supp *v*_ that satisfy (*ji*) ∈ supp *v*_*k*_ (each basis element passes through (*ji*)). There will then be a set of *d* real numbers {*α*^1^, …, *α*^*d*^} for which ∑_*k*_ *α*^*k*^ *v*_*k*_|_(*ji*)_ = *v*|_(*ji*)_. Any basis elements for ker 𝕊|_supp *v*_ that do not pass through (*ji*) are separately null in (*ji*)^⊥^, and can be ignored for transduction questions, as at most they affect the potential values *μ* of species outside *∂*_𝒢_ (*ji*) that are treated here as boundary conditions. Therefore, for simplicity, take dim ker 𝕊|_supp *v*_ = *d*, meaning that supp *v* − (*ji*) supports no null flows.

###### Chemical transduction efficiency

For spontaneous dynamics under the law of mass action under any vector of chemical potentials *μ*, 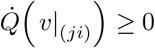.

Consider a case in which a null flow *v* on 𝒢 can be written as a sum of two flows in the complement of reaction (*ji*), *v*|_(*ji*)_^⊥^ = *v*_1_ + *v*_2_, with both *∂*_𝒢_ (*v*_1_) ⊆ *∂*_𝒢_ (*ji*) and *∂*_𝒢_ (*v*_2_) ⊆ *∂*_𝒢_ (*ji*); each component *v*_1_ or *v*_2_ is conservative on supp *v* except where it deposits or withdraws current from species in complexes *i* or *j*. Because this decomposition is done in the stoichiometric subspace of (*ji*)^⊥^, *v*_1_ and *v*_2_ are individually also feasible conversions. Then Eq. (24) for the conservation of energy can likewise be decomposed to 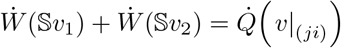.

Because 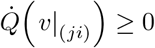, one or both of 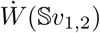 could be positive but they cannot both be negative. If 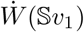 and 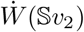 have opposite signs (say 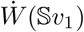 is positive), then Wachtel *et al*. [1] define the *transduction efficiency* of chemical work from *v*_1_ to *v*_2_ through reaction (*ji*) in the background *μ* as

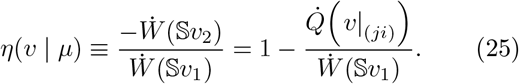

The interpretation of transduction efficiency considered in [1] applies directly to cases in which *v*|_(*ji*)_ is the entire current through reaction (*ji*). In addition to these, we will wish to consider the role of null cycles in redistributing chemical work and dissipation in cases where there may be other flows through the network, such that *v*|_(*ji*)_ is not the entire current through reaction (*ji*), and 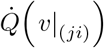 may have either sign. (Negative 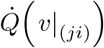 corresponds to a *reduction* in dissipation by null flows that redirect partial flux through pathways parallel to (*ji*), generally allowing for a reduction in overall network dissipation through a Le Chatelier response to external driving.)

###### Feasible conversions using the basis for the null space

Making use of the simplifying assumption dim ker 𝕊|_supp *v*_ = dim_*v*_ (*ji*) = *d* to eliminate extraneous null cycles, we may decompose the work-dissipation identity (22) in terms of a general basis for the null space of supp *v* as

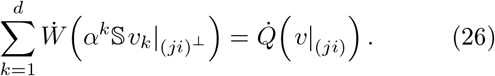

The work term in Eq. (26) for the restriction of each basis element *v*_*k*_ to the complement of (*ji*) in 𝒢, 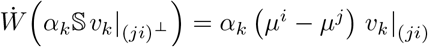.

###### Case *d* = 1: stoichiometric coupling

If *d* = 1, then the decomposition (26) reduces to a single, positive term. This is the case that Wachtel *et al*. [1] term “tight coupling”. As the authors note, for any decomposition into external flows *v*_1_ and *v*_2_ in the manner of Eq. (25), the fluxes in *v*_1_ and *v*_2_ have a fixed relation to *v*|_(*ji*)_ independent of *μ*, because *v*|_(*ji*)_^⊥^ is irreducible (dim ker 𝕊|_supp *v*−(*ji*)_ = 0). Therefore efficiency *η* in this case does not depend on the overall magnitude of *v* but only on the inner products of chemical potentials with parameters from the stoichiometry.

###### Case *d* > 1: non-stoichiometric (potential) coupling

For *d >* 1, in addition to decompositions of the form available when *d* = 1, it is also possible to consider cases in which external *v*_1_ and *v*_2_ are the restrictions of any partial sums of terms in Eq. (26) having respectively positive and negative signs, thus writing

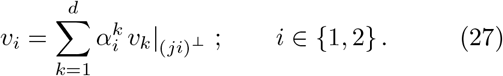

This is the case that Wachtel *et al*. [1] term “loose coupling”. It is transduction by independent null components coupled through the potential drop across a reaction they share. For this case we must have *∂*_𝒢_ *v*_1_ = *∂*_𝒢_*v*_2_ = *∂*_𝒢_ (*ji*), and the projection of their two completions on (*ji*) gives

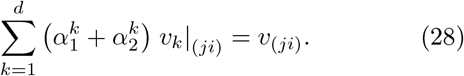

The efficiency expression (25) in this case becomes

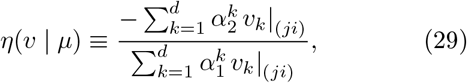

now independent of the chemical potentials given the expansions (27) for *v*_1_ and *v*_2_.

For currents and potentials *μ* solved jointly from the mass-action rate equations, the ratio (29) will vary jointly with potentials that generally are distributed throughout supp *v*, including any dependence on other flows through (*ji*) from external sources in addition to the null component *v*.

**Remark: Complementary regimes of maximum efficiency:** For stoichiometrically coupled reactions at *d* = 1, maximum efficiency (25) is attained in the limit of vanishing kinetic barrier *μ*_*j*_ − *μ*_*i*_ → 0, relative to the potential drops that determine 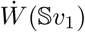. In the alternative case of fully non-stoichiometrically-coupled transduction at *d >* 1, the efficiency (29) is maximized in the limit of arbitrarily high reaction barrier, where *v*_(*ji*)_ → 0 in Eq. (28) and chemical work is transduced “around” the focal reaction from *v*_1_ to *v*_2_.^10^

**Remark: dependence of efficiency on grouping of source and sink currents:** Unlike the simple thermal Carnot cycle [6], coupled through temperature which is a scalar and having a single heat-supplying and a single heat-removing interface between the environment and the system, stoichiometric coupling to the boundary of a reaction permits many divisions of *v*|_(*ji*)_^⊥^ = *v*_1_ + *v*_2_ with opposite signs for work delivered, and corresponding efficiency measures. These choices arise naturally in applications to bioenergetics. Cellular free energy flow is generally seen as being organized around three central “energy buses” [79]: phosphate leaving groups provided in activated form by ATP, electron transfers to and from a variety of redox cofactors, and proton transfer across membranes. A variety of other more specific leaving groups (such as “activated” C_1_ or N_1_ groups carried on dedicated cofactors, or phosphoryl transfer specifically from GTP at the ribosome or uridilation of glucose from UTP) may also serve within reaction subsystems as general-purpose energy carriers [79, 80]. A choice to take the ATP/ADP/AMP couple as a source and to group all other reactions including biosynthesis and redox recharging as the load (as in the examples of [1], yields one efficiency measure, whereas taking the ATP system and the redox cofactor system as two components of the source, independently resupplied by external reactions, and biosynthesis as the load, yields a different measure.

Moreover, for many biologically relevant transductions between redox, phosphate, or proton energy systems, the full transduction occurs through multiple reactions in series, which may exchange buffered species such as protons or water with the solvent. Thus free energy transduced between feasible processes *v*_1_ and *v*_2_ that jointly with *v*|_(*ji*)_ make a null flow may exchange chemical work both by transduction through *v*|_(*ji*)_ and by direct exchange through species in *∂*_𝒢_ (*v*_1_ + *v*_2_) but not in *∂*_𝒢_ (*ji*).^11^

#### 7. Nested ensembles from graphs to subgraphs and flows

In this section and going forward, where we deal with conservative flows in a graph driven by external current boundary conditions, we omit explicit notation for the embedding context that defines feasible processes.

Let *v* then be a conservative flow in a graph 𝒢 with a source current −*J*^ext^ ≡ −*J*(*v*) on *∂*_𝒢_ *v*. Implicitly −*J*^ext^ is a feasible conversion in some larger environment-graph that contains 𝒢 and that we do not write down.

Under the mass-action rate law, with a current boundary source −*J*^ext^ and no other sources or sinks to species in 𝒢, each basis element in ker 𝕊 will add to *v*, redistributing chemical work and dissipation so as to produce a non-equilibrium steady state at a vector of chemical potentials *μ* that minimizes dissipation for the net flow through 𝒢^12^ If we reduce the the dimension of ker 𝕊, by eliminating reactions from 𝒢 while leaving a graph capable of the same net conversion *J*^ext^, the constraints on this minimization problem are thereby increased and the dissipation rate cannot decrease. We wish to study the response of dissipation along the reduction paths from diffusive flows through large graphs to single irreducible flows, which we will term *pathways*.

Therefore consider a series of solutions *v, v*′, *v*^′′^ … to a conversion problem *J*^ext^ = 𝕊*v*^*k*′^, first for a mass-action solution *v* in the full graph 𝒢, then the mass-action solution *v*′ on a subgraph 𝒢′ ⊂ 𝒢, and so on until we reach some irreducible flow *v*_irred_ with supp *v*_irred_ ⊂ … ⊂ 𝒢′ ⊂ 𝒢.

The minimum-dissipation flow *v*′ on 𝒢′ exists also as a flow solution on 𝒢 (where it does not minimize dissipation), and we can obtain *v*′ from the ensemble of events resulting in aggregate in the true minimum-dissipation solution *v* on 𝒢 by biasing that ensemble in a generating function for currents. The relative entropy of the biased from the unbiased distribution, which in a large-number limit becomes the large-deviation function for *v*′ from *v* in 𝒢, allows us to associate a log-(un)likelihood with the constraint arising from removal of the reaction that reduces 𝒢 to 𝒢′, and so on until arriving at supp *v*_irred_.

#### 8. The linear-response approximation for dissipation in detailed-balance systems

Below in Sec. IV E, through relations between the dissipation (23) and a large-deviation function for currents in the linear-response regime, we will show that a natural additive decomposition of dissipation exists for flows that can be written as mass-action solutions on nested graphs. These limits correspond, for stoichiometric networks, to the regime of Ohm’s law for electric circuits [56], and in them we can derive stoichiometric pathway-generalizations of notions such as element-level or circuit-level resistance to through-currents, which are functions of network and pathway topology and of the reference equilibrium, but not of the magnitude of the driving force.

From Eq. (17) for *v*, and the relation *J* = 𝕊*v* on a graph 𝒢, the current for a through-flow is related to the vector of species chemical potentials *μ* = *μ*_º_ + *k*_*B*_*T* log ((*n*) */n*) in the linear-response regime as

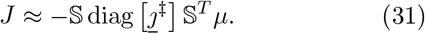

Invertin g Eq. (17), the vector of currents itself is given by diag [1*/J*^‡^] *v* = −𝕊^*T*^ *μ*. The dissipated free energy (23) then becomes

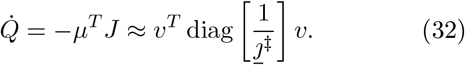

We must generally make case-specific modeling commitments to assign currents *J*^‡^. However, for some applications such as cell physiology, where comparable catalyzed currents must be produced at comparable physiological concentrations and activities for the reaction complexes, it can be reasonable to set *J*^‡^ ≡ 1, in some appropriate rate units characterizing catalysis at the diffusion limit. In such cases we can further reduce the dissipated heat (32) to what we will term the *topological contribution* to dissipation,

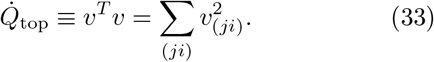

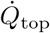 for mass-action rate law solutions can be computed directly from the source current −*J*^ext^ = −*J*(*v*) and the matrix inversion of the kernel 𝕊𝕊^*T*^ from Eq. (31), accounting properly for conserved quantities of the stoichiometry. The representativeness of 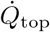 for more physiological realistic cases is studied in App. E.

## III. SAMPLE COMBINATORIALLY-GENERATED TOPOLOGY: SUGAR-PHOSPHATE CHEMISTRY FROM FIVE RULES

Here we illustrate the propagation of constraints and symmetries across a three-level system, from its generating rules to the thermodynamic phenomenology captured in its large-deviation functions, in a biochemical network model that is of independent interest as a putative instance of evolutionary combinatorial optimization. The example is drawn from biological sugar-phosphate chemistry, developed as a graph-grammar model in [2]. The uniform stoichiometry (CH_2_O)_*n*_ of sugars, which requires hydroxyl or carbonyl groups at every carbon center (or bridging oxygen in cyclic forms), leads to a high degree of potential combinatorial complexity in networks generated from very few mechanisms [34, 83].

Sugar chemistry in metabolism is distinctive for both retaining part of this combinatorics and excluding other large parts, suggesting questions about the criteria and complexity of search and optimization problems evidently solved by selection. At the same time, all core biological sugar metabolism is sugar-phosphate metabolism [80], with the positioning and reactions of phosphate groups providing crucial free energy modulation and protection for some carbon centers, so that sugar-phosphate metabolism remains much simpler than unrestricted sugar chemistry [84, 85].

In this section we present the generating rules for a model and a small, computationally-generated reaction network that can be extended to unlimited size by induction on the length of the largest carbon chains. The reaction network contains the two universal biochemical pathways in the Calvin-Benson-Bassham (CBB) cycle [47] and the Pentose-Phosphate Pathway (PPP), which perform the same sugar-phosphate recombination process in reversed orders. We present an exhaustive and ordered enumeration of all integer flows in the network performing this conversion, generated as integer linear programming solutions in the graph-grammar modeling system MØD [13, 16], which grows combinatorially in the network size, but can likewise be systematically extended to arbitrary networks.

We will emphasize the role of discrete *symmetries* respected by the generating rules, and the ways in which these, together with a net conversion such as that of CBB/PPP, constrain the forms of all possible pathways performing the conversion. In particular, the biological Calvin cycle can be shown to be one of two uniquely minimal solutions under these rules, by a direct graphical construction that bypasses a need for exhaustive enumeration.

We also use symmetries to construct a basis for the null space of the stoichiometry in this system, which we show decomposes into a classification of null cycles that remains complete in networks of any size from these rules. The null cycles presented here will be used later in examples of the information-geometric decomposition of large deviation functions and the assignment of costs to reduction sequences from network flows to irreducible pathways as defined in Sec. II D 2, a reduction plausibly performed by natural selection for progressively more substrate-specific enzymes. Finally we will use the null basis elements in examples of both stoichiometric and non-stoichiometric coupling of chemical work that is performed by the process of equilibration to arrive at Onsager’s minimum-dissipation solutions [81, 82] for flow through a network.

### A. System construction from rule declaration and network expansion

#### 1. Five rules

The five reaction mechanisms that support biological sugar-phosphate chemistry are listed in Table I. Each appears in the rule set from [2] as a pair of bidirectional reactions, which are listed in the table together. The list includes one isomerization (aldose-ketose conversion), one condensation (aldolase), one hydrolysis (phosphohydrolase), and two recombining reactions (transaldolase and transketolase) that transfer, respectively, the C_3_ or C_2_ ends from ketose onto aldose sugars.

**TABLE I:** The five rules that generate biological sugar-phosphate conversions, from [2]. The rule name and a brief annotation for its action are shown.

| Name | Description |
| --- | --- |
| Aldolase | Aldol addition / retro-aldol cleavage |
| Aldose-Ketose | Aldehyde-ketone interconversion |
| Phosphohydrolyase | Phosphate hydrolysis |
| Transaldolase | Move C<sub>3</sub> end |
| Transketolase | Move C<sub>2</sub> end |

The rule system is conservative of (CH_2_O) groups and redox-neutral. We note that we have omitted keto-enol tautomerization in the preparation of this example, so carbonyl migration within chains, which would permit a host of aldol cleavages and recombinations at sites other than the terminal carbon atoms, resulting in branched-chain as well as linear molecules, are not included in the model. (Their effects may be seen in [34].)

#### 2. A biologically motivated throughput problem

Biological sugar-phosphate metabolism is found in the core of both anabolic and catabolic chemistry. Anabolically, it is the subnetwork in the Calvin cycle that prepares ribulose-5-phosphate to receive CO_2_ driven by hydrolysis of the main backbone in the enzyme RuBisCO, forming two molecules of 3-phosphoglycerate, which are reduced to glyceraldehyde-3-phosphate to initiate two repeats of the pathway, rendering it autocatalytic. In anabolism, ribulose-5-phosphate is re-arranged to glyceraldehyde-3-phosphate to enter one of the glycolytic pathways of energy metabolism.

The common feature of both pathways is the lossless rearrangement of (CH_2_O) groups between chains of two lengths (3 and 5) that have no common divisor. As attested in the fact that both are ancient, very widespread, and near-reverses of each other, they can also operate near equilibrium at physiological substrate concentrations [52] except in the steps of phosphorylation or phosphate hydrolysis (from different cofactors and kinetically controlled), a property that is important for the thermodynamic efficiency of the Calvin cycle with the unusual mechanism of action of RuBisCO [86], and for the yield of chemical work in the Pentose-Phosphate Pathway.

The reaction schema that defines the universal conversion problem for the PPP and the CBB cycle is

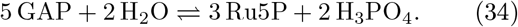

The forward direction characterizes CBB, and the reverse direction PPP.

A joint program of network expansion with molecule synthesis was carried out using the graph-grammar system MØD with the rules in Table I, and glyceraldehyde-3-phosphate and water as starting compounds. Labels for the chemical species up to length C_8_ are shown in Fig. 1 as they will be arranged in graphs of the reaction network in this paper. In addition to water and orthophosphate, sugar compounds appear as aldose-monophosphates, ketose-monophosphates, and bisphosphates. Their physiological data from eQuilibrator 3.0 [62, 87] are given in Table II in App. E 1.

**FIG. 1:**
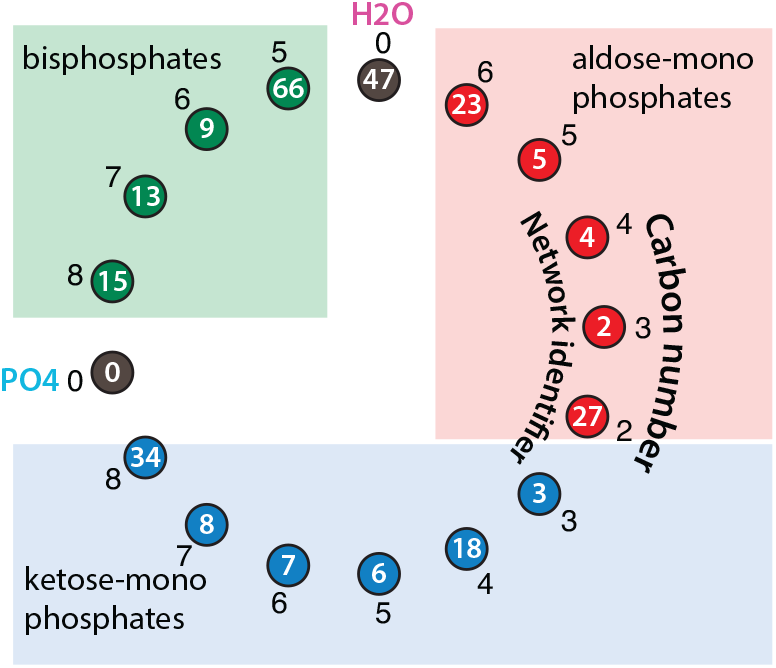
The diagram layout to be used. Filled circles are species in the hypergraph notation following Feinberg [20], in which white letters are the internal molecule identifier produced by MØD. Aldose and ketose monophosphates, and bis-phosphates, are arranged in the series shown, and the carbon number of the sugar is indicated in the outer ring.

The stoichiometric graph of all reactions among these compounds is shown in Fig. 2. This presentation of hypergraphs uses a doubly-bipartite representation in which each graphical element corresponds to an element in the analysis of [19, 20], and to a term in the rate equation. Filled circles are species, open circles are complexes; heavy lines are reaction edges, and thin lines show the stoichiometry of the complexes by connecting each complex to the species it comprises. Directionality of the reaction edges is indicated, in this case, by assigning different colors (green and red) respectively to the input and output complex nodes. For this network expansion, 17 chemical species and 28 unidirectional reactions appear in solutions to the conversion (34).

**FIG. 2:**
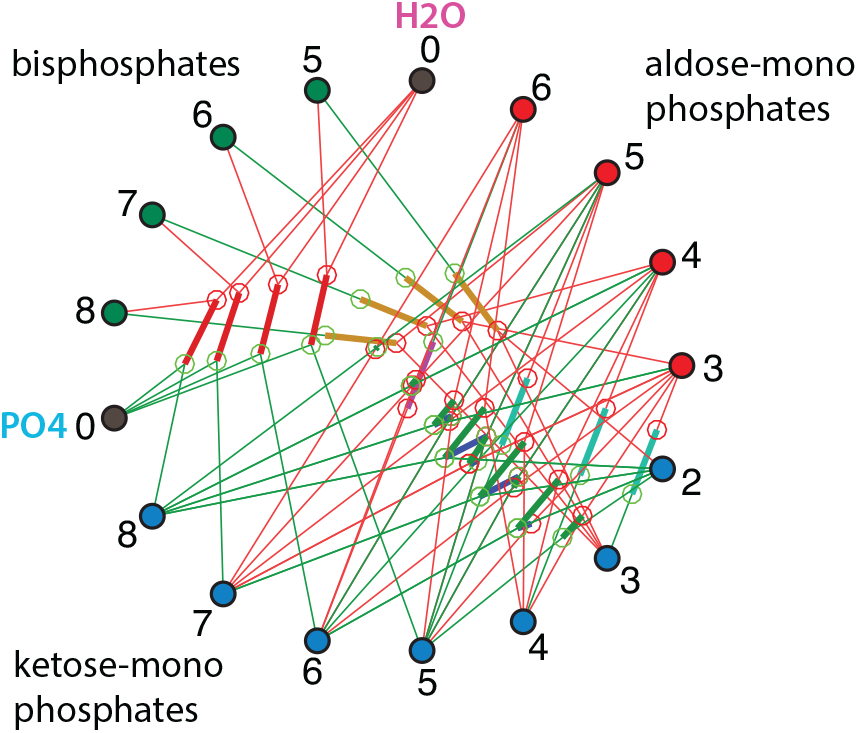
All edges generated for these compounds by the rules in Table I. Identifier numbers are omitted from the nodes to improve clarity.

**FIG. 3:**
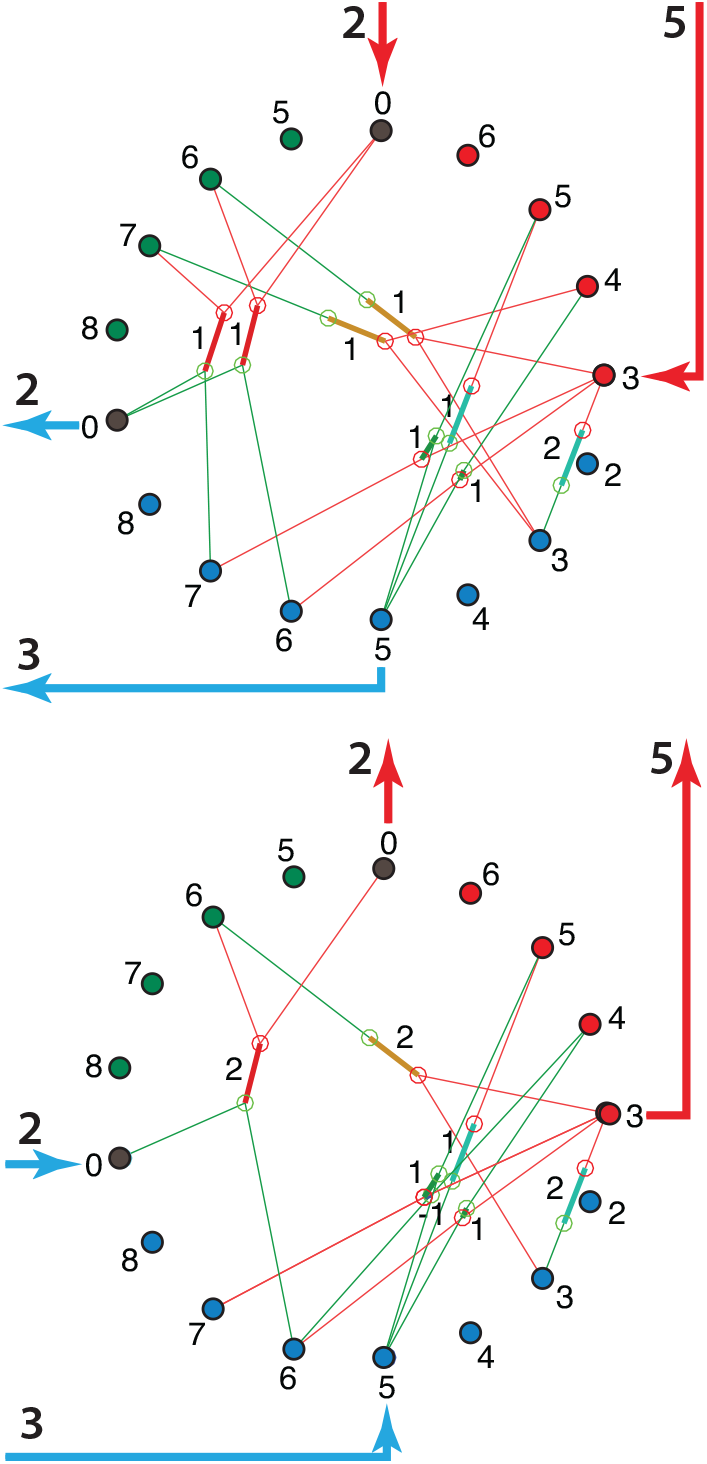
Top panel: the sugar re-arrangement part of the Calvin-Benson cycle as an integer-flow solution *v* with net currents *J*_ext_ = −𝕊*v* from the environment to the network. Bottom panel: one version of the canonical Pentose-Phosphate Pathway as an integer flow solution with the equal and opposite *J*_ext_.

#### 3. The instantiations of rules in literal reactions, and rule composition

As a first step in deriving the underlying patterns that structure the reaction network in Fig. 2, we separate the literal reactions according the the rule that generates each. App. A shows reactions from Fig. 2 grouped by generating rule, in a series of separate figures.

Because the aldehyde groups on which aldol addition acts are created only at the ends of sugars (in the absence of carbonyl migration), all the compounds in Fig. 1 are simple linear sugar-phosphates with a single aldose or ketose end-group, or else are phosphorylated at both ends. Moreover, the rules in this model do not distinguish stereochemistry, meaning that stereoisomers are treated collectively in each node. These simplifications make it possible to refer to all species with an index notation, by symbols *A*_*n*_, *K*_*n*_, or *B*_*n*_, respectively for aldose, ketose, or bis-phosphorylated sugars of *n* carbons. Table II shows the action of the five rules in that variable notation.

**TABLE II:** Rule symbols paired with the conversions the rules make on 1-complexes or 2-complexes of sugar-monophosphates. Legend: AlKe (aldose-ketose); KeAl (ketose-aldose); TAL (transaldolase); TKL (transketolase); AL (aldolase); PHL (phosphohydrolase). In the fifth line, TAL º TKL indicates function composition: the output of a TKL reaction is input to the TAL reaction. In the sixth line, AlKe⊕KeAl indicates parallel application, AlKe on input *A*_*n*_ and KeAl on input *K*_*n*+1_.

| reaction | schema |
| --- | --- |
| AlKe/KeAl | $A_n \rightleftharpoons K_n$ |
| TAL | $A_n + K_m \rightleftharpoons K_{n+3} + A_{m-3}$ |
| TKL | $A_n + K_m \rightleftharpoons K_{n+2} + A_{m-2}$ |
| AL | $A_n + K_3 \rightleftharpoons B_{n+3}$ |
| PHL | $B_n + H_2O \rightleftharpoons K_n + H_3PO_4$ |
| TAL $\circ$ TKL | $A_{n+1} + K_n \rightleftharpoons A_n + K_{n+1}$ |
| AlKe $\oplus$ KeAl | $A_n + K_{n+1} \rightleftharpoons K_n + A_{n+1}$ |

In addition to the elementary rules, which were listed in Table I, the last two lines of Table II show two composite reactions, with º representing function composition acting to the right, and ⊕ representing parallel action of the two rules on independent substrates. It is seen that serial application of transketolase followed by transaldolase has the same effect as two aldose/ketose isomerizations acting on backbones differing by length 1. This equivalence will be important later in the analysis of null cycles.

#### 4. Prelude and fugue structure of all solutions to the conversion problem entailed by the rule algebra

Distinct integer solutions *v* to *J* = 𝕊*v* for *J* corresponding to schema (34) can easily number many hundreds even on small graphs; for the graph of Fig. 2 we enumerated 744 distinct *supporting graphs*, and for each reducible flow the supporting graph will admit a tower of other solutions differing by integer multiples of null flows (details in App. C). Yet this nominal combinatorics can be can be quickly systematized because the combination of constraints from phosphate number, aldose number, and backbone length on the application of these rules imposes certain common architectures on all solutions.

The first architecture we note is a structure we term preludes and fugues. All aldol condensations must be followed by dephosphorylations to produce ketose monophosphates, and a minimum of two molecules of dihydroxyacetone phosphate (DHAP) are required to produce any bisphosphate from the glyceraldehyde-3 phosphate (GAP) inputs. Therefore all solutions possess (up to addition of null cycles) a composite reaction that we term a *prelude* of the form 2 AlKe ⊕ PHL º AL, where AlKe is the triose-isomerase (TIM) reaction *A*_3_ ⇌ *K*_3_ between GAP and DHAP. The aldol condensation may act on any aldose sugar, so the preludes as a class have the schema

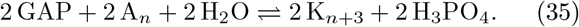

Four “pure” preludes are shown in Fig. 19 of App. B 1, which are instances of the schema (35) for a single A_*n*_ with *n* ranging from 2 to 5. Six mixed preludes also arise, which are sums (⊕) of 1/2 of two instances of the schema (35) with two different aldoses of length *n* and *m*. These are the only possibilities.

If, from the overall reaction schema (34) we extract the prelude schema (35), the remainder is a combination of TAL, TKL, and AlKe/KeAl edges that we term a *fugue* for its typically cyclic form, with the schema^13^

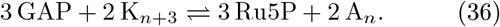

(For mixed preludes, corresponding mixed fugues are the complements.)

The remaining diversity of integer flow solutions results from adding integer multiples of null flows to any instance of this prelude-fugue backbone. Some null cycles are supported entirely within fugues, while others may span preludes and fugues. We turn now to a decomposition of the null space making use of rule symmetries.

### B. The null spaces

#### 1. A basis for null flows grouped by rule type

The rank of the null space for a reaction network such as Fig. 2 can of course be computed directly from the number of reactions and the stoichiometric dimension *s*. Rather than choose an arbitrary basis for the null space by applying a generic row-reduction algorithm to the stoichiometric matrix S, however, we use the organization imposed on the network by the symmetries and conservation laws entailed in the rules of Table I to construct four classes of null cycles that follow shared templates, which we show later are a complete basis in networks extended to arbitrary maximal *n*.

##### a. Null cycles from prelude differences

The difference of two preludes (35) beginning with *A*_*n*_ and *A*_*m*_, is twice the TAL reaction (see Table II)

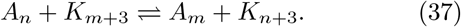

Therefore subtracting the TAL edge (37) from 1/2 the prelude differences results in a null cycle. Fig. 20 in App. B 1 shows graphs of a complete basis for null cycles of this form in the network.

##### b. Null cycles from TAL reactions

More generally, for three distinct integers *m, n*, and *p*, and a triple (*m*′, *n*′, *p*′) a cyclic permutation of (*m, n, p*),^14^ parallel action (⊕) of three TAL reactions in the relations

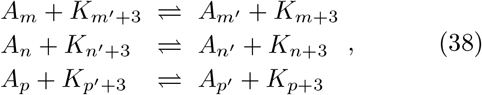

is null. We note from the fact that the ketose set in the schema (38) is the +C_3_ image of its aldose set, that the total carbon count in any such schema is odd.

Cycles of the form (38) will be called **trefoils**. Their properties are elaborated more fully in Fig. 24 of App. B 2 a. The trefoils are minimal instances of what we will call more generally **braided cycles**, because they perform a bucket-brigade style transport of a C_3_ or C_2_ end-group along an unoriented surface that makes a non-trivial cycle because it has the gluing rules of one or more Möbius twists (illustrated in Fig. 7 of Sec. III C). Fig. 21 in App. B 1 shows a basis for all linearly independent trefoils TAL trefoils in this network.

##### c. The TAL reaction with a zero-carbon aldose

The class of TAL trefoils can be unified with the previous null cycles from preludes by making a re-arrangement that is not meaningful as a chemical schema but that respects symmetry and conservation under the rules. If we formally identify phosphate groups net of the water they would release under dehydrating phosphorylation as “zero-length aldose-phosphates”,^15^

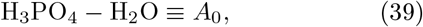

the rule composition PHLºAL in Table II follows the formula for a TAL reaction, with *m* = 3. The the null cycle from prelude differences then fits the schema (38) for a TAL trefoil with *p* = 0 and permutation (*m*′, *n*′, *p*′) = (*p, m, n*).

##### d. Null cycles from TKL reactions

The trefoil corresponding to schema (38) but for TKL reactions is

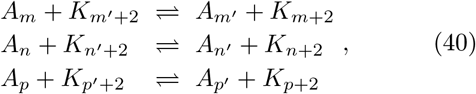

From the fact that the ketose set in a TKL trefoil is the +C_2_ image of its aldose set, it follows that the total carbon count in a TKL trefoil is even. Fig. 22 in App. B 1 shows a basis for all linearly independent TKL trefoils in this network.

##### e. Null cycles from opposed AlKe and KeAl reactions

The direct sum of the fifth and sixth lines of Table II is a null cycle that effectively substitutes an AlKe conversion on one carbon chain for a parallel conversion on the chain one carbon longer. Fig. 23 in App. B 1 shows the two linearly independent cycles of this pattern supported in this network.

#### 2. The rule-derived basis elements span the null space in a network continued to any maximum size

App. B 2 shows that the set of null cycles constructed here by rule groups are in fact a basis for the entire null space of networks of the form in Fig. 2, continued inductively to any maximal chain length *n*.

### C. Integer-flow solutions and their supporting graphs

#### a. List of flows and integer properties

An integer linear program (ILP) solver was used to enumerate the first 763 solutions 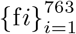, to *J* = 𝕊fi for *J* the net conversion (34). The solutions are exhaustively enumerated in increasing order of the number of reactions used (equal to the number of edges in the supporting graph), which we designate #_reactions_, and within each #_reactions_, in increasing order of the sum of (absolute) magnitudes of the currents through all reactions. App. C 1 summarizes combinatorial measures of solutions generated by the ILP solver, including the number of distinct integer solutions fi, the number of distinct supporting graphs supp fi, and the counts of reducible and irreducible flows by the definitions of Sec. II D 2. In plots of these solutions, we will label reaction edges with the respective current components of fi, with signs relative to the directionality of the edge used.

The sugar re-arrangement part of the CBB cycle was obtained as flow solution f13, and the canonical PPP was obtained as f1. These two solutions are shown in Fig. 3, accompanied by external source currents −*J*^ext^ = −𝕊f*i* that balance internal conversion in steady states. As a first step in identifying what properties of a flow might have served as criteria for the selection that have made these two solutions biological universals, we plot in Fig. 4 both #_reactions_ as a measure of the requirements a supporting graph places on genomic encoding of enzymes, and what we have termed the “topological” contribution to dissipation in the linear response regime, 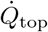 from Eq. (33) for the 763 stoichiometrically unique flow solutions.^16^

**FIG. 4:**
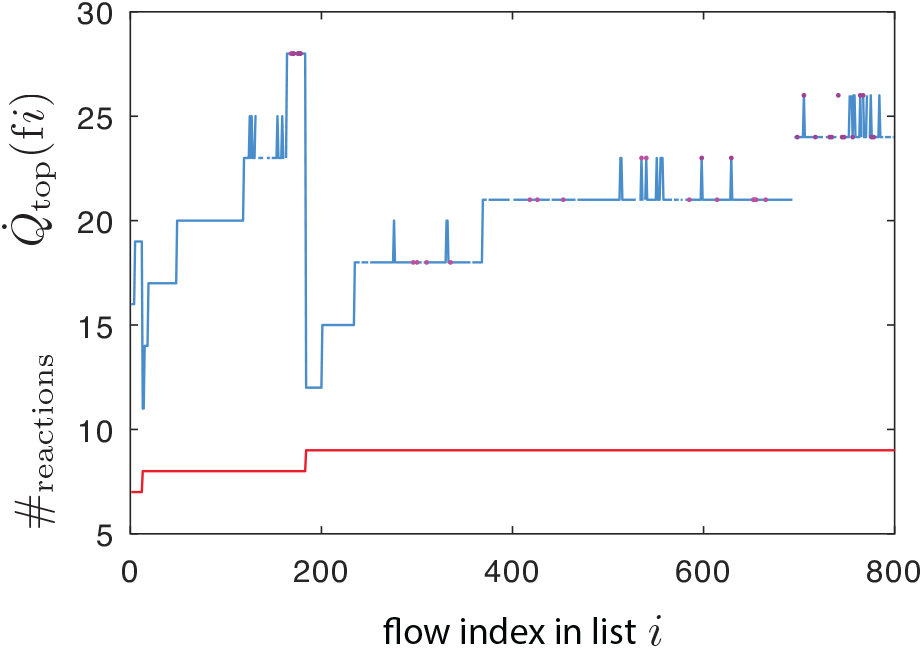
763 solutions fi to the throughput problem, showing the number of reactions in the supporting graph supp fi (red) and the topological contribution to dissipation 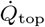 (blue and symbols) from Eq. (33) evaluated for fi. 19 integer flows (light magenta dots) produce a graph supp fi that appears again as the supporting graph of a second solution (dark magenta dots).

#### b. Flow irreducibility and nested supporting graphs

The list of integer flow solutions affords us opportunities to illustrate the nested-subgraph hierarchy for flow reduction from Sec. II D 7 and the utility of the basis of null flows from Sec. III B in understanding that decomposition, as well as examples of stoichiometric and non-stoichiometric transduction of chemical work during the relaxation to equilibrium on different graphs. To keep the presentation compact, we will illustrate subgraphs with 2- and 3-dimensional null spaces from the start, out of which several examples of flow reduction and work transduction will then be extracted.

Let 𝒢 stand for the whole network in Fig. 2. We begin by considering solution f14, shown in Fig. 5, an irreducible flow supported on 8 reactions, which shares uniquely with f13 the property that it has the minimal value 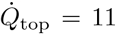 in the plot of Fig. 4. Brief inspection shows that f14 is an image of f13 transposed downward in chain length of the bisphosphates by one carbon, a homology to which we return in Sec. III D.

**FIG. 5:**
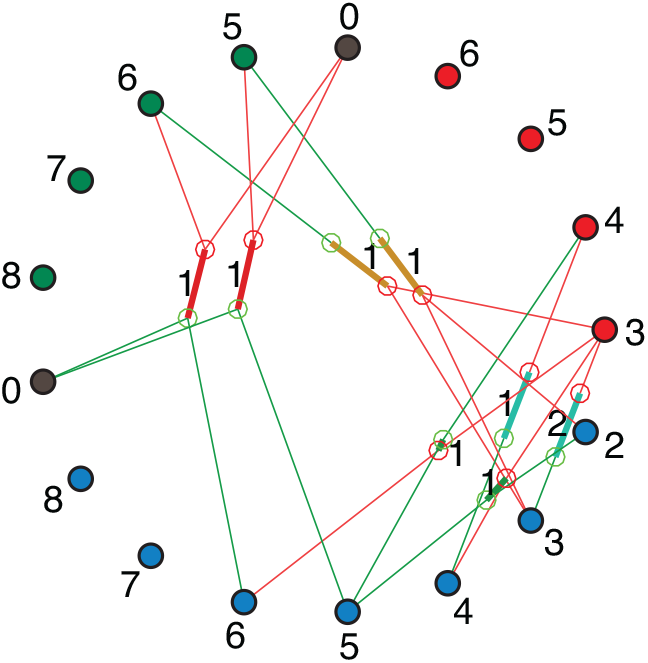
Flow 14, corresponding to the first panel in the lattice graph of Fig. 10 below.

Table III lists the integer current counts fi for seven integer flows in the ILP list including f14. Edges are designated e⟨edge-num⟩ where ⟨edge-num⟩ is an index assigned by the ILP solver. A common mixed prelude (35) acting on one copy of A_2_ (glycolaldehyde phosphate, GLP) and one copy of A_3_ (glyceraldehyde-3 phosphate, GAP) constitutes the first five rows in the table.

**TABLE III:** Count vectors on seven flows in the ILP list supported on the graphs of Fig. 6 and Fig. 7. First five rows are the prelude, not shown in Fig. 7, shared by all flows. e5 is the TIM reaction converting GAP to DHAP. e1 and e69 convert GAP+DHAP+H2O to F6P+H3PO4. e40 and e182 convert GLP+DHAP+H2O to Ru5P+H3PO4. The triple e22, e98, e96 are the 234 TKL trefoil. The triple e96, e23, e97 are the 235 TKL trefoil. The quadruple e6, e7, e15, e97 are the AlKe null cycle.

| edge | f629 | f18 | f14 | f15 | f535 | f193 | f184 |
| --- | --- | --- | --- | --- | --- | --- | --- |
| e5 | 2 | 2 | 2 | 2 | 2 | 2 | 2 |
| e1 | 1 | 1 | 1 | 1 | 1 | 1 | 1 |
| e69 | 1 | 1 | 1 | 1 | 1 | 1 | 1 |
| e40 | 1 | 1 | 1 | 1 | 1 | 1 | 1 |
| e182 | 1 | 1 | 1 | 1 | 1 | 1 | 1 |
| e22 | -1 | 0 | 1 | 2 | 3 | 1 | 1 |
| e98 | -2 | -1 | 0 | 1 | 2 | 0 | 0 |
| e96 | 3 | 2 | 1 | 0 | -1 | 0 | 0 |
| e23 | 0 | 0 | 0 | 0 | 0 | 1 | 1 |
| e97 | 0 | 0 | 0 | 0 | 0 | 1 | 0 |
| e6 | 1 | 1 | 1 | 1 | 1 | 1 | 0 |
| e7 | 0 | 0 | 0 | 0 | 0 | 0 | 1 |
| e15 | 0 | 0 | 0 | 0 | 0 | 0 | 1 |
| $\dot{Q}_{\text{top}}$ | 23 | 14 | 11 | 14 | 23 | 12 | 12 |

It is immediately apparent that five flows (f629, f18, f14, f15, f535) form a series in which 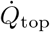 depends parabolically on the offset in the list from f14 which is the minimizer. The flows differ by integer multiples of a TKL trefoil on edges e22, e98, e96, shown in Fig. 22 of App B 1, which lies in the supporting graph of the “fugue” conversion (36). The three backbone lengths of this trefoil are (*m, n, p*) = (2, 3, 4), so we will refer to it (uniquely) as the “234” TKL trefoil. Flows f18, f14, and f15 are all irreducible. The union of the supporting graphs of any two of them yields a subgraph 𝒢′ ⊂ 𝒢 that is the supporting graph of reducible flows f629 and f535 and has null dimension 1. Reduction of 𝒢′ to any of the three irreducible flows is performed by adding the appropriate integer multiple of the trefoil to cancel one edge in its support.

We extend the previous example by taking the union of the supporting graphs of f535 and f193 in Table III to form the subgraph shown in Fig. 6. This graph has null dimension 2. In addition to the 234 TKL trefoil, it supports a 235 TKL trefoil through edges e23 and e97, non-stoichiometrically coupled to the 234 TKL trefoil through e96. All four of f18, f14, f15, and f193 are irreducible, and are obtained by canceling edges through one or both trefoils.

**FIG. 6:**
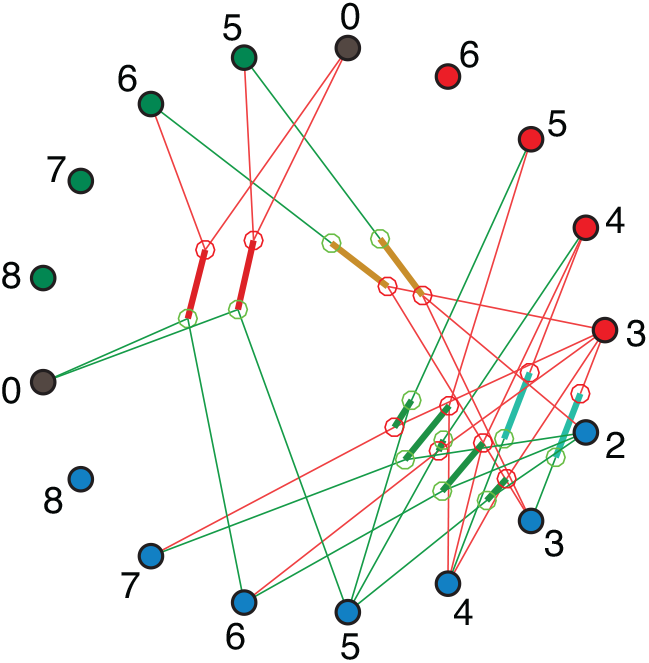
A network formed from the union of the supporting graphs for three irreducible flows f14, f18 and and f193. A subgraph with 8 edges hosts f14 uniquely. It is the first lattice diagram in Fig. 9 below. The union of that graph with e98 (the lattice edge from (*A*_2_, *K*_6_) to (*A*_4_, *K*_4_,)) hosts f535 and f629 with one additional TKL trefoil of backbones 234. The further union with e23 (the lattice edge from (*A*_3_, *K*_7_) to (*A*_5_, *K*_5_)) and e97 (the lattice edge from (*A*_2_, *K*_7_) to (*A*_5_, *K*_4_)) adds a second TKL trefoil with backbones 235 and common edge e96 with the first trefoil. Two TIM reactions and single firings of each AL/PHL sequence are fixed by the topology for the conversion (34), so the prelude on this graph is independent of the background. The 5 TKL edges and the remaining C_4_ AlKe edge constitute the fugue. Like the preludes, the AlKe reaction is topologically constrained to fire one time. The only two degrees of freedom responsive to the kinetics are the circulations in the two trefoils, illustrated below in Fig. 14.

**FIG. 7:**
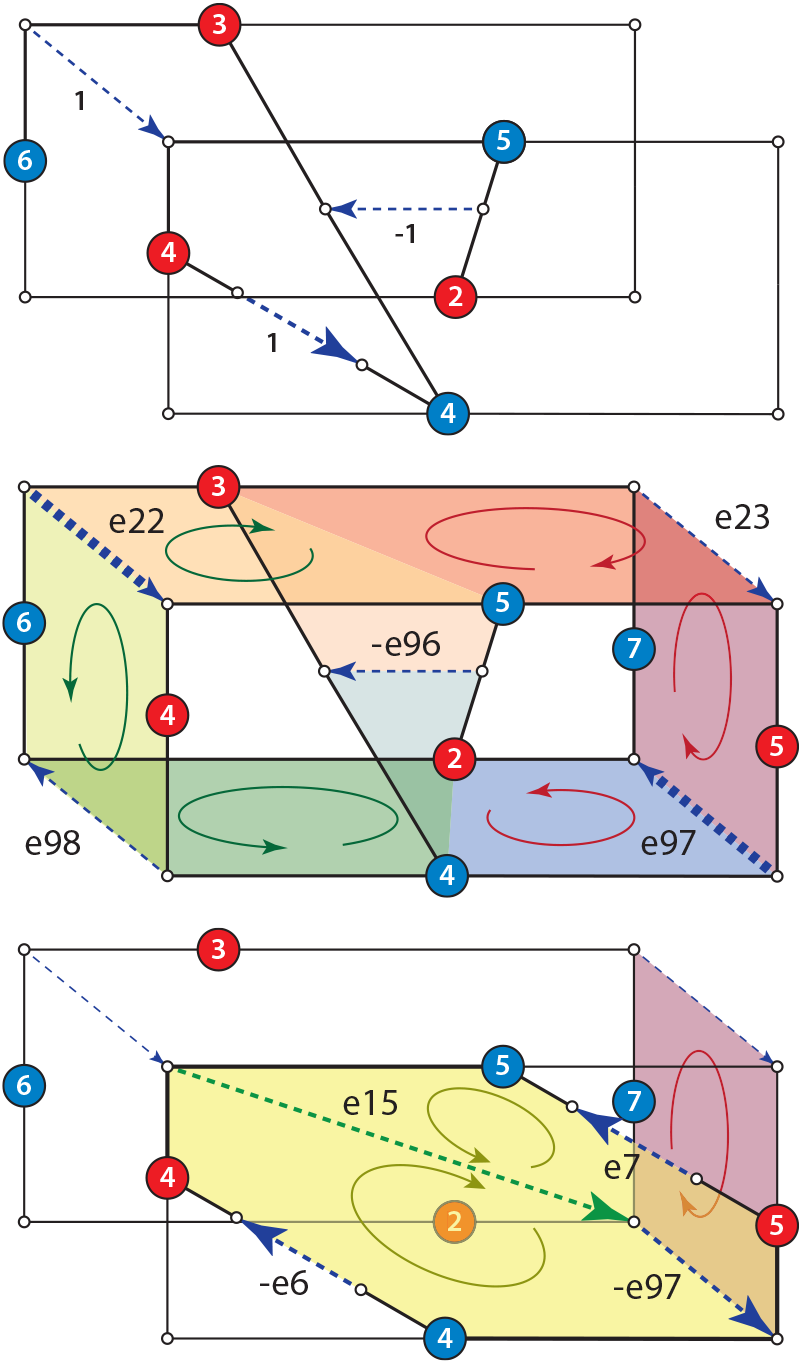
Top panel shows the three reactions in f14 that are not part of the prelude creating F6P from GAP + DHAP and Ru5P from GLP and DHAP. Middle panel expands the set of species and reactions to correspond to the supporting graph in Fig. 6. A TKL trefoil with aldose backbone lengths 2, 3, 4 covers the faces on the left-hand side of the cube, shown with green circulation arrows. A second TKL trefoil with aldose backbone lengths 2, 3, 5 covers the faces on the right-hand side of the cube, shown with red circulation arrows. The two trefoils are non-stoichiometrically coupled through any potential drop that arises on e96. Arrows are shown in the directions that make trefoils null. The sense of edges in the MØD listing, relative to the drawn arrows, is indicated with ± signs. All currents and potentials will be similarly signed relative to drawn arrow directions. Bottom panel shows the AlKe null cycle of Fig. 8 overlain on the 235 TKL trefoil, with which it shares edge 97. The pair of dark-gold circulations shows the stoichiometrically coupled reaction directions.

The nature of null cycles and their connection in reducible flows can be further clarified by suppressing nodes and reactions from the prelude in the full network and arranging the remaining fugue subgraph spatially to express patterns of symmetry in the network. We do this in Fig. 7 for the subgraph formed from the union of supporting graphs for f535, f193, and f184 from Table III.

To set out what functions as a minimal fugue, the edges and their integer currents in f14 are shown as the top panel in the figure. The second panel shows the rearranged graph from Fig. 6, with the 234 trefoil on the left-hand side and the 235 trefoil on the right-hand side of a bisected rectangular solid. Differently-colored facets on each surface have perimeters around which carbon backbones cycle in the two TKL trefoils, as the C_2_ ketose end of the sugar is passed around three of them in sequence. The “braided” nature of the trefoil, expressed by crossing lines in Fig. 24, is shown here by the way these facets must be joined at their edges to form a Möbius band to close the cycle.

The third panel in the figure shows, separately, the additional face and a new AlKe null cycle brought in with the inclusion of the supporting graph for f184. The AlKe null cycle is one of the two shown as the first panel of Fig. 23 in App B 1. In Table III, an integer multiple of this flow replaces the e6 AlKe isomerization and e97 TKL recombination of f193 with the e7 isomerization and the e15 TAL recombination of f184.

#### c. Stoichiometric versus potential coupling

In Sec. IV D below, after we have introduced flow solutions under the mass-action rate law (and the thermodynamic landscape needed to compute them), we will consider cases in which the null cycles of Fig. 7 appear as additions to a backbone such as f14 with non-integer coefficients. In any solution *v* that activates the 234 and 235 trefoils, dim_*v*_(e96) = 2, and chemical work will be transduced non-stoichiometrically among the two aldose/ketose pairs in *∂*_𝒢_ (e96).

For contrast, The AlKe null cycle (shaded yellow) in the third panel of Fig. 7 is shown in isolation in Fig. 8. Two different decompositions of this graph into environment and coupling-graph components illustrate the distinction between feasible environment transitions that either are, or are not, stoichiometrically coupled themselves.^17^ If the edges e6 and e7 are regarded as environmental input and output, the coupling graph is the link-age class e15 ∪e97. For this model, the input and output have no direct stoichiometric connection. Alternatively, e6 ∪e7 can be used as the coupling graph, with e97 as an environmental input, and e96 as an environmental output. For this example, which we develop quantitatively in Sec. IV D 2, the input and output are coupled by the complex (*A*_2_, *K*_7_), and chemical work is transduced between two other complexes relative to it.

**FIG. 8:**
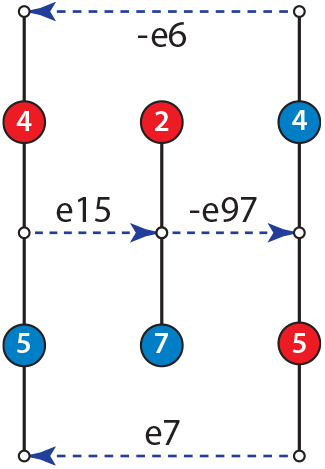
Example of stoichiometric coupling between two AlKe edges in a null flow. If the AlKe edges are regarded as the environmental input and output, then coupling is through the TKL º TAL linkage class in the center. If the pair of AlKe edges are taken as the coupling graph, then the TKL edge may be the environmental input and the TAL edge its output, the case illustrated below in Fig. 15.

### D. Deriving constraints on flows from rule properties using lattice representations of reactions and flows

So far the sequence we have exhibited in rule-based modeling has used rules to generate networks, and then used enumeration to derive bases for the Kirchhoff flow decompositions on the networks, without further direct appeal to the rules. We may also use properties of rules to directly answer questions about flow solutions without the intermediary step of enumeration.

In this section as an example we will derive the minimum value 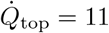 in Fig. 4 and the reason exactly two flow solutions are possible from this rule set that achieve that value. Along the way we will show that several network properties, such as the number and sizes of linkage classes in the complex network, and the deficiency, can be computed from simple graphical arguments.

The rule properties we will use are discrete symmetries and associated conservation laws, which propagate upward from the rule level through graph generation to emerge as constraints on flow solutions. We will introduce a lattice representation for reactions as an alternative to the hypergraph, on which solutions to the conversion problem *J* = 𝕊*v* become simple closed curves on the lattice in place of stoichiometric flows.

#### 1. Symmetries and conserved quantities of rules

##### a. Reflection symmetries of TAL, TKL, and AlKe reactions

Table IV shows how each of four rules or rule compositions induces a mapping of a certain function of the carbon numbers of aldoses and ketoses between its input and output complexes. The schema for all of the transformations has the form *A*_*n*_ + *K*_*m*_ ⇌ *A*_*n*_′ + *K*_*m*_′. If *m* = 0 the reaction acts on 1-species complexes. Reflection symmetry implies conservation: the first two lines in Table IV conserve |*n* − *m* + 3| = |*n*′ − *m*′ + 3|, the third line conserves |*n* − *m* + 2| = |*n*′ − *m*′ + 2|, and the final line conserves |*n* − *m*| = |*n*′ − *m*′|

**TABLE IV:**
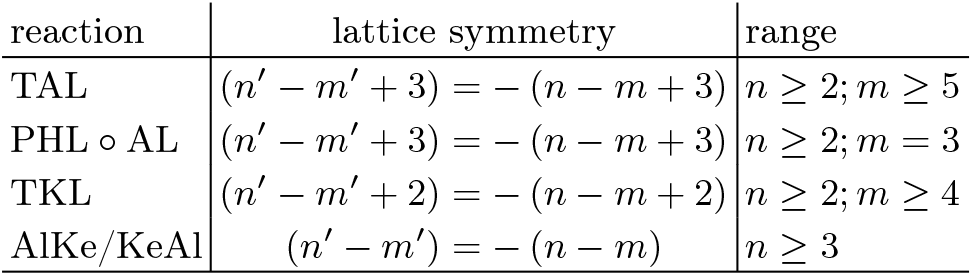
Four reactions with schemata in the form *A*_*n*_ + *K*_*m*_ ⇌ *A*_*n*_′ +*K*_*m*_′, where *m* = 0 indicates a 1-species complex. Reflection symmetries in the lattice of 1-complexes and 2-complexes map to coordinates (*n, m*) on the abscissa and the ordinate respectively, in Fig. 9.

##### b. Lattice representation of reactions

Each of the reactions in Table IV acts between either 1-complexes or 2-complexes, where the 2-complexes contain one aldose and one ketose. We may plot any complex (*A*_*n*_, *K*_*m*_) in an integer lattice at coordinates (*n, m*). 1-complexes occupy the axes with either *n* or *m* equal to zero. By the identification (39) of orthophosphate net of its eliminated water as the “zero-carbon” aldose-phosphate *A*_0_, the output of the rule composition PHL º AL, a ketose monophosphate with eliminated orthophosphate, may be regarded interchangeably as a 1-complex or the 2-complex with *A*_0_. The symmetry maps of Table IV become links connecting the input and output complexes of the reaction, as shown in Fig. 9.

**FIG. 9:**
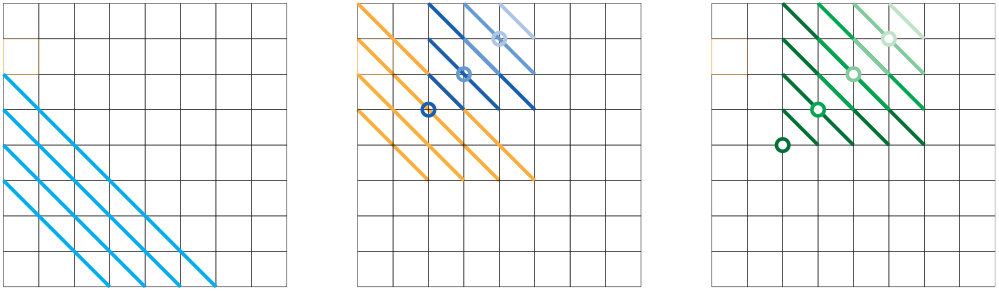
Lattice diagrams for all edges (plus the composite PHL ° AL) in null flows. Left panel: AlKe edges (cyan). Center panel: TAL edges (blues) and PHL ° AL (orange). Right panel, TKL edges (greens). To aid visibility for edges that overlap, TAL and TKL edges are grouped into “tiers” anchored at fixed points (circles), with darkness distinguishing between tiers. TAL plus PHL ° AL and TKL both have 10 distinct edges.

All links have slope -1 because carbon number is conserved (a conservation law on the full graph). The figure shows that the symmetry axis of AlKe/KeAl reactions has as intercept the origin; that for TAL and PHL ° AL links has intercept (0, 3), and that for TKL links has intercept (0, 2).

**Remark:** The TAL and TKL reactions are reflections in Fig. 9, meaning that their compositions TAL2 = TKL2 = *I*, the identity on complexes (as transformations, they are nilpotent). The equivalent inversion on 1-complexes is between unidirectional reactions KeAl°AlKe = AlKe° KeAl = *I*.

#### Aldose-minus-ketose count

In addition to the symmetries acting on differences of carbon number in aldoses and ketoses, the rules in Table IV act on the difference of aldose and ketose counts. Since TAL, TKL, and PHL ° AL have both input and output 2-complexes with one aldose and one ketose, the difference count is zero and is invariant. Only AlKe changes aldose-minus-ketose count to its opposite.

**Lemma:** It follows that no flow omitting AlKe edges can accomplish the net conversion (34), which may be written using the identification (39) as 5 *A*3 ⇌ 2 *A*0 + 3 *K*5, and so requires a net AlKe conversion of 3.

### 2. The complex network and deficiency

The complex network visible in Fig. 2 consists only of chains. All linkage classes in the AL, PHL, AlKe/KeAl reactions are singletons. The only linkage classes with more internal structure are those involving TAL and TKL edges, and their cardinality and structure can be understood using the lattice maps of Fig. 9.

Extended linkage classes arise only where multiple reaction types share a complex, and this occurs only within the lattice region (*An, Km*) with 2 ≤ *n* ≤ 5 and 5 ≤ *m* ≤ 8 spanned by TAL edges in the middle panel of Fig. 9. The net reaction from composition of the two edge types in either order is listed in Table V.

**TABLE V:** The C1 ratchet resulting from composition of TAL and TKL edges. Serial application of these compounds traces the chain-form linkage classes from Fig. 2 in reverse orders.

| reaction | schema |
| --- | --- |
| $\text{TAL} \circ \text{TKL}$ | $A_n + K_m \rightleftharpoons A_{n-1} + K_{m+1}$ |
| $\text{TKL} \circ \text{TAL}$ | $A_n + K_m \rightleftharpoons A_{n+1} + K_{m-1}$ |

Because carbon number is conserved, both the TAL and TKL edges that share common complexes form networks only within single lines. So the number of linkage classes in Fig. 2 equals the number of diagonals in the region spanned by TAL edges (including the fixed points at corners), which is 2 × 4 − 1 = 7.

Moreover, within each linkage class, because TKL edges cover complexes outside those spanned by TAL edges, we can associate, recursively, each lattice point within the blue region 2 ≤ *n* ≤ 5, 5 ≤ *m* ≤ 8 as the origin of a unique edge, alternating TKL and TAL. Thus the number of lattice points (4 × 4 = 16) saturates the number of TKL and TAL edges (5-choose-2 + 4-choose-2 = 10 + 6 = 16). The lengths of the linkage classes, proceeding from smallest to largest total carbon number, are 1, 2, 3, 4, 3, 2, 1. Because the composition of Table V is a ratcheting transfer of C1 groups, all linkage classes are chains.

**Corollary:** The complex network has no cycles, so the right null space of *Y* 𝔸 has no flows in ker 𝔸, the adjacency matrix from Eq. (5). Therefore the deficiency *δ* [20] equals dim ker (*Y* 𝔸), the dimension of the null space.

#### 3. Mapping hyperflows to simple cycles in lattice graphs

By adding three classes of elementary links to the maps between complexes from Fig. 9, we may also represent flows with lattice maps, where they appear as simple closed loops (or collections of loops). The three new classes of links, which connect 1-complexes and 2-complexes to represent inputs or outputs of species to the network, are listed in Table VI.

**TABLE VI:** Three “reactions” that interconvert 1-complexes and 2-complexes through the input or output of species in the net conversion schema (34), and to simplify later plots, the species DHAP which is always produced from GAP by AlKe at flux 2.

| reaction | lattice map |
| --- | --- |
| $\text{GAP\_in}$ | $(K_n) \rightarrow (A_3, K_n)$ |
| $\text{DHAP\_in}$ | $(A_n) \rightarrow (A_n, K_3)$ |
| $\text{Ru5P\_out}$ | $(A_n K_5) \rightarrow (A_n)$ |

Unlike the links representing reactions in Table IV, the addition and removal links in Table VI are operations we perform on the lattice, assembling or disassembling complexes, and so they do not conserve total carbon number between the complexes linked. Inclusion of GAP (*A*3) in a complex is a positive horizontal shift in the lattice, while input of DHAP (*k*3) or removal of Ru5P (*K*5) are respectively upward and downward vertical shifts. To represent the source *J*ext strictly we need only incorporation of GAP and elimination of Ru5P, but since all solutions use the TIM reaction twice to isomerize GAP to DHAP, we may combine those into a formal loop (shown in the final panel of the next figure) to “add” GAP to the empty complex, isomerize, and then “eliminate” DHAP so that we may then add it by hand to other complexes.

Taking the maps in Table IV and Table VI as the set of available elementary links, we show in Fig. 10 examples of a series of irreducible flow solutions as lattice maps. Each flow solution is a simple closed curve in the lattice, because it converts all incorporated inputs to eliminated outputs. Moreover, this series of solutions is *minimal* in the sense of using the fewest reactions solving the schema (34) from a seed *An*, entailed by the conservation laws of these rules.

**FIG. 10:**
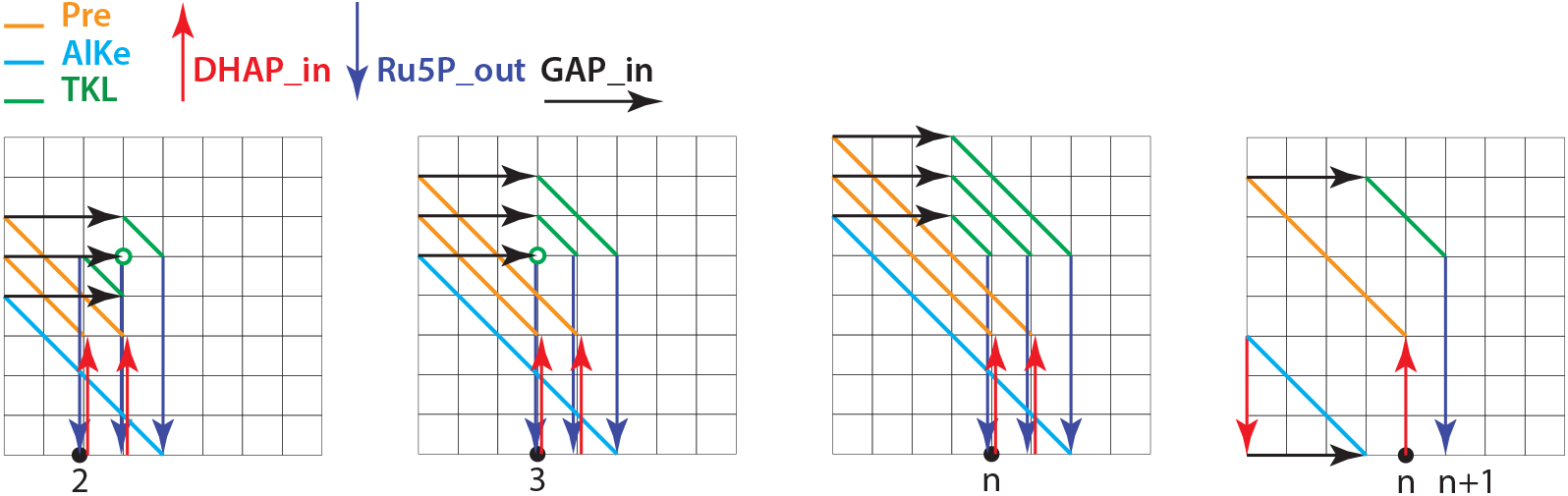
Lattice diagrams for a sequence of complexes that carry one autocatalytic *An* backbone through the sequence of its conversions in one full turn of the CBB cycle. To simplify the diagrams, the inputs are taken to be 3 *A*3 + 2 *K*_2_ (corresponding to 3 GAP+ 2 DHAP). First panel, starting with A2, is f14 from Fig. 5. Second panel, starting with A3 is f13 from the top panel of Fig. 3. Third panel, starting with *A*_4_ and representative of *A*_n_ for any *n* ≥ 4, is f194. The links in each path are to be read in the order that makes a closed cycle. The starting aldose is a 1-complex *A*_n_, on the lower axis (black dot). Additions of DHAP are red up arrows, forming the 2-complex input to PHL ° AL edges (orange), producing 1-complexes (ketoses) on the vertical axis. Additions of GAP are black right arrows, forming the 2-complex inputs to TKL edges (green). Extractions of RuB5 are blue down arrows, converting 2-complexes back to single aldoses on the horizontal axis, from which new complexes are created by DHAP addition. The sub-flow Ru5P_out ° TKL ° GAP_in ° PHL ° AL ° DHAP_in, shown in the fourth panel, forms one turn of an “algorithmic” loop incrementing the value *n* for the starting aldose 1-complex *A*_n_. AlKe conversions (cyan), followed by Ru5P_out ° TKL ° GAP_in in each of the first three panels, reset two iterations of this algorithmic loop to its starting complex. Also shown in the fourth panel is a 3-cycle of *A*_3_ → *K*_3_ which, added to the previous graphs, would permit a starting configuration of 5 *A*_3_ + 0 *K*_3_, at the cost of slightly greater complexity in plots.

More can be noticed about properties of solutions from their representation by lattice maps, but we remove most of that elaboration to App. D. Here we consider only two results: the surprising *algorithmic* character of an infinite series of solutions, and the origin of the minimum value of 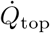 and the two solutions realizing it.

##### a. Algorithmic character of a series of mixed-prelude flow solutions

The open line segment shown in the fourth panel of Fig. 10 serves as a lumped transformation repeated in all the foregoing solutions. Its two links, one PHL ° AL and one TKL, add one DHAP and then one GAP to an *An* 1-complex, and eliminate one Ru5P, returning an *An*+1 1-complex. The series can be repeated indefinitely, converting C6 added into C5 eliminated and C1 that increments the aldose that serves as a *counting register*. In contrast, when a KeAl link is followed by GAP addition, TKL reshuffling, and Ru5P elimination, the result is a downward shift *An* → *An*−2. Therefore a minimal closed curve is formed by two repeats of the +1 function and one repeat of the -2 function.

Because none of these operations passes repeatedly through the same link,18 the three reactions in the +1 function contribute +3 to 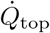 in Eq. (33), and two reactions in the -2 function contribute +2 to 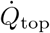, summing to +8 for all loops in the third panel of Fig. 10 at *n* ≥ 4. A remaining +4 is contributed to 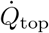 from the twice-used TIM reaction, shown as a simple closed curve through the origin in the final panel of Fig. 10. Thus 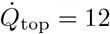 for all solutions in the third panel for *n* ≥ 4.

More complex solutions at larger 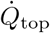 can be obtained by splicing simple closed curves corresponding to null flows into the series shown in Fig. 10. Because the three AlKe conversions required by the schema (34) and the TKL re-arrangements required by the lack of common divisor of *A*3 and *K*5 are already represented without overlap in the members of the serious, though, no addition of further loops lowers the resulting 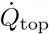.

##### b. The unique minimality of solutions f13 and f14

The sole exception to the value 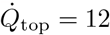 for the infinite series in Fig. 10, as a minimum enforced by conservation laws, arises for the flows f13 and f14 shown in the first two panels of the figure. As the “counting register” *An* that starts the series is decremented in *n* through values 3 and 2, exactly two of the TKL edges can be made to pass through the fixed-point complex (*A*3, *K*5), reducing 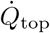 to the value 11. The third TKL edge can never be made to pass through this fixed point, because no C1 aldose phosphate exists as a realization of the formal label *A*1.

**Remark:** As a corollary we see that the two flows f13 and f14 with 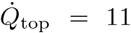 in Fig. 4 are also the only two in the series from Fig. 10 that are not network catalytic [59, 88] by necessity. Any curve passing through the TKL fixed-point complex (*A*3, *K*5) (GAP_in, then Ru5P_out) can be “re-routed” through the origin (Ru5P_out, then GAP_in); in the language of *siphons* [68], they are “self-priming”.

## IV. THE INFORMATION ASSOCIATED WITH LARGE DEVIATIONS AT SINGLE INSTANTS AND THROUGH TIME

We have so far considered the first generative transition in our three-level system: from generating rules to stoichiometric networks and the Kirchhoff decomposition of the possible currents on those networks. We now turn to the second generative transition: from the stochastic reactions on stoichiometric graphs to the ensemble dynamics they produce, and particularly in the large-number limit, macroscopic state and transport relations, and associated measures of work and dissipation.

We will frame these from the outset not as an outgrowth of mechanics and calorimetry as in classical approaches to thermodynamics [6, 89, 90], or even many modern approaches [58], but from the contemporary perspective based on information [60, 91–93] particularly in the form of relative entropies and, in large-number or other scaling limits, of large-deviation functions [43, 45]. These information divergences include as special cases the thermodynamic potentials of equilibrium systems, but they generalize much more widely: to ensembles of histories [46, 94] or coarse-grained models lacking reaction reversibility [32], and to multi-level or multi-scale population processes including biological evolutionary processes [29–31].

Fundamental concepts and basic constructions are introduced in their simplest forms, on distributions *ρ*n at single instants. These are then generalized to information divergences including large-deviation functions on ensembles of histories, which will be our main working context. These summaries will be brief; didactic treatments are widely available [29, 32, 46, 95–100]. Our new contribution is the application of natural information-geometric definitions of orthogonality [60] to flow reduction through hierarchies of nested subgraphs as developed in Sec. II D 2. We will show that differences in large-deviation rates between more permissive and more restrictive networks are precisely measures of the improbability for spontaneous occurrence in a network of the flow pattern that removing an edge from the network ensures, and thereby natural lower bounds on the information required to maintain the restriction through selection.

### A. Generating functions, Legendre duality, and relative entropy of a fluctuation at an instant

#### 1. Hartley information and the large-deviations relation of entropy to macro-fluctuation interpretations

The basic quantity from which our construction of information divergences will start is the cumulant-generating function (CGF) *ψ*(*θ*) for the number vector n under a probability distribution *ρ*n, defined as

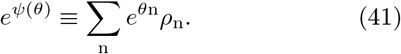

Here *θ* ≡ [*θ*_*p*_] is a (row) vector of (generally real) numbers indexed by the species index *p* of n, and *θ*n is the Euclidean inner product.

*e* ^*θ*n^ *ρ*_n_ is an exponentially *tilted* measure, which we convert to a probability by normalizing with *ψ*:

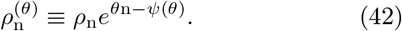

The distributions (42) form an *exponential family* over the base distribution *ρ* in which *θ* provide a coordinate system on the family.

The gradient of the CGF is the expectation of n in the distribution 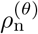:

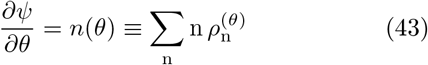

We note for later use that if 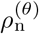 is monomodal and sharply peaked, it will have a saddle point near *n*(*θ*) with value ∼ 1, where ∼ denotes leading exponential dependence on *n*(*θ*).

By convexity of *ψ*, the function *n*(*θ*) is invertible to a function *θ*(*n*) ^19^, and the Legendre transform of *ψ* is is also its *Bregman divergence* about 0 (note that *ψ*(0) ≡ 0), which we denote

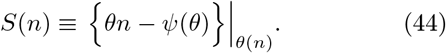

By construction the gradient of Eq. (44) gives the inverse function

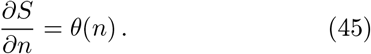

From the definition (42) it is immediate that

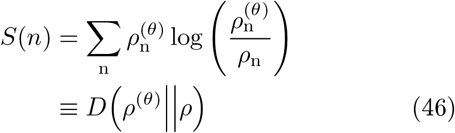

is the *Kullback-Leibler (KL) divergence* [101] or negative of the relative entropy of *ρ*^(*θ*)^ referenced to *ρ*.

Note the relation between *S*(*n*) and the tilted measure, that

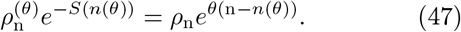

In the same leading-exponential approximation where 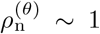 at its saddle point, we have the envelop approximation

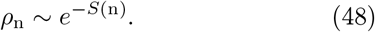

As an approximation to − log *ρ*n, *S*(n) is an instance of a *Hartley information* [102].

The leading-exponential approximation of expectations in *ρ* or *ρ*^(*θ*)^ by their saddle-point values is known as the large-deviation limit [45] and in this limit *S*(*n*)| _*n*=n_ appearing in the envelop approximation (48) is called the *large-deviation function* (LDF). Our designation is motivated by a construction that will be possible for *S* as a Lagrange-Hamilton *action* functional that we develop below.

The relations (41,48) are illustrated in Fig. 11. From the figure we see that the KL divergence, in contrast to the microscopic probability *ρ*n to occupy any state n, measures the divergence of the distribution *ρ*^(*θ*)^ in which *n*(*θ*) is *typical* [101] from the distribution *ρ* in which *n*(0) is typical. Distributions such as *ρ*^(*θ*)^ produced by tilting provide a natural definition of *macrostates* [32], and for this reason −*S*(*n*) carries the interpretation of the log-probability for a macrostate fluctuation.

**FIG. 11:**
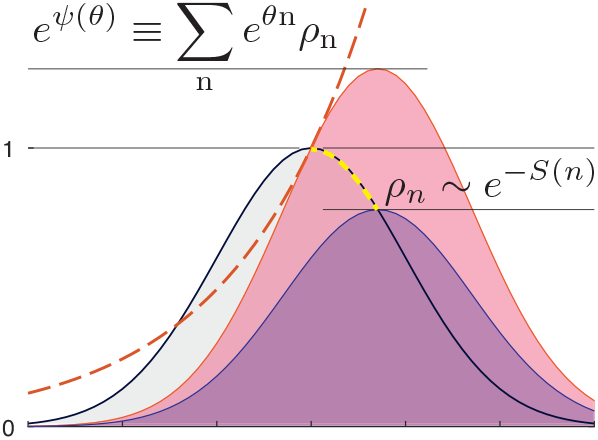
Shaded grey with black outline is the original distribution *ρ*_n_. Red dashed is likelihood (or “tilt”) 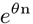. Shaded pink is 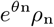 with local maximum near *n*(*θ*) at value ∼ *eψ*(*θ*). Shaded purple represents the distribution 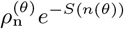 of Eq. (42), with local maximum ∼ *ρ*_n_|n=*n*(*θ*). Heavy yellow dashed line between *n*(0) and *n*(*θ*) traces, in logarithmic coordinates, the integral ∫ *θ dn* derived below in Eq. (58).

#### 2. Dual convexity of the CGF and the LDF

In any region where the function *n*(*θ*) is invertible, the Legendre transform (44) as a Bregman divergence converts convexity of *ψ* into convexity of *S*. From Eq. (43), the Hessian of *ψ* is the Jacobian of a coordinate transform from *θ* to *n*:

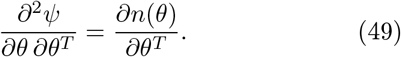

The corresponding Hessian of *S*, from Eq. (45), is the Jacobian of the inverse transform:

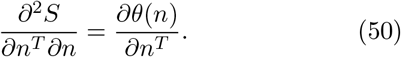

It follows that

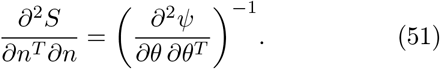

These two matrices are the Fisher-Rao [103, 104] metric of information geometry [60], in contravariant and covariant coordinates.

#### 3. The Hamilton-Jacobi equation for the past history of generating-functions with final-time conditions

The CGF and LDF of the previous subsection are computed for a base distribution *ρ* considered in isolation. In the case that *ρ* arises as a single-time evaluation of a distribution evolving under a master equation (1), the same information divergences at a single-time can be computed from an integral construction, which attaches a macro-fluctuation probability to the distributional *history* conditioned on arriving at a single-time deviation.

In the large-number limit where saddle-point approximation can be used, the CGF and LDF evolve under a *Hamilton-Jacobi equation* [64–66], which is the ray-theoretic approximation [69, 70] for the bundles of histories that dominate the probability of the least-improbable histories that arrive at improbable states. The resulting Hamiltonian dynamical system with its conserved volume element is the saddle-point representation of the conservation of probability [105] familiar from Liouville’s theorem [63].

When the distribution *ρ* evolves under the master equation (1), its associated CGF (41) evolves under a differential equation in *θ* and *∂/∂θ* known as the *Liouville equation* [63]. In general this equation couples moments of all orders under *ρ* and requires either all-orders solution or some moment-closure scheme [39, 40] to be solved. However, in the saddle-point approximation (corresponding to the mean-field equation in the rate law (7)) where the effect of *∂/∂θ* can be replaced with *n*(*θ*) through Eq. (43), the Liouville equation is closed to leading exponential order as

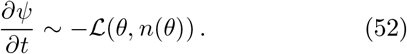

Henceforth we will work only in this saddle-point approximation, noting that it can be extended to formally exact solutions by functional integral methods explained in [29, 32, 46, 95–100].

For a stochastic process generated by a transition matrix with the stoichiometric, mass-action form (5), the associated Liouville function for Eq. (52) evaluates to

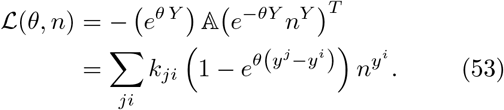

Equivalently, from the Legendre transform (44), *S*(*n*) evolves as

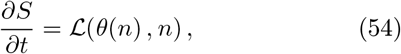

recognizable as a *Hamilton-Jacobi* equation [3] with Hamiltonian − ℒ, when we substitute *θ*(*n*) for its evaluation (45) as the gradient of *S*. The equations (52) and (54), together with (43) and (45), define the Legendre-dual system of the CGF and LDF. *S*(*n*) fills the role of *Hamilton’s Principle function*, which we construct in the form of an action integral in the next subsection.

#### 4. Cumulant-generating and large-deviation functions as extended-time integrals from the Hamilton-Jacobi equations

From a functional form *ψ*0 that serves as initial data at any time *t* = 0, Eq. (52) may be integrated in time and along the first-derivative condition (43) to produce a function *ψT* at a later time *t* = *T* in the integral form

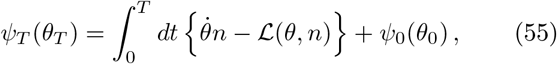

where the integral is evaluated along time-dependent functions *θ*(*t*), *n*(*t*) that are solutions to vanishing of the first variational derivative:

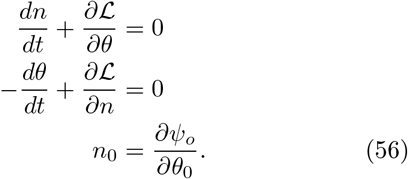

The Legendre transform (44) requires only an integration by parts in Eq. (55) to absorb surface terms, giving *S* in the integral form of Hamilton’s Principle function

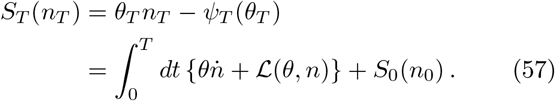

In this paper we will not study the Hamilton-Jacobi representation of transient distributions, but will consider only steady states. For these, from Eq. (52), ℒ = 0 along all stationary trajectories. Let *n*_*o*_ be the saddle point of *ρ* in the stationary distribution, and therefore the minimizer of the LDF at *S*(*n*_0_) = 0. Then *S* at any other value *n* is evaluated as

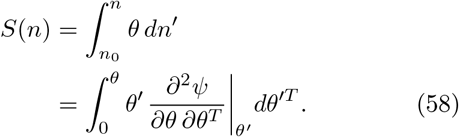

Convexity of *S* ensures that the trajectory 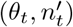 between *n*_0_ and *n* reduces to a con/tour *θ*(*n*′) that we term an *escape path*. The expression ∫*θ dn*′, which follows by direct integration of Eq. (45) in the single-time construction (shown as the yellow dashed in Fig. 11), has been arrived at in Eq. (57) through a time integral from a fixed point *n*_*o*_ in the asymptotic past.

The second line of Eq. (58) follows from Eq. (49). From the definition (41) we observe that the Hessian

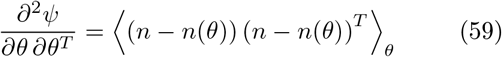

is the covariance matrix in the distribution *ρ*^(*θ*)^ of Eq. (42), and the Fisher-Rao metric [103, 104] on the exponential family of distributions produced by tilting of the stationary distribution as the base distribution *ρ. ∂*^2^*S/∂n*^*T*^ *∂n* is then the matrix inverse by Eq. (51).

The relations derived in this section between *ψ* and *S*, and between the Legendre-dual variables *θ* and *n*, for a distribution at a single time, will be recapitulated in the large-deviation rate for persistent departures of currents from their mass-action laws when we study extended-time large-deviation functionals in the next section.

#### 5. Escape paths, the large-deviation function, and dissipation

The CGF for an ordinary Gibbs equilibrium distribution tilted by chemical potentials is the *grand potential* [106], and the LDF is the Helmholtz free energy (both scaled by *k*_*B*_*T*). It is conventional [24, 58] from the energetic view to describe relaxation toward a thermodynamic equilibrium as “dissipating” free energy to heat, with the grand potential viewed as a system resource that is “dissipated”.

The construction (58) of *S* from integrals along escape paths, however, makes only incidental reference or no reference at all to the “relaxation” path that is a solution to the mass-action rate equations without source, depending on the balance class of the generator in the terms of Sec. II C 2. Indeed, a derivation is required to show why and in which approximations this *S*_*T*_ is a Lyapunov function for spontaneous relaxation dynamics, and why it should connect to any notion of dissipation.

It is a general result [37] that the full large-deviation function (46) on distributions is a Lyapunov function for general relaxations under the free process. For the less-general reduction to the saddle-point approximation, it can be shown [32] that the sign of 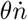along relaxation paths is opposite to the sign along escapes. Since 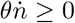 along escapes by construction, ensuring non-negativity of *S*_*T*_, we have that *S*_*T*_ is a Lyapunov function for relaxation trajectories that constitute the leading-exponential contribution to relaxations. ^20^

Any stronger result is class-dependent: For systems with detailed-balance steady states, the relaxation paths are time-reverses of escape paths, and the construction (58) can be literally interpreted as the time-reverse of “dissipation”. For systems with complex-balanced steady states, the value of 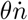 along escape and relaxation trajectories is exactly opposite, though the trajectories may depart from time reversal by “torsion” terms orthogonal to *θ*. For more general cases, only reversal of the sign is assured.

We note, finally that *ψ*_*T*_ in Eq. (55) and *S*_*T*_ in Eq. (58) are identically zero for any relaxation trajectory 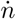 at *θ* ≡ 0, assigning no resource to the initial condition to be “dissipated”, and the leading-exponential likelihood 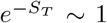 is independent of the degree of divergence of the initial value of *n* from its late-time steady-state value.

Thus we argue that the interpretation of energy dissipation to heat as a measure of irreversibility is contingent on detailed balance, and even in that case arises only because the Boltzmann factors (11) giving the rate constants are themselves large-deviation probabilities in the even-more-microscopic process of thermal fluctuation that describes thermalized reservoirs (See Fig. 12).21 All that relaxation paths do in the general case is surrender large-deviation improbability for which the measure is the cumulative unlikelihood along escape paths.

**FIG. 12:**
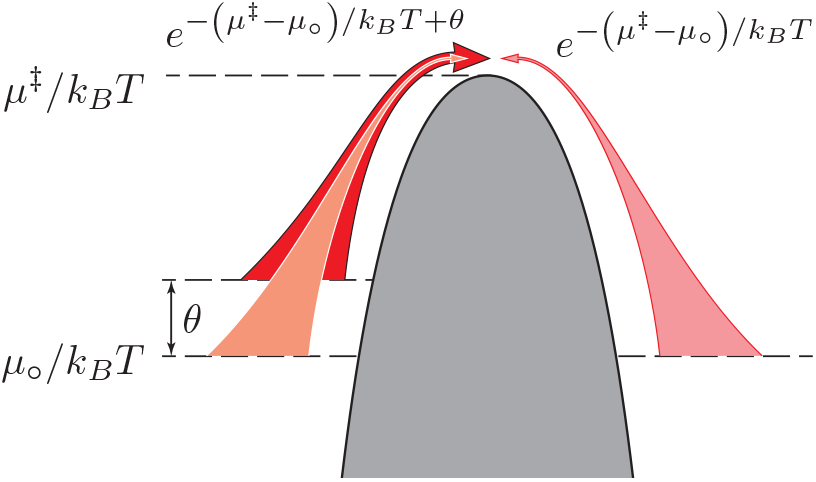
The microstate landscape (grey region) of a reaction with barrier *μ*‡ between two single-particle chemical potentials *μ*_°_. Balanced left-ward and right-ward currents 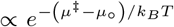 (light red) result from the exponential attenuation of thermal-fluctuation large-deviation probabilities at the top of the barrier. A tilting parameter equivalent by the thermal activation scale to a shift in free energy, *θk*_B_ *T* ↔ ∆*μ*, shortens the range of exponential attenuation of escape probability, resulting in a current 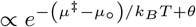 (dark red).

To have established this probability-first view of stochastic causation, relegating energy flow to a contingent and system-dependent role, will be useful in Sec. IV E when we evaluate the large-deviation functional for steady-state currents in relation to the conventional heat dissipation, to which it converges in limited cases.

### B. Conditioning along the course of trajectories: cumulant-generating and large-deviation functionals

The likelihood measured by *S*(*n*) in the previous section was evaluated as an action integral, but was for only one deviation value *n* imposed at a single time *t* = *T*.

Being now in possession of the integral form, we recognize that a large-deviation function can be computed for any deviation of a probability distribution over its history, from the natural stationary distribution or relaxing solution under the bare master equation (1).

We proceed in the construction as for *ψ*(*θ*), but now we must introduce tilt parameters that act along the course of an evolving distribution to form generating function*al* s on tilting histories, and their functional Legendre transforms that are KL divergences between untilted and tilted distributions on histories. We will present the abstract construction for general time-dependent tilts and responses, but only evaluate particular solutions for the steady states considered in previous sections.

#### 1. Tilting reactions to produce null cycles on closed graphs

On a non-equilibrium process we can construct generating functionals and large-deviation functionals both for counts *n* as before and also for reaction fluxes *v*. To bias sampling for reaction events we modify the Liouville representation (53) for the generator to a form

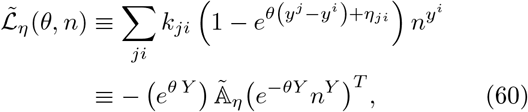

in which {*η*_*ji*_} are independent parameters (potentially time-dependent) on all unidirectional reactions. They may be thought of as activities of special fictitious species that augment the complexes in each (directed!) reaction, and that are buffered in external reservoirs used as controllers. To simplify the treatment here, we only consider tilts that are antisymmetric on bidirectional reactions, meaning that *ηji* ≡ −*η*_*ij*_. ^22^

The quantities recognized as reaction velocities from the first stationarity equation in Eq. (56) are, under the tilted generator,

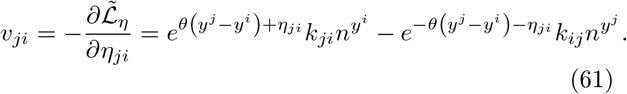

The altered generator (60) is sufficient to perturb both counts and currents because the full vector *η* ≡ [*η*_*ji*_] may be written as a sum of two terms: one the difference of a potential on complexes and the other a pure circulation. Any component of *η* derivable from a potential can be absorbed into a shift of *θ* in Eq. (60), as we can see from the form (61) and from the gradient relation

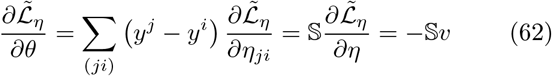

Any remaining components of *η* that cannot be canceled in this way induce null currents on the stoichiometric graph.

**Remark: gauge symmetry on closed graphs and the** *θ* 𝕊 = 0 **gauge:** Only the combination *θ* 𝕊 + *η* determines the current in the steady-state solutions at the instantaneous values of the generating parameters. ^23^ The model including all stationary solutions is thus invariant under a *gauge transformation*

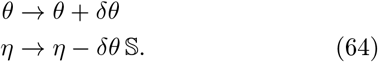

At times in the later derivation we will take *δθ* 𝕊 equal to the entire stationary value of −*θ* 𝕊, absorbing this into the initial choice for *η*. That choice defines the *θ* 𝕊 = 0 gauge.

A new cumulant-generating functional on the underlying stochastic process is written in its saddle-point approximation by adopting the integral form (55), now with the generator (60),

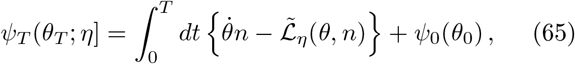

and evaluated on the corresponding stationarity equations that generalize Eq. (56) as

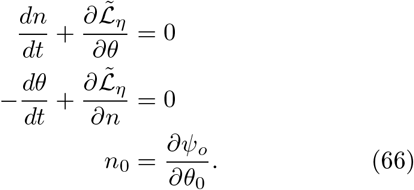

We can check that *ψ*_*T*_ is indeed a generating function as before for *n*_*T*_ and also now a generating functional for currents at any *t* ∈ [0, *T* ] by evaluating the first-order variations that generalize Eq. (43):

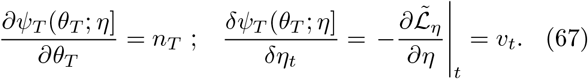

The second-order variational derivative with respect to *η*, a functional counterpart for currents to the single-time Hessian (49), is given by

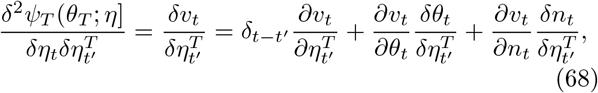

in which 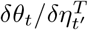 and 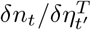 are the variations in the trajectory solution (66) at any time *t* resulting from variation of *η* in 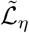 at a time *t*′.

#### 2. Functional Legendre transform and variations of the resulting large-deviations functional

We have from Eq. (67) that *v* is the Legendre dual variable to *η*, and the functional Legendre transform of *ψ*_*T*_ proceeds by the same construction as the single-time Legendre transform except that the product term constructing the Legendre transform as a Bregman divergence is now the integral of *ηv*, yielding

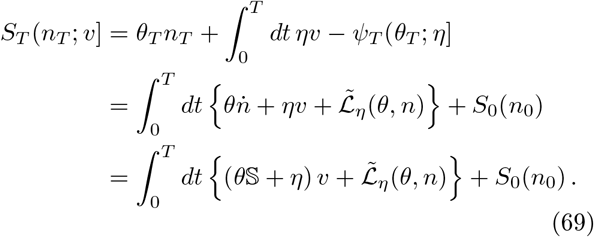

The third line of Eq. (69), emphasizing that *S*_*T*_ like 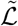 depends only on the gauge-invariant combination *θ* 𝕊 + *η*, follows from the second line by the stationarity equation for 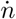 in Eq. (66) and Eq. (62). The first variational derivative of *S*_*T*_ evaluates to

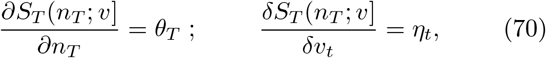

establishing that *S*_*T*_ is a potential for the dual variables to *n*_*T*_ and *v*.

The second functional variational derivatives of *ψ*_*T*_ and *S*_*T*_ multiply to the identity now also on their time indices,

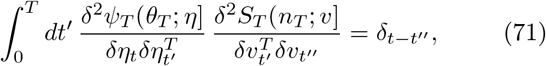

so we write these formally as a matrix inversion, generalizing Eq. (51) for the single-time construction from Sec. IV A 2, to

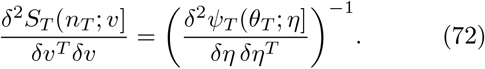

All information-divergence constructions, introduced in simple form for a distribution *ρ*_n_ at a single time on countable indices {n}, are thus generalized to divergences for continuous functional deviations on currents in an extended time interval.

#### 3. Integral expression for the functional Bregman divergences

Evaluating a non-stationary functional CGF or LDF for currents entails either solving Hamiltonian equations (66) with unstable directions both forward and backward in time [69, 70, 107–109], or possibly functional gradient descent [110], either method involving considerable numerical effort at stabilization. Here we perform the evaluation only for steady states, where the rate of accumulation of improbability along paths involves an expression in −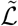 analogous to that from *ψ* for single-time fluctuations.

The functional Bregman divergence giving *S*_*T*_ in the third line of Eq. (69) is shown in App. G 2 to have an integral expression in terms of increments of current deviation *dv*′ under *η* in the form

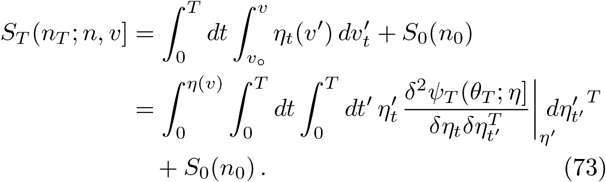

Eq. (73) is the functional generalization for currents – with *t* and *t*′ merely further matrix indices – of the integral representation (58) of the single-time Bregman divergence for counts *n*. The concept of “escape”, however, in the function *η*(*v*′), is not derived from a trajectory in time, as *η* and *v*′ are already source and current histories defined on the full time interval.

The restriction of Eq. (73) to the case of time-independent sources *η* that create deviations through sequences of steady states evaluates to

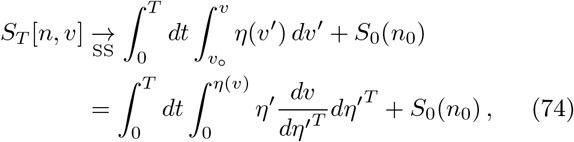

and it is shown in App. G 2 a that for this case

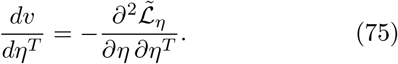

We note the parallel formal role of the Hamiltonian 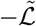, with respect to the *temporal density* in measure *dt* in Eq. (74), to that of *ψ* in the single-time expression (58).

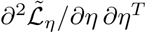 is diagonal in the reaction index *ji*, and will be negative definite if all reactions have at least one half-reaction current. In that case the matrix (75) is invertible, and the relation between the Hessian and its inverse is

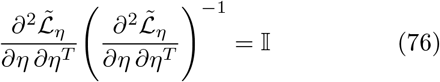

the identity matrix on reactions.

If we work in a gauge for which the explicit component *η* ≡ 0 on some reactions in a subgraph, but *θ* 𝕊 ≠ 0 across those reactions, the relation (62) allows us to write the potential-part of the Hessian (75) as

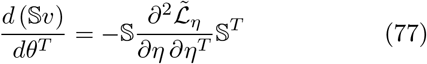

The matrix (77) will be used in Sec. IV E 4 to compute solutions to 𝕊 *v* = *J*ext within the hierarchies of subgraphs used for flow reduction. Because im S is the stoichiometric subspace, it will generally have a kernel. Where an inverse is required, that must be constructed within the stoichiometric subspace as explained in App. F.

#### 4. Functional “escape paths” and a complete disconnection from relaxation as dissipating an energy resource

The integrand of the double integral in the LDF of Eq. (74) admits the interpretation of the differential of a dissipation rate, with *η*(*v*) as an auxiliary chemical potential difference across reactions required to maintain a current profile *v*, and *dv* the increment of divergence of that current profile from its value *v*_°_ at *η* ≡ 0. Minimizing dissipation at the solution *S* = 0 is just Onsager’s [81, 82] principle, appropriately extended beyond linear response.

We argued in Sec. IV A 5 that the information divergence is fundamental and also general, whereas the interpretation of dissipation (of energy) is derived and in chemistry is contingent on the detailed-balance property of networks from Sec. II C 2. The source of that argument was the construction of the LDF in Eq. (58) from escape paths in the action functional (57), for which the tangent 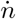 points in the *same* direction as the increase of *θ*, rather than in the opposite direction as is true for relaxation paths relative to the chemical potential.

Whereas, however, the time coordinate along an escape trajectory to a single-time fluctuation was a “removable” coordinate provided by the Hamilton-Jacobi solution and connected along its length to the imposed information condition only as a final-time boundary condition, in the LDF for currents, time is a coordinate along the entire length of which independent information conditions are imposed. A steady-state current compounds improbability everywhere along its length that it deviates from the mass-action current solution. The “escape contour” in the current LDF is now a functional contour *dvt* in which infinitely many increments of “dissipation” *η dv* (the rate of compounding of trajectory improbability) are connected to previous increments by the trajectory probability measure.

Thus for currents the interpretation of an energetic “resource” that is “dissipated” under free relaxation in time is entirely dispensed with. The compounding rate for trajectory improbability scales with *v* – the rate at which any events happen *at all* – and with sampling bias *η* against the background trajectory needed to generate the divergent rates.

#### 5. Factoring subgraph from environment

Our strategy for using the LDF (74), to characterize flow reduction through nested subgraphs in terms of information, is to produce general *null* current profiles on a closed graph 𝒢 – the only kind that can arise as steady states – using the uniform device of sources *η* on all reactions, and then by isolating the parts of these currents that flow through any subgraph, to associate their contributions to the LDF from inside the subgraph with the source currents *J*^ext^ on its boundary, conditionally independent of the way the closure of the flow is generated in the environment. To perform this two-step analysis we must decompose the LDF for currents on a subgraph and its complement as defined in Sec. II D.

Let 𝒢′⊂ 𝒢. To streamline notation, in this section let 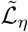 refer to the partial sum in Eq. (60) over reactions in the subgraph 𝒢′, and 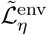 the sum over the remaining reactions in the complement (𝒢′)⊥ ⊂ 𝒢. The generator for all of 𝒢, which we denote 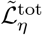, decomposes as

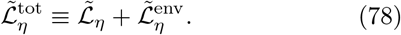

We add and subtract surface terms that are nonzero only on the species in *∂* _𝒢_ 𝒢′, in the form of Legendre dual pairs in both *n* and *θ*,

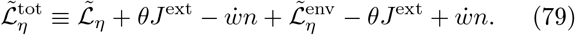

*J*^ext^ and 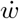are not new source terms in addition to *η*; rather they are chosen to define a separating hyperplane between the subgraph and its environment. We assign them values for which, on the solution *v* of interest in the whole graph 𝒢, the restriction of *v* to 𝒢′ will be a solution to the variational equations on the subgraph species

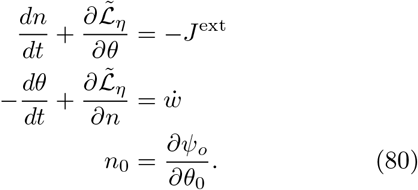

Eq. (80) are the stationarity conditions for a CGF defined only on 𝒢′ with the form

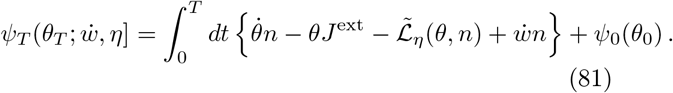

**Remarks:**

1. Recalling Eq. (41) for the single-time CGF of counts *n*_*T*_ formed with tilting parameter *θ*_*T*_, 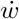 is the generalization of such a tilting parameter to extended time. As a density in measure *dt*, its units are those of a *rate* of the chemical potential *θ*. Here that source term results from marginalization of the environment with respect to the gradient on boundary-species *n*.
2. A new term, not applicable to an instantaneous distribution that is not embedded within a history, is *J*^ext^, resulting from marginalization of the environment with respect to the gradient on the conjugate momenta *θ* to obtain a CGF on the part of currents in the subgraph derivable from a potential.
3. We show in Eq. (G53) of App. G 3 a that, for 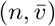 the species and current profiles on a subgraph driven by 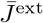, the same profile *n* will form a steady-state in the CGF with generator 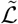, along with reversed velocity 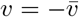, reversed source current 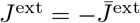, and 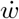and *J*^ext^ having the relation 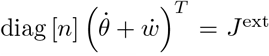. ^24^ These results are implied only for a generator with detailed-balance solutions at *η* ≡ 0. The reversal of *v* and *J*^ext^ in the tilted stationarity equations (80) that construct the LDF *S*_*T*_ has the same origin as the time-reversal of escape paths relative to relaxation solutions to the mass-action rate law in Eq. (56), discussed in Sec. IV B 4.

The functional Legendre transform of the CGF (81) on the subgraph 𝒢′ is then shown in Eq. (G46) of App. G 3 to take the form

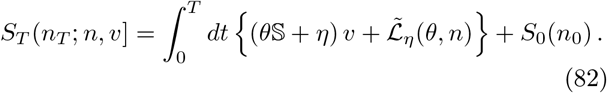

The term 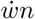 in the functional Legendre transform has canceled the source term of the same form in the integrand of Eq. (81), in the same way that *θ*_*T*_ *n*_*T*_ cancels a total derivative in the Legendre transform (69) for single-time deviations. The term *θ*𝕊*v* = *θJ*^ext^ is the uncanceled remnant of marginalization from Eq. (81). The term *ηv* is the product of dual Legendre variables making *S*_*T*_ a Bregman divergence.

It is important that the form (82) of the LDF on the subgraph is the same as the form (69) for the full graph, differing only by the reactions in the sum, (*ji*) ∈ 𝒢′ versus (*ji*) ∈ 𝒢. This separability makes the cost of a deviation in G𝒢′ conditionally independent, given *J*^ext^ and 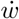, of any way the source current might be created by choices within the complement (𝒢′) ^⊥^. Such independence allows us to use null flows in the full graph 𝒢 to define the feasible conversions while studying the cost of their effect as functions of properties of the subgraph.

##### a. Gauge invariance and gauge choices

We verify that, as in the constructions of Sec. IV B 1, only the gauge-invariant combination *θ*S + *η* determines currents in steady-state solutions. To provide a simpler notation for several expressions in the next section that use this result, we will adopt the notational shorthand 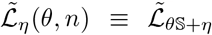. The extension of the gauge invariance (64) for null flows, to the subgraph including through-flows absorbed by boundary currents, is

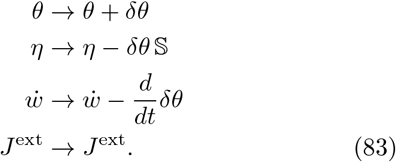

**Remarks on gauge choices:**

1. One gauge choice for *η* sets *θ*S ≡ 0, putting Eq. (82) directly in the form (73).
2. An alternative use of gauge freedom to setting *θ*𝕊 = 0 is to set only the components of *η* coupling to null cycles *within the subgraph* non-zero, so that the two potentials *θ* and *n* are dual variables to the boundary conditions *J*^ext^ and 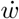 on *∂*𝒢 𝒢′.

### C. Information-geometric decomposition of the steady-state cost function

We can now define a protocol to compute the large-deviation function for the current response to a feasible conversion 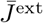 performed through a subgraph 𝒢′ by any combination of through-flows and null flows in 𝒢′. Of particular interest will be the additive contribution from the flow solution to the mass-action rate law and from additions to it by null flows to create a mass-action flow supported on a proper subgraph of 𝒢′.

1. Having shown that the separation (81) makes large-deviation behavior in the subgraph 𝒢′ conditionally independent of the environment given boundary conditions *J*^ext^ and 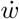, we first find steady-state solutions *v*′ and *n*′ to 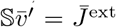 under the massaction rate law at *θ*𝕊 + *η* = 0, at each point 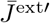along a contour from 0 to some final value 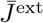.
2. The elementary construction (62), written out in detailed form as Eq. (G3) of App. G 1, shows that, for every such 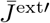, a value *θ*𝕊 + *η* ≠ 0 exists (in the subgraph) for which solutions 𝕊*v*′ = *J*^ext^′ have the same profile *n* and opposite velocities 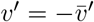 and source current 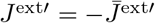 from the source-free solution. If the rates in the subgraph have detailed balance, that velocity-reversing tilt is gauge-equivalent to *η* ≡ 0. If the subgraph rates do not have detailed balance, some *η ≠* 0 will generally be required for velocity reversal. In either case, the value of *θ*𝕊 + *η* ≠ 0 is specified by preservation of the profile *n*. The boundary values 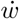, satisfy a relation to *n* given in Eq. (G53), with 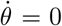 for the detailed-balance case.
3. The velocity-reversing profile serves as a point of reference relating 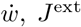 and *n* through a simple reversal of signs from their mass-action solutions in Eq. (80). Only for detailed balance is the contour of these solutions the one that gives the LDF; more generally an additional step in the construction will be required (next):
4. If the subgraph rates do not have detailed balance, the velocity-reversing solutions differ by a null flow in the subgraph from some solution that is gauge-equivalent to *η* ≡ 0 and so we may construct the latter by subtraction of the needed *δ*_*η*_.^25^ The resulting solution will still be stationary but will no longer generally preserve *n* from its profile at *θ*𝕊 + *η* = 0, and the relation of *J*^ext^ to 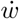 and 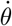 will become the more complicated form (G54), within which 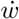 and *J*^ext^ do remain static and finite up to possible additive terms proportional to the conserved quantities.
5. ***Key point in the construction:*** The solutions that are gauge-equivalent to *η* ≡ 0 are the essential construction to enable the decomposition of the current LDF by the extended Pythagorean theorem explained next in Sec. IV C 1.
6. We advance the environmental driving rates and solve for these potential-only tilts until the current through the subgraph is 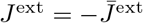.
7. To obtain the final steady-state LDF (74) for any flow *v* through 𝒢′ with *J*^ext^ = 𝕊*v*, we add any further components *η* coupling only to null flows in the subgraph to produce the desired *v*.
8. The Pythagorean decomposition is simple and has a uniform form for subgraphs with detailed balance. Otherwise it admits existence proofs but may be hard to compute. This is the usual situation with CRNs not restricted to detailed balance.

#### 1. Extended Pythagorean theorem for the cost function as a Bregman divergence

Following the above program for computing LDF integrands of the form in Eq. (82), for general sources 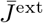 on a graph and general conservative flows within it, we show in App. G 4 that these integrands satisfy a decomposition known as the *extended Pythagorean theorem* in Information Geometry [60, 111]:

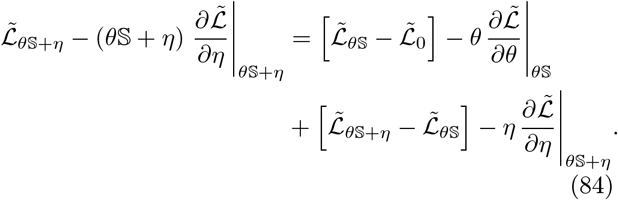

The left-hand side behaves like the squared length of the hypotenuse of a right triangle, and the two terms on the right-hand side like the squares of the base and rise in Euclidean geometry. The interpretation is that the source path from no current (*θ*𝕊+ *η* = 0) to the current for an arbitrary source *θ*𝕊+*η* can be broken into two legs – from 0 to *θ*𝕊 and from *θ*𝕊to *θ*𝕊 + *η* – and that these two legs are orthogonal under the natural geometry induced by the large deviation function.

The first line of Eq. (84) defines a potential-only trajectory, as flow solutions under the mass-action rate law are built up in response to increments *dJ*^ext^ leading from zero to the final value *J*^ext^ at *θ*𝕊. For subgraph rates with detailed balance, *θ*^*T*^ = log (*n/n*) is derived directly from the mass-action solution with 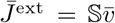 and the stoichiometric compatibility class of the equilibrium *<u>n</u>*.^26^

The second line of Eq. (84) is straightforward by comparison, involving only addition of null flows to produce the desired *v* (and its consequent *n*) keeping fixed *J*^ext^ = 𝕊*v*. Because null flows result in no dissipation against any chemical potential (as we noted in Sec. II D 5), the possible cross-terms in writing the left-hand side of Eq. (84) as two summands vanish, and only the remaining terms on the right-hand side are nonzero.

The individual orthogonal legs of these large-deviation integrands can then be written as the integrals of a quadratic form, as we did in Eq. (58) for the single-time case, along the contours for *θ* and *η* identified above:^27^

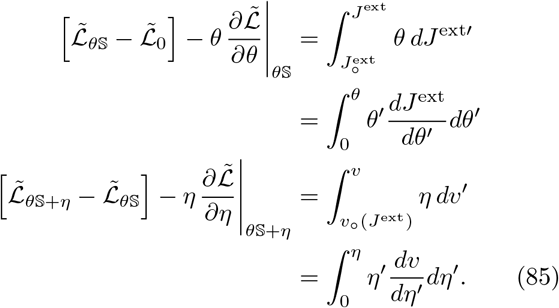

**Corollary:** Because both terms in the Pythagorean decomposition (85) are non-negative, we identify the source *θ*𝕊 and its associated flow as the minimum-dissipation solution for a given subgraph 𝒢′.

### 2. Triangle inequalities between graphs and subgraphs

The extended Pythagorean theorem (84,85) provides a systematic decomposition of information-divergence contributions to an LDF along the sequence of nested graph restrictions introduced to realize flow reduction in Sec. II D 7. We illustrate here for one step in such a sequence, in which two flows *v*_1_ and *v*_2_ both solve 𝕊*v* = *J*^ext^ for a common *J*^ext^, but the two differ in being solutions under the mass-action rate law on a graph and its proper subgraph. The steps in the decomposition are these:

1. Let 𝒢_2_ ⊂ 𝒢_1_ ⊂ 𝒢 be a nested hierarchy of proper subgraphs in 𝒢.
2. Let *J*^ext^ be a source obtained from a null flow *v* on 𝒢 by the separation in Eq. (79) at *∂*_*𝒢*_ 𝒢_1_. Suppose that *v* passes through both 𝒢_1_ and 𝒢_2_, and that the support of *J*^ext^ is contained in *∂*_*𝒢*_ 𝒢_1_ ∩ *∂*_*𝒢*_ 𝒢_2_. Thus both 𝒢_1_ and 𝒢_2_ can carry out the net conversion *J*^ext^.
3. We consider a trajectory *J*^ext′^ of source currents with support in *∂*_*𝒢*_ 𝒢_1_ ∩ *∂*_*𝒢*_ 𝒢_2_, originating in the the undriven steady-state current 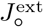(which will be zero for networks with detailed balance) and terminating in the final driven current *J*^ext^ of interest.
4. Within the two subgraphs 𝒢_1_ and 𝒢_2_ there will be two paths of current profiles 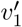 and 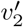 with 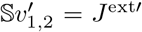, which are solutions under the mass-action rate laws at *η* ≡ 0 on their respective graphs, with corresponding paths of the *θ*-profiles *θ*_1_(*J*^ext′^) and *θ*_2_(*J*^ext′^).
5. Note that the graph 𝒢_1_, by construction, supports flows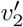, but the converse (𝒢_2_ supporting any 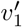) will generally be impossible for mass-action solutions. Therefore a path/ of sources *η*′ exists, starting at 0 and having ∫*dη*′ = *η*, coupling only to null cycles within G_1_, for which a corresponding path of current profiles *v*(*θ*_1_(*J*^ext^) 𝕊 + *η*′) has initial and final values *v*(*θ*_1_(*J*^ext^) 𝕊) = *v*_1_ and *v*(*θ*_1_(*J*^ext^) 𝕊 + *η*) = *v*_2_, the respective endpoints of the paths 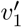 and 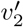 in 𝒢_1_ and 𝒢_2_.

The extended Pythagorean theorem (85) for the two integration paths to reach the same final current profile then requires that

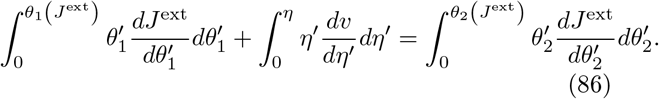

in which the two integrals on the left-hand side are performed in the subgraph 𝒢_1_ and the integral on the right-hand side is performed in the subgraph 𝒢_2_. Eq. (86) can be re-arranged as

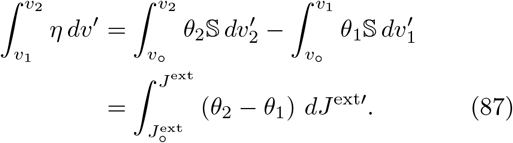

As noted in Eq. (19) for all null flows, the null deviation of the current at *θ*𝕊+ *η* from the current at *θ*𝕊 dissipates no net chemical work itself; rather it redistributes chemical potential to match the altered mass-action rates, in the process changing the potentials in *∂*_*𝒢*_ 𝒢_1_ against which *J*^ext^ does chemical work.

### 3. Example from the sugar-phosphate network

To produce a large-deviation decomposition parallel to the Kirchhoff current decomposition shown for the combinatorial sugar-phosphate network in Sec. III, we must first attach thermodynamic and kinetic parameters to that network to define mass-action rate constants. We will do this in a very simple way drawing from thermochemical data sources for sugar-phosphate molecules, summarized in App. E; a more complete treatment emphasizing data properties and their evolutionary implications is left to other work.

We obtain free energies of formation to produce standard-state chemical potentials *µ*_°_ for species from the thermochemical computation platform eQuilibrator 3.0 [62], and from these compute concentrations in a thermodynamic equilibrium near physiological temperature, pH, ionic strength, and metabolite concentrations. To assign kinetic parameters we adopt a limiting case of “barrierless” kinetics, in which the half-reaction rate constant from the complex with lower activity is set to a constant value over all reactions that we take to be 1 in appropriate rate units. The half-reaction rate constant from the complex with higher activities is then the equilibrium constant. This assumption is equivalent biologically to the assumption that all enzymes have evolved turnover rates up to the diffusion limit for substrates to the enzymes.

From the resulting thermodynamic and kinetic landscape, we compute steady-state solutions under the massaction rate law (7) on variously nested subgraphs of the network from Fig. 2. The net conversion *J*^ext^ is, as elsewhere, that of schema (34).

#### a. Triangle inequalities along transects connecting irreducible flows by single null cycles

We can both illustrate the extended Pythagorean theorem in simple plots, and also identify the degrees of freedom in which the Kirchhoff current decomposition is sensitive to the thermochemical landscape in mass-action solutions, by studying 1-parameter families of flow solutions on the supporting graphs connecting pairs of irreducible flows. We draw here from the examples introduced in Table. III.

We consider first the reduction of a flow by elimination of the single 235 TKL trefoil, shown in the second panel of Fig. 7, that connects flows f14 and f193. Here 𝒢_1_ is the union of supporting graphs for the two irreducible flows, and *v*_1_ is the mass-action solution to 𝕊*v* = *J*^ext^ on 𝒢_1_, which we denote *f*_14∪193_. 𝒢_2_ may be the supporting graph for either irreducible flow in which case *v*_2_ is either f14 or f193.

We illustrate results in the linear-response regime, where the two integrals 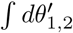in Eq. (86), along trajectories of mass-action solutions in either 𝒢_1_ or 𝒢_2_, reduce to the dissipation rates 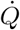from Eq. (23) on the respective graphs. The values 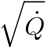are shown for the two irreducible flows and for the mass-action flow *f*_14∪193_, in Fig. 13. Flows along the base of the triangle are all those produced by adding to the mass-action flow multiples *v*° of the TKL trefoil. These pass through f14 and f193 one unit apart in *v*°, and 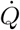 is parabolic in *v*° about *f*_14∪193_. Currents and 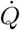 values are given in Table VII for the transect along the 235 TKL trefoil between f14 and f193, and also along a second transect in the AlKe null cycle connecting f193 to f184 in Table III, shown in the third panel of Fig. (7). The near-equality of the kinetic impedance presented for AlKe conversion through the C_4_ and C_5_ edges e6 and e7 causes the mass-action solution to split net flux nearly symmetrically through the two irreducible paths. This null cycle is of interest as a case of stoichiometric coupling between two potential branches, by the construction from Fig. 8.

**TABLE VII:** Flow properties of the Pythagorean theorem illustrated in Fig. 13. Edges in the graph of Fig. 2 are listed in the first column, and fluxes *v* at through current *J*^ext^ = 𝕊*v* are given in columns for each flow solution. 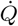 in the final row is multiplied by [GAP]^2^ in the thermochemical background where the force-flux relation is solved. The trefoil current *v*^*°*^ (the only null flow in the union graph) equals the flux in edge e23. At the minimum-dissipation solution 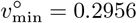. The relation (86) is fulfilled with the dissipation in the final row 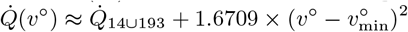.

| edge | f14 | f14∪193 | f193 | f193∪184 | f184 |
| --- | --- | --- | --- | --- | --- |
| e5 | 2 | 2 | 2 | 2 | 2 |
| e1 | 1 | 1 | 1 | 1 | 1 |
| e69 | 1 | 1 | 1 | 1 | 1 |
| e40 | 1 | 1 | 1 | 1 | 1 |
| e182 | 1 | 1 | 1 | 1 | 1 |
| e22 | 1 | 1 | 1 | 1 | 1 |
| e96 | 1 | 0.7044 | 0 | 0 | 0 |
| e23 | 0 | 0.2956 | 1 | 1 | 1 |
| e97 | 0 | 0.2956 | 1 | 0.5000 | 0 |
| e6 | 1 | 1 | 1 | 0.5000 | 0 |
| e7 | 0 | 0 | 0 | 0.5000 | 1 |
| e15 | 0 | 0 | 0 | 0.5000 | 1 |
| $\dot{Q}$ | 1.7796 | 1.6336 | 2.4626 | 2.1739 | 2.4627 |

**FIG. 13:**
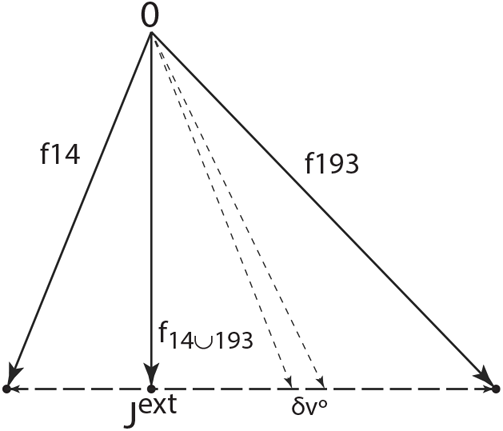
The Pythagorean theorem (86) in the linear-response regime where it is the ordinary Euclidean theorem. Two irreducible flows f14 and f193 differ by a TKL trefoil current *v*^*°*^ with magnitude 1. In the hierarchy 𝒢_2_ ⊂ 𝒢_1_ ⊂𝒢, 𝒢 is the full graph from Fig. 2, and 𝒢_1_ is the union of the supporting graphs for f14 and f193. 𝒢_2_ may be the supporting graph for either f14 or f193. Vertical solid arrow is square root of the integral 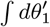 in Eq. (86) from zero to *θ*_1_(*J*^ext^) in 𝒢_1_. Hypotenuse solid arrows are square roots of integrals 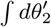 in Eq. (86) in either graph G_2_. Horizontal dashed arrows are square roots of integrals ∫ *dη*^*′*^ in either direction of *v*^*°*^ in Eq. (86). Details of the final flow parameters and the functional dependence of 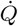 on *v*^*°*^ are given in Table VII.

#### b. Bottleneck reactions as organizers of flow redundancy and chemical work transduction

Highly evolved biological pathways often have large linear or directed acyclic regions, with redundancy eliminated unless it is under allosteric or other kinetic control [112, 113]. In such pathways the notion of the *rate-limiting step* is important because it can imbue long reaction sequences with the Arrhenius thermal response characteristic of single reactions, and simplify analysis to near-equilibrium conditions for other reactions. In putatively more primitive networks with significant redundancy through null cycles, the corresponding role is played by what we term *bottleneck reactions*, which partition flows into subgraph components, and become focal points for transduction of chemical work.

In the spatial network layouts shown in Fig. 7, the inputs GAP (*A*_3_) and F6P (*K*_6_) are delivered from the prelude at the “back face” of the cuboid, and the output Ru5P (*K*_5_) is exported from the front face. Because of the Möbius connection of the possible backbone cycles shown in the second panel, the edge e96 stands as a bottleneck between inputs and outputs.

The subgraph in the second panel of Fig. 7 has null dimension 2, from the 234 TKL trefoil passing through e96 from the left and the 235 TKL trefoil passing through e96 from the right. Table VIII lists values of currents *v* and chemical-potential drops Δ*µ* across the edges in the TKL trefoils in mass-action solutions on a sequence of nested graphs, starting from irreducible f14, adding e98 to produce the supporting graph for reducible f535, and then adding e23 and e97 to produce the union supporting graph f535 ∪ f193. The flux through e96 consists of three contributions: a through-flow of magnitude *v*_6_ = 1 that is constrained by the prelude topology, -*v*_98_ from the 234 trefoil, and *v*_23_ from the 235 trefoil. Currents, potential drops, and 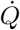 values are listed for two thermochemical landscape explained in App. E, corresponding to models with or without stereochemical contributions to molecule Δ_*f*_ *G*^0′^s.

**TABLE VIII:** Currents *v* and potential drops Δ*µ* on the edges in TKL trefoils for the masks of f14, f535, and f535∪f193. f535 adds one (234) TKL trefoil to f14 by adding edge e98, and f193 adds a second (235) TKL trefoil by adding edges e23 and e97. The final columns, f535∪f193, provide the labeling for the transduction in Fig. 14 in either potential. (eQNS) labels the thermochemical background for group contribution from non-stereochemical SMILES, and (eQS) labels the background with stereochemistry, approximating KEGG data values. The relation *v*_22_ − *v*_98_ = 1 = *v*_6_ in all cases is the topologically constrained AlKe velocity E4P *⇀* Eu4P. Chemical-potential cycles must cancel around trefoils; thus Δ*µ*_23_ + Δ*µ*_97_ − Δ*µ*_96_ = 0 and Δ*µ*_22_ + Δ*µ*_98_ − Δ*µ*_96_ = 0.

| | $f_{14}^{(eQNS)}$ | $f_{535}^{(eQNS)}$ | $f_{535 \cup 193}^{(eQNS)}$ | $f_{14}^{(eQS)}$ | $f_{535}^{(eQS)}$ | $f_{535 \cup 193}^{(eQS)}$ |
| --- | --- | --- | --- | --- | --- | --- |
| $v_{22}$ | 1.0000 | 0.9609 | 0.8553 | 1.0000 | 0.3919 | 0.3645 |
| $\Delta\mu_{22}$ | 0.5559 | 0.5341 | 0.4755 | 1.6440 | 0.6443 | 0.5992 |
| $v_{98}$ | | -0.0391 | -0.1447 | | -0.6081 | -0.6355 |
| $\Delta\mu_{98}$ | | -0.0209 | -0.0772 | | -0.3395 | -0.3548 |
| $v_{96}$ | 1.0000 | 1.0391 | 0.8063 | 1.0000 | 1.6081 | 1.2891 |
| $\Delta\mu_{96}$ | 0.4939 | 0.5133 | 0.3983 | 0.1896 | 0.3049 | 0.2444 |
| $v_{23}$ | | | 0.3384 | | | 0.3465 |
| $\Delta\mu_{23}$ | | | 0.2032 | | | 0.1824 |
| $v_{97}$ | | | 0.3384 | | | 0.3465 |
| $\Delta\mu_{97}$ | | | 0.1951 | | | 0.0620 |
| $\dot{Q}$ | 1.7796 | 1.7772 | 1.6035 | 1.9743 | 1.0898 | 0.9842 |

#### c. Upstream and downstream trefoils: parallel Kirchhoff transport before or after a bottleneck

The 234 trefoil lies upstream (on the input side) of the bottleneck edge e96, while the 235 trefoil lies downstream and provides two parallel branches to the outputs from the supports of f535 and f193. While addition of both trefoils contributes to reducing 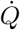 in Table VIII, they do so through different impacts on chemical potential and current profiles.

When e98 is added, the potential drop Δ*µ*_22_ is decreased as the 234 trefoil routes a part of the current *v*_22_ that is 1 in f14, around edge e22 and through −*v*_98_. The result is an *increase* in both the potential drop across, and the current through e96, and hence of the rate of chemical work delivered to its input complexes.

When e23 and e97 from f193 are added *parallel to* e96, activating the downstream 235 trefoil, the potential Δ*µ*_96_ and current *v*_96_ are both diminished, as e96 has been alleviated as a bottleneck. The net impedance of the left-hand subgraph from the input complexes of e96 is reduced, so flux *v*_98_ through the 234 trefoil is further increased, while the potential drop Δ*µ*_22_ and current *v*_22_ are further decreased.

### D. Application to transduction of chemical work

We may use these solution series to illustrate the non-stoichiometric or stoichiometric transduction of chemical work defined in Sec. II D 6, in particular as null flows allocate current along parallel pathways to redistribute chemical potential to minimum-dissipation values.

#### 1. The non-stoichiometrically-coupled TKL trefoils

The network in the second panel of Fig. 7 regarded as a graph 𝒢, with the sum of its two TKL currents defining a null flow *v*, and the edge e96 with dim_*v*_(e96) = 2, furnish an instance of the chemical work transduction through non-stoichiometric coupling introduced in Sec. II D 6. We take 𝒢′ ≡ 𝒢−*e*22−*e*97 to be the subgraph through which *v* flows, and with transduction from chemical work input across the boundary complexes of e22 and output across the boundary complexes of e97. Fig. 14 shows the graph 𝒢 (upper panel) and subgraph 𝒢′ (lower panel), respec-tively closed and open with respect to the two-trefoil null flow, with current components *v* and potential drops Δ*µ* marked from the solution 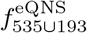 in Table. VIII.

**FIG. 14:**
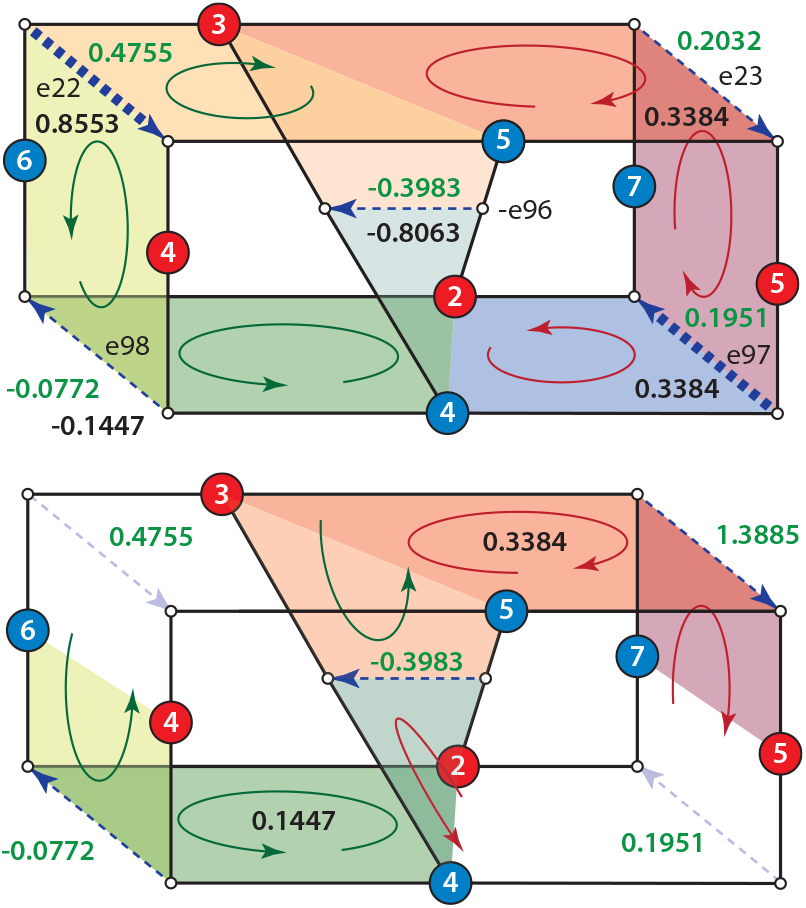
Two linked TKL-trefoils that appear in the union of the supporting graphs for two flows (f193 and f535). The Möbius boundary of each trefoil is formed by tracing a continuous path of solid (stoichiometric) links as they pass through species and complexes. A set of chemical-potential drops (green lettering) and currents (black lettering) for a 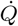-minimizing solution are shown for each reaction. Upper panel shows two basis elements for null flows on the complete supporting graph: the 234 trefoil (green shades) on the left and the 235 trefoil (red shades) on the right. The period-2 backbone cycles of Fig. 24 appear as simple circulations in their respective faces of the cube. Lower panel shows “environment” reactions *K*_6_ + *A*_3_ *⇀ A*_4_ + *K*_5_ and *A*_5_ + *K*_4_ *⇀ K*_7_ + *A*_2_ removed as explicit sources of dissipation. A red cycle with current *j*_2_ = 0.3384 fully accounts for the current supplied to the complexes bounding the “output” conversion *A*_5_ + *K*_4_ *⇀ K*_7_ + *A*_2_. A green cycle with current *j*_1_ = 0.1447 flows through the subgraph, alongside a supply of the boundary complexes *K*_6_ + *A*_3_ *⇀ A*_4_ + *K*_5_ at rate 0.8553 across the potential 0.4755. from the complement to this subgraph in the complete graph. Other sources (TAL edge and preludes) also couple to the boundary complexes, and serve to maintain their chemical potentials.

##### a. The “slip” of non-stoichiometric coupling and transduction loss

We noted in Sec. II D 6 that maximal transduction efficiency in non-stoichiometric coupling occurs if the transducing reaction presents infinite impedance to flow, so that all current is redirected “around” that reaction, directly transmitting all chemical potential from inputs to outputs. Any current through the coupling reaction is a source of dissipation and thus of inefficiency.

As noted in Sec. IV C 3, the current through e96 is composed of three parts: *v*_96_ = *v*_6_ − *v*_98_ − *v*_23_. The first two (*v*_6_, -*v*_98_), are inputs respectively from the topologically-dictated through-flow and from the 234 trefoil that we are taking to be driven by the potential drop across the boundary complexes of e22. The third (*v*_23_) is the induced flow that we take to be driven by the potential drop across e96 and delivering chemical work output to the boundary complexes of e97.

Relative to f14 as a reference (which lacks *v*_98_), the 234 trefoil is active with current *j*_1_ = 0.1447, and the 235 trefoil is active with current *j*_2_ = 0.3384. The former (green arrows in the figure) takes chemical work from the boundary complexes of edge e22, and the latter (red arrows) delivers chemical work to the boundary complexes of e97. They are coupled through the potential drop across e96. The current *j*_1_ = 0.1447 delivers chemical work to the boundary complexes of e96 and contributes to maintaining its chemical potential drop in the forward direction. The current *j*_2_ = 0.3384 cancels part of the dissipation by *j*_1_, routing it instead as chemical work to the complexes bounding the remainder of the 235 trefoil. Part of that work is dissipated in e23 (top right edge), while the rest is delivered as chemical work to the output complexes that are the boundary of e97.

We note that *j*_2_ *> j*_1_: the total work transported away from the boundary of e96 by *j*_2_ is larger than the part delivered by *j*_1_. This is because the net current through G, performing the overall conversion (34), also flows through e96 and contributes to maintaining the potential drop Δ*µ*_96_. The total currents in Table VIII always satisfy *v*_6_ − *v*_98_ ≥ *v*_23_. The difference is the “slip” current from non-stoichiometric coupling that creates inefficiency.

##### b. Impacts on whole-network throughput and dissipation

Table X embeds the foregoing account of chemical work transduction between the two TKL trefoils as part of the larger current re-arrangement by activation of first upstream and then downstream trefoils, for the three nested flow solutions and two thermochemical equilibrium landscapes shown in Table VIII.

**TABLE IX:** Potential drop Δ*µ*_96_ and total current (*v*_96_ + *v*_23_) across the input and output complexes to e96, delivered y the graph complement to the 235 trefoil (e23, e97, - e96) in the whole graph. Total chemical work 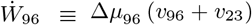 delivered to these boundary complexes is the product. The chemical work is divided between an amount Δ*µ*_96_*v*_96_ dissipated within e96, and a remainder Δ*µ*_96_*v*_23_ = (Δ*µ*_23_ + Δ*µ*_97_) *v*_23_ delivered to the subgraph formed by e23 and e97. Those may be considered in any combination, internal dissipation through edges retained “within” a transduction graph, and chemical work delivered to complexes that are the boundary of the graph, as shown in the lower panel of Fig. 14. The fraction of chemical work delivered to this boundary, out of chemical work delivered to the overall graph (measures as 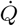) also shown.

| | $f_{14}^{(\text{eQNS})}$ | $f_{535}^{(\text{eQNS})}$ | $f_{535 \cup 193}^{(\text{eQNS})}$ | $f_{14}^{(\text{eQS})}$ | $f_{535}^{(\text{eQS})}$ | $f_{535 \cup 193}^{(\text{eQS})}$ |
| --- | --- | --- | --- | --- | --- | --- |
| $\Delta\mu_{96}$ | 0.4939 | 0.5133 | 0.3983 | 0.1896 | 0.3049 | 0.2444 |
| $(v_{96} + v_{23})$ | 1.0000 | 1.0391 | 1.1447 | 1.0000 | 1.6081 | 1.6355 |
| $\dot{W}_{96}$ | 0.4939 | 0.5333 | 0.4559 | 0.1896 | 0.4902 | 0.3997 |
| $\dot{Q}$ | 1.7796 | 1.7772 | 1.6035 | 1.9743 | 1.0898 | 0.9842 |
| $\dot{W}_{96}/\dot{Q}$ | 0.2776 | 0.3001 | 0.2843 | 0.0960 | 0.4498 | 0.4061 |

**TABLE X:** Summary of flow and subgraph counts from the ILP enumeration of flow solutions to conversion (34) used in the text.

| Description | count |
| --- | --- |
| stoichiometrically distinct flows $f_i$ | 763 |
| stoichiometrically distinct reducible flows $f_i$ | 85 |
| stoichiometrically distinct irreducible flows | 678 |
| distinct supporting graphs $\text{supp } f_i$ | 744 |
| distinct $\text{supp } f_i$ from reducible flows | 66 |
| distinct $\text{supp } f_i$ with “superfluous edges” | 6 |
| distinct dissipation-minimizing flows | 738 |
| distinct reducible dissipation-minimizing flows | 60 |

**TABLE XI:** Designations and properties of species used in the network models. First column gives C count, aldose/ketose/bisphosphate grouping, or other designations. Second column is the MØD index in the simulation. Third column is the pubchem name returned by eQuilibrator 3.0 in a search on the SMILES string (all sugars are (aq), not written) Fourth column is a shortened tag label. Fifth column Δ_*f*_ 𝒢^*′*0 (Li)^ is standard-state free energy of formation from [135]. Δ_*f*_ 𝒢^*′*0 (eQNS)^ is from eQuilibrator 3.0’s group-contribution estimator from non-stereochemical SMILES. Δ_*f*_ 𝒢^*′*0 (eQS)^ is eQuilibrator 3.0’s default estimate computed by stereochemical group contribution and benchmarked against KEGG [62, 87]. Δ_*f*_ 𝒢^*′m* (eQS)^ is the value from the fifth column shifted in eQuilibrator to physiological conditions. Stereochemical SMILES distinguishes two C5 ketose-monophosphates (ribulose and xylulose) and two C6 ketose monophosphates (D-fructose and *β*-D-fructose). Since this network uses non-stereochemical SMILES, the nodes are merged. The free energy value used here in examples for C5 is xylulose, and for C6 is *β*-D-fructose, to more nearly match empirical networks. eQuilibrator entries that are duplicates in eQNS are greyed out. The chosen eQS entries are closer to the eQNS values in all cases (within about 3 kJ/mol). eQuilibrator parameters are set to pMg = 3.1 (to match the 0.8 mM values in [135]), and ionic strength = 0.25 M. Red assignments are those without eQS eQuilibrator entries, generated by a second-order polynomial fit for aldose-monophosphates, and a first-order (linear regression) for ketose-monophosphates and bisphosphates, to the free energies of formation with carbon number.

| Cn-A/K/B | MØD | PubChem name | label | $\Delta_f G'^0(\text{Li})$ | $\Delta_f G'^0(\text{eQNS})$ | $\Delta_f G'^0(\text{eQS})$ | $\Delta_f G'^m(\text{eQS})$ |
| --- | --- | --- | --- | --- | --- | --- | --- |
| H<sub>2</sub> |  | H2 (dissolved) | H2 |  |  | 99.1 | 82.0 |
| CO<sub>2</sub> |  | CO2 (dissolved) | CO2 |  |  | -386.0 | -403.1 |
| H<sub>2</sub>O(l) | 47 | water (liquid) | H2O |  | -156.65 | -156.6 | -156.6 |
| HPO<sub>4</sub><sup>2-</sup> | 0 | hydrogen phosphate | PO4 |  | -1058.55 | -1058.5 | -1075.7 |
| C2 aldose-P | 27 | glycolaldehyde-phosphate | GLP |  | -1019.97 | -1026.4 |  |
| C3 aldose-P | 2 | glyceraldehyde-3-phosphate | GAP | -1187.64 | -1091.91 | -1088.8 | -1105.9 |
| C4 aldose-P | 4 | erythrose-4-phosphate | E4P | -1306.62 | -1163.80 | -1159.0 | -1176.1 |
| C5 aldose-P | 5 | ribose-5-phosphate | R5P | -1435.72 | -1235.65 | -1234.7 | -1251.8 |
| C6 aldose-P | 23 | glucose-6-phosphate | G6P | -1567.08 | -1307.49 | -1319.0 | -1304.7 |
| C5 bis-P | 66 | ribulose-1,5-bisphosphate | Ru15BP |  | -2130.55 | -2132.7 | -2149.8 |
| C6 bis-P |  | fructose-1,6-bisphosphate | F16BP | -2401.08 | -2202.40 | -2205.4 | -2222.5 |
| C6 bis-P | 9 | β-D-fructose-1,6-bisphosphate | bF16BP | | -2202.40 | -2205.0 | -2222.1 |
| C7 bis-P | 13 | sedoheptulose-1,7-bisphosphate | S17BP |  | -2274.24 | -2274.2 | -2291.4 |
| C8 bis-P | 15 | D-glycero-D-altro-octulose-1,8-bisphosphate | O18BP |  | -2346.09 | -2345.5 |  |
| C3 ketose-P | 3 | dihydroxyacetone-phosphate | DHAP | -1194.65 | -1094.38 | -1094.4 | -1111.5 |
| C4 ketose-P | 18 | erythrulose-4-phosphate | Eu4P |  | -1165.76 | -1165.8 | -1182.9 |
| C5 ketose-P |  | ribulose-5-phosphate | Ru5P | -1434.72 | -1237.64 | -1232.6 | -1249.7 |
| C5 ketose-P | 6 | xylulose-5-phosphate | X5P | -1436.00 | -1237.64 | -1236.0 | -1253.1 |
| C6 ketose-P |  | fructose-6-phosphate | F6P | -1563.96 | -1309.49 | -1316.3 | -1333.4 |
| C6 ketose-P | 7 | β-D-fructose-6-phosphate | bF6P | -1563.96 | -1309.49 | -1312.0 | -1329.1 |
| C7 ketose-P | 8 | sedoheptulose-7-phosphate | S7P | -1685.66 | -1381.34 | -1385.7 | -1402.8 |
| C8 ketose-P | 34 | D-glycero-D-altro-octulose-8-phosphate | O8P |  | -1453.18 | -1457.4 |  |

e96 and its complement (e23 + e97) within the supporting graph of the 235 TKL trefoil are parallel pathways for current from the same boundary complexes. The total current delivered to those complexes can be expressed as either of the two sums (*v*_96_ + *v*_23_) = (*v*_6_ − *v*_98_). From this current and the potential drop Δ*µ*_96_, the chemical work 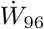 delivered to this boundary by the graph-complement of the 235 trefoil is computed.

The second row of Table X shows that the current delivered to the boundary of e96 increases with the activation of both upstream and downstream trefoils. However, as noted in Sec. IV C 3, activation of the 234 trefoil upstream of e96 increases Δ*µ*_96_ and overall work delivered, whereas activation of the 235 trefoil reduces Δ*µ*_96_ enough to reduce work 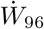 delivered despite the increase of total current.

The total dissipation 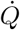 in this graph 𝒢′, carried forward from Table VIII, decreases as each added edge relaxes constraints for the throughput solution. The ratio of work delivered to the boundary of e96, relative to total work delivered to the entire graph, is shown in the table as 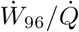. That ratio also is not monotonic, suggesting that the edge e22 is a greater bottleneck than e96 for the whole conversion.

#### 2. The stoichiometrically-coupled AlKe null cycle

Fig. 15 shows the AlKe null cycle from Fig. 8, embedded in the subgraph 𝒢′ from Fig. 7, with potential drops for the solution *f*_193∪184_ of Table VII, in the thermochemical background labeled eQNS from App. E. The flow f193 uses the 235 TKL trefoil to route species conversions through the cycle (dark red arrow in the figure) between *A*_5_ and *K*_7_ and the AlKe edge e6, with current *v*_23_ = *v*_97_ = *v*_6_ = 1, eliminating the septum through e96 in Fig. 14. The flow f184 provides a parallel route through the TAL edge e15 and the AlKe edge e7, in place of the TKL edge e97 and AlKe edge e6. The near-equality of TAL and TKL conductances and of all AlKe conductances in this thermochemical background results in a flow on the union graph that almost equally divides current through these two parallel conduction pathways, reducing by half their contribution to total-graph resistance, in the manner of parallel resistors in an electric circuit.

**FIG. 15:**
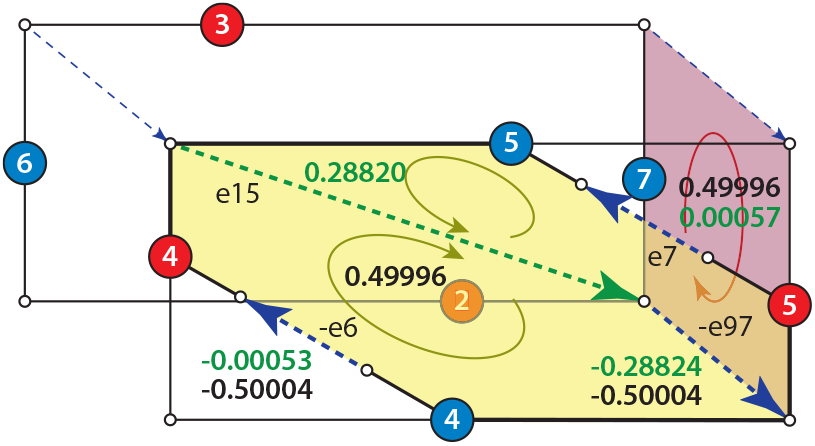
The chemical potentials in the solution on the union of supports for f193 and f184, across all edges of the AlKe null cycle. Current through the null cycle is shown in black; potential drops in green. The potential drop across the AlKe edges is ∼ [GAP] ∼ 10^*−*3^ smaller because they are first-order in organics, whereas the TAL and TKL edges are second-order. Therefore almost-all chemical work delivered by the current *v*_23_ = 1 (dark red circle) to the boundary of e97, which is saved from dissipation in e97 by the flow around the AlKe null cycle, is transduced stoichiometrically to the boundary of e15, where it is dissipated.

##### a. Efficient work transduction through stoichiometric coupling by AlKe edges with 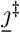

We noted in Sec. II D 6 that, because all flows in an irreducible null cycle have fixed proportions, the efficiency of stoichiometric transduction does not depend on the magnitude of the current through the coupling graph, but only on the ratios of chemical potential drops Δ*µ*. If in Fig. 15 we regard the pair of AlKe links as a coupling graph, because they are monomolecular their equilibrium currents *j*^‡^ are larger by a factor of 1*/* [GAP] ∼ 10^3^ than those of TAL or TKL edges, resulting in Δ*µ*_6_ ∼ Δ*µ*_7_ ∼ 10^−3^Δ*µ*_97_. The AlKe null cycle, in canceling ∼ 1*/*2 the current through e97 and transferring it to e96 – and because there can be no “slip” between the stoichiometrically coupled currents *v*_15_ and *v*_−97_ – delivers nearly all the chemical work conserved at the boundary of e97 to the boundary of e15 where it is dissipated.

### E. The dissipation functionals in geometric coordinates

#### 1. Why geometric coordinates?

The fundamental result we have been seeking is a natural and interpretable way to decompose the large-deviation measure (46,48) of more specific flow pathways through a network relative to less-specific ones, which can be carried out in parallel to the Kirchhoff decomposition for flows that exists as a straightforward exercise in the linear algebra of null spaces (19).

Part of the need for a natural probability decomposition actually stems from the problem of Kirchhoff decomposition itself, particularly as it arises for stoichiometric systems. For simple superposable electric currents in the linear regime of RLC circuits [56], an ordinary Euclidean inner product on current components gives a natural and unambiguous notion of maximal independence among basis elements. Stoichiometric flows, in contrast, are “coupled cycles”, on superpositions of which multiple nonlinear complex activities determining the mass-action rate law can depend simultaneously. Decomposition of dissipation functions can provide natural notions of orthogo-nality for independent flow components.

A formulation of maximal independence of additive components of probability distributions was given by Chentsov [111] in terms of KL divergences from exponential families, connecting to the local metric distance on probability distributions from Fisher [103] and Rao [114], and extended it to the unusual construct of dual connections that became the foundation of information geometry [60, 61]. We have demonstrated in Sec. IV the parallel forms (58) for the the LDF for counts in a single-time distribution using the Hessian of the CGF *ψ*, and the integrand (74,75) of the LDF for steady-state currents using the Hessian of the generator 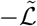 that is the Hamiltonian of the corresponding Hamilton-Jacobi theory.

We then showed in Equations (84,85) that the extended Pythagorean theorem from information geometry constructs the space of null flows interior to a sub-graph as a natural exponential family with coordinates *η* about a base distribution which is the mass-action solution with source *J*^ext^; and that that solution is the “nearest” solution in the exponential family to the background in the subgraph without an external source. (That is, in the language of information geometry, the family of mass-action solutions from the source-free steady state to driven steady states with sources *J*^ext^ is a *mixture family* in the current coordinate *J*^ext^.) Following on the observation [46, 115] that relinquishing the time-reversal symmetry that defines equilibrium introduces entropy functions in which currents as well as counts become the extensive state variables, we see that both mixture and exponential families are defined here in terms of currents, as the convex potential *ψ* is replaced by 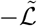, and the single-time potential is replaced by a dissipation rate in an integrand.

This section presents the expansion of the dissipation function, working at quadratic order to simplify, in terms of the eigenvectors of the Fisher metric obtained from 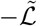, and shows how the current solution about a general equilibrium is defined by minimum dissipation subject to a vector of constraints reflecting the network reduction.

#### 2. Definition of the metric from Hessians of the CGF and LDF integrands

We begin with a construction of the Fisher metric for a general 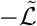, with any half-reaction rate parameters and any sources *η*. Let 𝒢 be a graph with stoichiometric matrix 𝕊 and let {*f*_*µ*_} be a basis for ker 𝕊. Each *f*_*µ*_ ≡[*f*_*µ* (*ji*)_]is a column vector on the reaction index (*ji*).

For any such basis we introduce a dual basis 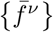 of projection operators, which are row vectors 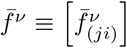 on the same reaction indices, defined so that a representation of the identity matrix on ker 𝕊 can be expanded as

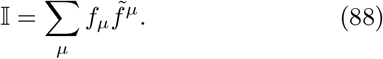

Eq. (88) is equivalent to requiring

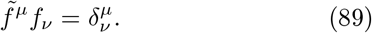

The choice of basis {*f*_*µ*_} is arbitrary, but if we wish to consider a subgraph 𝒢′ ⊂ 𝒢 and solutions *v* to 𝕊|𝒢′ *v* = *J*^ext^ for currents through the boundary of 𝒢′, it may be convenient to choose a basis in which ker S|(𝒢_′_)_⊥_ is carried by a subset of the basis elements, which will then not appear in sums for flows within 𝒢′.

The Hessian of the Hamiltonian 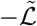 in Equations (74,75) defines an inner product between vectors in the *dual* basis,

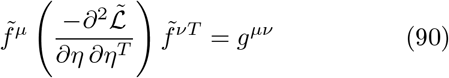

which are the elements of the matrix inverse *g*^−1^ of a metric tensor *g*, which is just the Fisher-Rao tensor [103, 104] for the dual geometry defined by current variations under sources *η* [60, 61].

We now expand the gauge-invariant expression for the source terms in the dual basis as

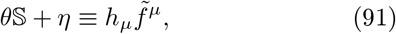

with differential

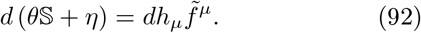

The components of *h* are affine coordinates on the space of flows written as an exponential family.

To obtain the dual coordinates on the flows themselves which make a mixture family, we first assume as in Eq. (76) that the Hessian 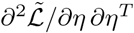 is invertible (all reactions have sources). We nest the two representations (88,76) of the identity on the space of flows within the inner product (89), to obtain

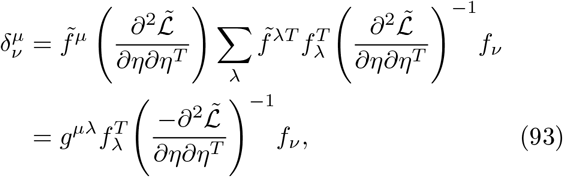

from which we identify matrix elements of the metric *g* on flows as

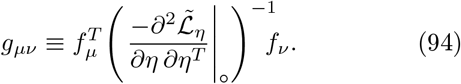

Current vectors *v* are assigned coordinates *α* ≡ [*α*^*µ*^] on the basis {*f*_*µ*_} as

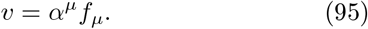

We denote by *α*_°_ the coordinates of any current vector *v*_°_ under the generator ℒ in the absence of sources.

##### a. Duality of exponential and mixture coordinate differentials through the Fisher metric

Because the CGF *ψ*_*T*_ is constructed as a function of arguments *η*, while the LDF *S*_*T*_ is constructed as a function of the dual arguments *v*, it is conventional to write the second variational derivatives of these potentials – including their evaluations in terms of 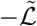 – as functions of these arguments. The counterpart to Eq. (71), for steady states and written in geometric coordinates, becomes

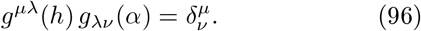

The relation (75) between differentials of *v* and *η*, when written in terms of the Fisher metric using the expansions (91,95), becomes by Eq. (96)

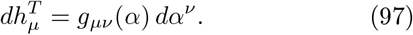

In terms of these the integrands in Eq. (82) become, in the dual geometry,

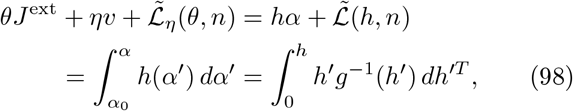

where the vector products mean (in the Einstein convention that paired upper and lower indices are summed):

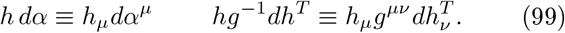

Eq. (74) for *S*_*T*_ correspondingly becomes

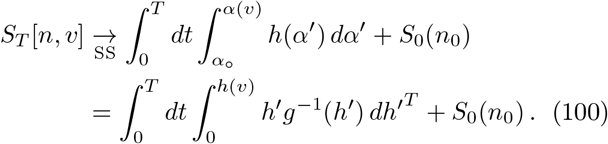

##### b. Extended Pythagorean decomposition in ge-ometric variables

The two components of the Pythagorean decomposition (85) are written in Eq. (100) as two legs in a contour of integration ∫*dh*′. If we write *<u>h</u>* for the source coordinate of the mass-action solution to 𝕊*v* = *J*^ext^ on 𝒢′, then information-geometric orthogonality of the two legs is expressed by the relation *h dα*′ = 0 on the second leg of the integral that activates null flows diverging from that mass-action solution.

#### 3. Linear-response near the untilted steady state

By studying the integrand in the LDF (100) in the regime of linear response about a steady state in the untilted generator ℒ, we may take the metric as constant at its value at the minimum of *S*_*T*_. The integrand to quadratic order equals the thermodynamic dissipation, and we may express any current solution as a minimum-dissipation solution subject to a linear constraint.

We begin by writing *α* ≡ [*α*_*µ*_] as a column vector, and arranging the column vectors {*f*_*µ*_} into a matrix *f* ≡ [*f*_*µ*_]^*T*^ with column index *µ*. Eq. (95) for the offset of *v* from the value *v*_°_ in the absence of sources then becomes as

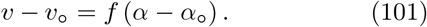

The Fisher metric from the generator 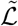 is evaluated only at its value without sources (*θ, η*), and we denote the metric and its inverse at the undriven steady state as

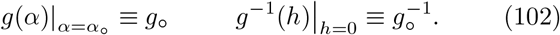

For an application to the sugar-phosphate example, we need only those basis functions that flow through the graph 𝒢′, and may choose the basis so that every entry satisfies

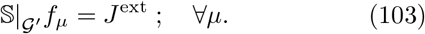

Then for any pair of distinct indices *µ* and *v, f*_*µ*_ − *f*_*v*_ is a null flow in 𝒢′. As a corollary, any solution *v* = *fα* with 𝕊| _𝒢′_ *v* = *J*^ext^ must then have 1^*T*^ *α* = 1.

The dissipation integrand (98) is evaluated only to quadratic order in sources, and becomes

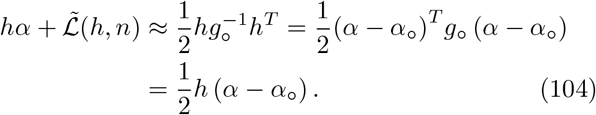

The Legendre dual condition (62) on *η*, which in the steady state incorporates the stationarity condition (66) for *θ*, takes the form from Eq. (104) of

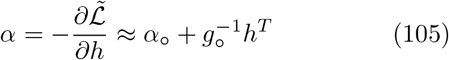

Performing a coordinate transformation from *h* to *α*−*α*_°_, Eq. (105) is equivalent to the dual condition

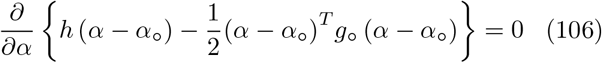

Eq. (106) is none other than the familiar condition that the Legendre transform is the functional form (44) evaluated at the minimizer of the extensive variable, which gives the inverse function (45) to the intensive variable relation (43), familiar from the construction of free energy from entropy in equilibrium thermodynamics but applied here to dissipation in the space of currents.

As illustrated in Fig. 16, the minimizer of the quadratic form (104), corresponding to the first line in the Pythagorean decomposition (85), is given when *h* ∝ 1.^28^ The normalization condition 1^*T*^ *α* = 1 then requires 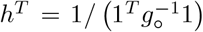 in Eq. (105). The coordinates *α* for the mass-action current, which we denote by *α*_∥_, are then given by

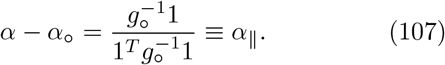

We will refer to the dissipation 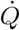 net of the contribution from any background currents *α*_°_ from the un-driven steady state as the *excess dissipation*, 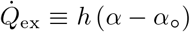. On the mass-action solution (107), it is given by

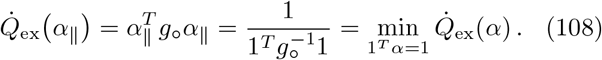

If the source *η* induces further null flows in 𝒢′, we designate these as *α*_⊥_. Then the total current and the value *h*^*T*^ from Eq. (105) become

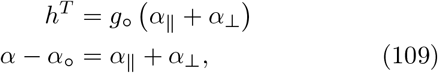

in which we must have 1^*T*^ *α*_⊥_ = 0. The integrand (104) and its relation to the excess dissipation then evaluate to

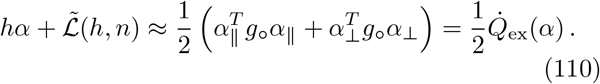

The Pythagorean decomposition (85) reduces in the linear response regime to the Euclidean triangle equality shown in Fig. 13, expressed in the metric *g*_°_. The steady-state LDF (100) accordingly becomes

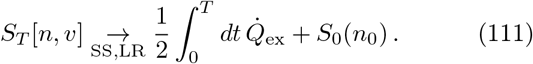

**Remark:** The LDF for steady-state currents is the integral of the thermodynamic heat-dissipation rate *only* in the regime of linear response. Thus the unlikelihood of forming a persistent current diverged from the mass-action solution is an inherently kinetic characteristic: it is not fully accounted-for by differences in rates of deposition (“production”) of equilibrium entropy in the thermochemical reservoir that defines boundary conditions for the system in terms of equilibrium state variables. In addition to recapitulating the functional form, the kinetic origin of these LDF integrands recapitulates the argument from Sec. IV A 5 that escape paths, and not energy deposition, capture the measures of cumulative improbability in information divergences.

**FIG. 16:**
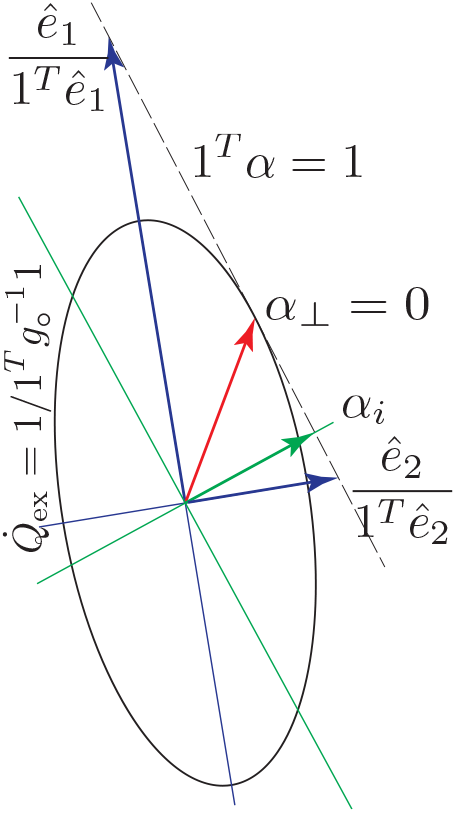
The stationarity condition (105), written in the dual coordinates of Eq. (106). Ellipse is a surface of constant excess dissipation 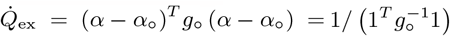 from Eq. (110) below. This is the minimum of 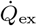 on the surface 1^*T*^ *α* = 1 of fixed flux *J*^ext^. *ê*_1_ and *ê*_2_ are the eigenvectors of *g*_*°*_, normalized here to the 1^*T*^ *α* = 1 surface. *α*_*i*_ is the coordinate vector of some other integer flow solution fi, restricted to support on a subgraph of the graph defining *g*_*°*_.

#### 4. Evaluation of the linear dissipation in terms of flow components

We may apply the Pythagorean decomposition (110) for the excess dissipation to the sugar-phosphate example, in the metric coordinates illustrated in Fig. 16. Since this is a detailed-balance example, *α*_°_ = 0 and the excess dissipation 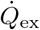 is just the total dissipation from Eq. (23). On any graph 𝒢′ with null cycles, the minimum-dissipation flow is shown as the red arrow labeled *α*_⊥_ = 0 in Fig. 16. Other integer flows correspond to instances marked *α*_*i*_ (green arrow) in the figure with *α*_⊥_ ≠0. As shown in App. G 1 e, the current in the LDF integrand (82) with the same species counts *n* as a steady-state mass-action solution has conjugate-momentum *θ*^*T*^ = log (*n/n*), the same value as the chemical potential *µ* in Sec. II D 8. The linear-response kernel for *θ* is then given by

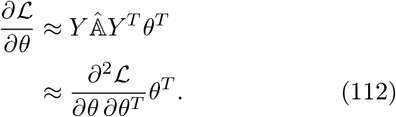

The stationarity condition (80) gives the connection to *J*^ext^, from which the dissipation becomes a quadratic form in *θ*,

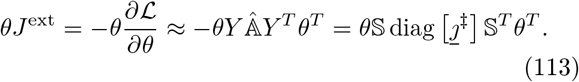

The current, in turn, is given by

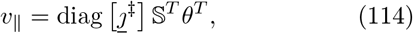

the opposite of the mass-action value (17), with likewise time-reversed source *J*^ext^ = 𝕊*v*_∥_ .^29^

The minimizing dissipation for that *J*^ext^ is given in both metric coordinates and reaction coordinates by

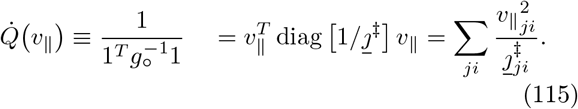

For any other solution such as 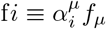 an integer-flow solution necessitating *α*_⊥ ≠_0 in Eq. (109), the excess dissipation evaluates to

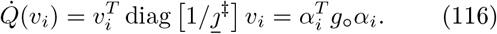

For the case *j*^‡^ ≡ 1 where 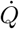 of Eq. (116) corresponds to what we have called the “topological” contribution to dissipation 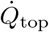 (33), we may compare these to 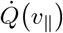 on the supporting graphs of the 763 integer-flow solutions from Fig. 4. For the irreducible flows, the two are necessarily the same. For reducible flows, the minimum-dissipation solution is a linear combination of the irreducible flows on that supporting graph. The informativeness of these topological measures about empirically grounded thermochemical equilibrium is assessed in Fig. 30 of App. E 1 a.

#### 5. Metric eigenvectors

The metric eigenvectors – principle axes of the ellipsoid in Fig. 16 – provide another way to assess the divergence of mass-action flows on complex networks from their reductions to one or another irreducible pathway flow. If one of the metric eigenvectors is closely aligned with the whole-network mass-action flow, then flows contributing to transverse eigenvectors may be eliminated with little increase in dissipation on the reduced network. If one or more integer flows are are close in the metric embedding to one of the eigenvectors, then network pruning to isolate that subset of flows can eliminate transverse directions without significantly increasing the dissipation relative to the unrestricted network. The degree to which the dissipations associated with eigenvectors of *g*_°_ approximate those of integer flows, with an evaluation of the specificity/dissipation trade-off resulting from extraction of the Calvin cycle, are given in App. E 2.

#### 6. A resistance measure for net conversions in stoichiometric networks in the linear-response regime

Just as we have described Kirchhoff decomposition of currents on stoichiometric networks as a generalization of its counterpart for simple electric circuits, we can define a measure of impedance for conservative flows with sources and sinks, which is the corresponding generalization of Ohmic resistance in circuits. From Eq. (116) for dissipation, we may define a resistance to the mass-action flow on any subgraph 𝒢′ as the ratio of excess dissipation to the squared magnitude of the through-current:

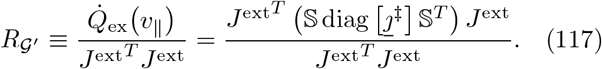

*R*_𝒢′_ is independent of the magnitude of the current at linear-response order, and depends only on the interaction of the structure of the through-current with the topology of the graph and the thermochemical equilibrium *j*^‡^ about which the response is induced. Eq. (117) is the stoichiometric counterpart to Ohmic resistance defined by the relation *R* = *V/I* = *P/I*^2^ for voltage *V*, current *I*, and power dissipation *P*, so we call *R*_*𝒢*_′ a conversion resistance.

To compare resistance of a flow through multiple nested graphs 𝒢_2_ ⊂ 𝒢_1_ ⊂ 𝒢, we take 𝒢_1_ to define geometric coordinates *α*_∥_ in Eq. (110), and 𝒢_2_ to define *α* = *α*_∥_ + *α*_⊥_, in terms of which

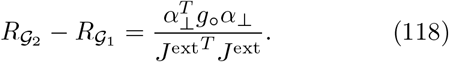

### F. The biological questions motivating a study of information

#### 1. The information distinguishing a pathway within a network derived from rules

We chose sugar-phosphate chemistry to illustrate principles and strategy of rule-based modeling because it provides two combinatorial diversifications to be understood: from few rules come indefinitely large and networked stoichiometric graphs; and from the usual many-particle limits of thermodynamics the reactions in these graphs generate macroscopic state and transport behaviors. A highly combinatorial reaction network from few chemical mechanisms isolates an interesting problem in the evolution of pathway specificity and control, which is likely present for most biological pathways but nowhere else so clearly isolated as a combinatorial problem.

Many enzyme families are organized [54, 55] around a retained mechanism and catalytic site under strongly conservative selection, with sub-families diverging in substrate specificity or other modes of subfunctionalization. For reactions that form networks by many serendipitous combinations [116, 117], the asymmetry of conservation between mechanism and substrate discrimination in enzyme families suggests that the earliest primitive enzymes, by opening reaction mechanisms but not yet selecting precise substrates, would have brought large combinatorial networks into existence. Only through later selection for specificity – in part made possible by the productivity of earlier diffuse networks – would reactions be eliminated until only precise pathways remained.

Selection is understood as imparting adaptive information in populations, at some cost in organisms and the embodied resources to form them. What is increasingly appreciated is that a natural measure of the information imparted can be the same as the cost function measured in logarithmic units of population growth [118–121].

Here we wish to identify lower bounds on the information required to evolve primitive enzymes, and the degenerate networks they produce, into specific enzymes catalyzing networks assembled from precise pathways. This information will not generally be localized in single reactions, as it depends on substrate availability which is jointly determined across the network. An absolute lower bound on the information in specificity should not be rooted in the molecular biology of particular catalysts, though the latter could impose tighter bounds requiring more information than the abstract lower limit. We will propose here that a lower bound is given from the information divergence *S*_*T*_ of Eq. (82) and its decompositions (84,85), which assign a likelihood cost to the sampling bias needed within the underlying permissive network to produce the same current pattern as the one from removal of reactions by enzyme specificity.

#### 2. The information divergence as a direct measure of evolutionary cost

Our proposal to use the LDF for currents as a cost measure to tie physiological events to selective events is driven by two considerations: First, relative entropies [122–124] and a variety of functions either derived from [125, 126] or related to [127] them are understood as measures of the both information and cost [31, 118, 120, 121, 124] through selection in population processes. Second, the events within physiology form a nested partition function with the population-level events of reproduction or death through which selection acts in the form of differential rates.

The overall LDF for such a nested process may be decomposable (for example by a chain rule for relative entropies) into summands of which the network divergence *S*_*T*_ of Eq. (82) characterizes the likelihood for events within physiology (chemical conversions) by alternate pathways, while another summand characterizes likelihoods through Malthusian selection events that alter population composition [29–31] by replacing more open with more restrictive networks to produce altered current routes through constraints. The robust population trajectory must be one that is jointly maximal in the product of these contributions to probability (inevitably along with many others), giving the implication that *ceteris paribus*, the improbability-cost of forcing current through a restricted subnetwork must not accumulate to a larger value than whatever probability gain is achieved at the population level, as reflected in selection to maintain the more evolved and constrained network.

We have not aimed in this paper to construct a model with both selection and physiology, which must be a separate project. We aim in this section only to justify large-deviation log-likelihood as a common denomination in which costs-per-event in chemical kinetics and in biological lifecycles can be quantitatively compared.

**Remark:** A naïve story to bolster the intuition for this argument may be made from the relation (111) of the LDF and the integrated dissipation in the linear-response regime: network restriction increases the chemical work dissipated per unit of some chemical conversion performed, such as PPP catabolism or Calvin-cycle anabolism, which is essential to an organism’s viability. Excess work lost to dissipation should appear as a fitness cost in some other essential function, favoring lower-dissipation networks. For selection to maintain the more specific but more dissipative network, it must yield some other advantage – elimination of substrate loss to diffusion or side-reactions, toxicity, *etc*. – which favors the specific phenotype even at a higher throughput cost in dissipated work.

## V. CONCLUSIONS: CONSEQUENCES AND FUTURE DIRECTIONS

Our purpose in this paper has been to introduce rule-based modeling for chemistry in a framework of relations that we call “three-level systems”, and to illustrate the interactive and complementary use of rules, stoichiometric graphs, and probability dynamics to recognize and characterize the loci of causation in such systems.

The main emphasis in the three-level framework is that each level has a generative relation to the one following, with the consequence that constraints, patterns, or information imposed at smaller simpler levels can propagate through the generative relations to govern patterns and dynamics at later levels even when those are combinatorially larger than their progenitors. The possibility for pattern preservation across indefinite ranges of scale is the source of the concept of macrostates [32, 43, 45], and the origin of thermodynamics from stochastic processes. Here we include that relation and augment it with the earlier generative relation that rule systems hold over the process models that become the generators of state and probability dynamics.

We developed some formal methods at two levels: from graph theory for the decomposition of graphs, currents on and through them, and work transduction and efficiency following [1]; and from large-deviation theory for currents as a tool to assign probabilistic costs to topological modifications of networks, by expressing mass-action flows on nested graphs in terms of the sampling bias required to produce more restricted flows from less restricted ones.

We have used the context of evolution as a guide to the kinds of analysis that are desirable for the combinatorial chemical systems that three-level frameworks are well suited to study. These include general problems of understanding how topology governs the decomposition of flows and how flow decomposition and large-deviation probability decomposition relate, and also more specific problems such as identification of shortest paths, paths of least resistance, or particular instances of transduction of chemical work and redistribution of chemical potential within networks and between networks and their environments.

Evolutionary control naturally crosses levels in real molecular systems, from reaction mechanisms to substrate specificities to kinetic regulation, and the three-level framework gives some explanation of how coordination of these under selection can be possible: global properties of complexity or minimality of reaction currents can be governed at the mechanism level and derived directly from rule properties without the insertion of costly and complex steps such as exhaustive enumeration and search of large combinatorial spaces, which selection may have limited capability to solve [128].

We have introduced specific methods to solve particular problems pertinent to chemistry but also to more general stochastic stoichiometric population processes. One of these is a lattice-graph representation that expresses pathways as simple closed curves rather than integer flow solutions on stoichiometric graphs, and thereby allows the expression of rule symmetries and conservation laws. An interesting result from this construction is that the Calvin cycle is one of the two unique shortest paths for the net conversion it performs. The absence of thermochemical data on 2-phosphoglycolate may suggest that this compound is not readily formed or is unstable, thereby eliminating from possibility the only other comparably short path to the Calvin cycle.

A second result is the demonstration that the natural *additive* decomposition of large-deviation functions for currents on nested subgraphs is an instance of the familiar extended Pythagorean theorem of information geometry, identifying the natural dual geometry that follows from the construction of the LDF as an information divergence. An interesting small feature of the latter construction is the analogous role played by the Liouville function in the LDF for currents, to the role played by the CGF in the construction for distributions of states at a single time.

Finally, we have suggested that large-deviation probabilities – precisely because they nest in the manner of partition functions for multilevel systems, and because they can be additively decomposed on nested networks – provide a natural measure of evolutionary cost from events at the physiological level, which must be at least compensated by selective advantage expressed as a large-deviation probability [31], to explain the persistence of evolved specificity. A denomination of all rate effects, from physiological to organism-population and ecological scales, in common units of large-deviation probability, is meant to provide robust lower bounds on constraints of thermodynamics on fitness, without regard to particular molecular and other mechanisms that may connect the two (and that may create more stringent bounds) in nature.

## Acknowledgment

ES and HBS acknowledge support from JSPS grant # 22K03792, and support through the Earth-Life Science Institute from the Japanese ministry of Education, Culture, Sports, Science and Technology. The authors wish to thank Christoph Flamm for extensive input during the course of this work.

## Appendix A The stochastic-process generator in graphical form

This appendix groups the reactions in the sugar-phosphate example that are images of the same mechanism in the MØD network expansion, and displays each group graphically on the network representation from Fig. 2. The five rules from [2] are listed by name and description in Table I, and described as transformations on aldose and ketose sugar-phosphates by carbon number in Table II. (A sixth rule from [2], the phosphorylating cleavage reaction performed by phosphoketolase, is not used as it is not a feature of the sugar-phosphate re-arrangement system responsible for the Calvin-Benson cycle and the standard Pentose Phosphate pathway.^30^)

The platform MØD was used to recursively generate a stoichiometric network from seeds of glyceraldehyde-3-phosphate and water, up to size C_8_ in bisphosphates and their associated ketose-monophosphates. The resulting network contains 17 species and 28 bidirectional reactions. Fig. 18 shows the reactions in the full graph grouped by the rules (reaction mechanisms) that MØD embeds by double-pushout in literal molecules to generate them.

## Appendix B Characterizing the null space of the sugar-phosphate conversion network

In the sugar-phosphate example presented here, the set of solutions *v* to the problem *J* = 𝕊*v* with fixed *J* may be written as any one solution plus the null space ker 𝕊. A basis for that kernel can be computed from general algorithms such as row reduction. However, as noted in the text, the elaborate system of constraints from the generating rules of Table I and Table II imposes many regularities on all such solutions, including a prelude/fugue decomposition and several overall symmetries and conserved quantities.

We can obtain a fuller understanding of the structure of the null space, and thereby of solution spaces, by choosing a particular basis organized according to symmetry, which this appendix presents. In addition to understanding, we can show with a counting argument that the classification of basis elements is sufficient for the null spaces of graphs that extend the one given here to any largest molecule size.

### 1. A rule-derived basis for the null space of balanced flows

We begin with the prelude/fugue decomposition from Sec. III A 4, which highlights a rule composition of aldol condensations and phosphohydrolyses that will fit into the later symmetry classification of null flows. Fig. 19 exhibits what we have termed the four “pure” preludes in the network from Fig. 2. These are the preludes that convert two molecules of the same aldose phosphate, plus DHAP, to two molecules of the same bisphosphate. Six other “mixed” preludes (four-choose-two) may be formed by combining one-half of the flows in any two of these panels.

Fig. 20 shows that three independent null cycles are formed by joining two adjacent preludes activated in opposite reactions with a single TKL re-arrangement edge. For convenience we treat each edge as activated exactly once, equivalent to taking one-half the flow in the “pure” preludes on the same graphs.

Fig. 21 shows three independent TAL trefoils in ker S for the network of Fig. 2. We note in passing that, had the network expansion by MØD been terminated with *B*_7_ as the largest bisphosphate, the TAL trefoil with the lowest carbon count would be the only one remaining in the network. We give a counting argument later for the binomial coefficient that counts independent TAL trefoils as a function of the largest bisphosphate in the network.

Fig. 22 shows six independent TKL trefoils in ker S for the network of Fig. 2. Had the network only been extended to *B*_7_ as its largest bisphosphate, only the three with the lowest carbon count would have been present.

Finally Fig. 23 shows the two adjacent null flows in ker S shifting the action of AlKe upward by one carbon. These are the two instances with the general graph topology from Fig. 8 in this network.

### 2. Saturating the counting of the null space, and induction on the largest molecule size

We show next how null cycles of each of the above categories can be counted, as a function of the limiting molecule size in the network expansion. We begin in Sec. B 2 a with an analysis of the trefoils as the minimal non-trivial braided cycles, and explain the maps by which each counting problem for them is reduced to a binomial coefficient. The continuation of the same counting is used in Sec. B 2 b to show that the four basis categories constructed from symmetry in fact span the null space of the reaction graph at any size.

#### a. The counting of trefoil cycles

We first observe from Table IV and Fig. 9 that the two self-compositions TAL^2^ = TKL^2^ = *I*, the identity on aldose-ketose complexes. It follows that no null cycle of period 2 can be made from only TAL or TKL edges. Fig. 24 diagrams the smallest kind of non-degenerate braided loop as a hypergraph, for the example of a TAL trefoil on three backbones of length (*m, n, p*).

To streamline the following arguments, we invert the index notation to make the molecule size rather than type the independent variable, and the type a labeling index.

Where, in the main text, we have labeled chemical species *A*_*n*_, *K*_*n*_, *B*_*n*_ for aldose- or ketose-monophosphate and bis-phosphate sugars, here we let *n*_*A*_, *n*_*K*_, *n*_*B*_ stand for the respective carbon counts in aldose, ketose, or bisphosphate sugars, according to where *A, K*, and *B* species enter a calculation.

We proceed in four steps to count first the number of null cycles by the foregoing classification, inductively on the network size, then to compute the species and edge counts and dimension of the null space, and finally to show that the null flows accounted for in the above classification saturate the dimension of the null space.

- Let *n*_*B*,max_ ≡ max *n*_B_ in an exhaustive enumeration to a finite bisphosphate size be the variable of induction.

#### **Range of** *n*_*A*_ **and** *n*_*K*_ **through which braided cycles can pass:** The smallest aldose is *A*_2_. For any TAL trefoil, the smallest ketose compatible with the schema (38) will be *K*_5_. In comparison, the smallest ketose compatible with the TKL schema (40) will be *K*_4_. (In the example network of Fig. 2 there are thus five compounds a TKL trefoil can visit and only 4 for the TAL trefoil.)

#### Counts of TAL and TKL edges

Any complex 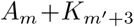 with *m ≠ m*′ in the admissible range is a valid input complex defining a non-null TAL edge, or *A*_*m*_ + *K*_*m*′+2_ for a TKL edge, in the schemata of Table II. Call the number of eligible *m* values (*D* + 1)_TAL_ or (*D* + 1)_TKL_, respectively; they will be the vertices of simplices of dimension *D*_TAL_ or *D*_TKL_. TAL or TKL reaction edges are the complexes *A*_*m*_ +*K*_*m*_′_+3_ or *A*_*m*_ +*K*_*m*_′_+2_, formed as pairs of non-identical (*m, m*′) in the respective simplices, so the number of distinct edges will be (*D* + 1)-choose-2 in the appropriate simplex.

#### Decomposition of general braided cycles into trefoils

Braided cycles of either TAL or TKL edges map to simple paths on some set of aldose indices {*m, n, p*, …}. Because these are now ordinary paths on edges of a simplex, they can always be decomposed into sums of elementary 3-cycles which are the trefoils.

#### Rank of TAL and TKL null spaces

Fig. 24 shows how the TAL or TKL hyper-edges in trefoils stand in 1-to-1 correspondence to ordinary 3-cycles on the vertices of the appropriate *D*-simplex. The rank of any basis for all circuits on the simplex will be the number of independent dimensions of flow along its edges. Although these can be implemented with a subset of 3-cycles on faces, the number of independent cycles will be the number of distinct rotations in *D* dimensions. Any rotation specifies a plane, which is defined by two dimensions. So the rank of the null space, or the number of independent trefoils that can be constructed, is *D*-choose-2.^31^

We relate the values *D*_TAL_ and *D*_TKL_ to whole-network size in the next subsection.

##### b. Topological and cycle basis measures of the null dimension

##### a. Counting number of edges

In a graph with the largest bisphosphate carbon count *n*_*B*,max_:

- The number of AL edges will be 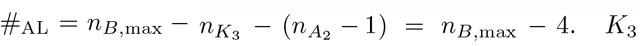 is DHAP, input to all AL reactions. *A*_2_ (glycolaldehyde-2-phosphate, or GLP) is the smallest aldose-monophosphate.
- The number of AlKe/KeAl edges will be 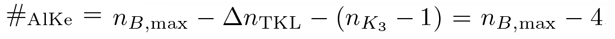. Here Δ*n*_TKL_ = 2 and (*n*_*B*,max_ − Δ*n*_TKL_) is the size of the largest aldose monophosphate produced in a TKL reaction by C_2_ exchange from the largest ketose-monophosphate. *K*_3_ is the smallest ketose capable of isomerization.
- The number of aldose vertices coupled through TAL edges, which we have denoted (*D* + 1)_TAL_, is given by 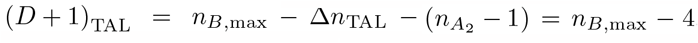. Here Δ*n*_TAL_ = 3 is the size of the C_3_ end exchanged between the largest ketose in the graph and the largest aldose that can appear in a TAL edge. The number of TAL edges connecting all pairs of vertices is #_TAL_ = (*n*_*B*,max_ − 4) (*n*_*B*,max_ − 5) */*2, which is (*D* + 1)_TAL_-choose-2.
- The same construction for the number of ketose vertices coupled through TKL edges, denoted (*D* + 1)_TKL_, is given by 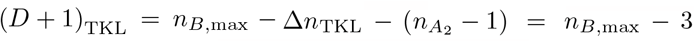. The number of TKL edges connecting all pairs of vertices is #_TKL_ = (*n*_*B*,max_ − 3) (*n*_*B*,max_ − 4) */*2, which is (*D* + 1)_TKL_-choose-2.
- Adding the above gives the total edge number: #_edges_ = #_AL_ + #_PHL_ + #_AlKe_ + #_TAL_ + #_TKL_ = (*n*_*B*,max_ − 4) (*n*_*B*,max_ − 1).

##### b. Number of species, stoichiometric dimension, and nullity by edge counting

- H_2_O and H_3_PO_4_ contribute 2 species in an enumeration to any maximum size *n*_*B*,max_.
- The number of bisphosphate species in the graph relates to their maximal size in the same way as the count of the AL reactions that produce them: #_B−spec_ = #_AL_ = *n*_*B*,max_ − 4.
- Aldose monophosphates start at *A*_2_, which is one unit smaller than ketose monophosphates, so #_A−spec_ = #_AlKe_ + 1 = *n*_*B*,max_ − 3.
- Ketose monophosphates are found at every count larger than 2, so 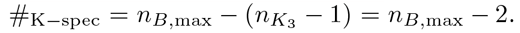.
- The number of species passed through in the enumeration is then the sum #_spec_ = 2 + #_B−spec_ + #_A−spec_ + #_K−spec_ = 3*n*_*B*,max_ − 7.
- There are only three independent conserved quantities with this rule set, which are the counts of C, P, and O atoms (H is fixed by the octet rule). So the dimension of the stoichiometric space is *s* = #_spec_ − 3 = 3*n*_*B*,max_ − 10.
- The nullity is the reaction number minus the stoichiometric dimension, so #_null_ = #_edges_ − *s* = (*n*_*B*,max_ − 4)^2^ − 2.

##### c. Count of independent null basis cycles

- The number of null aldol cycles is one fewer than the number of AL reactions: #_AL−null_ = #_AL_ −1 = (*n*_*B*,max_ − 5).
- The number of AlKe null cycles is *two* fewer than the number of AlKe reactions, because the coefficient 2 of the TIM reaction *A*_3_ *⇀ K*_3_, and a constraint of one additional AlKe conversion, is fixed by the stoichiometry of the net conversion (34) (which may be written 5 *A*_3_ ⇌ 2 *A*_0_ + 3 *K*_5_) and the conservation laws of the rule system. The count of null cycles from the remaining, variable AlKe reactions is then #_AlKe−null_ = #_AlKe_ − 2 = (*n*_*B*,max_ − 6).
- The number of independent TAL trefoils is #_TAL−null_ = (*n*_*B*,max_ − 5) (*n*_*B*,max_ − 6) */*2, which is *D*_TAL_-choose-2 for *D*_TAL_ the dimension of the simplex bounded by TAL edges, computed above.
- The number of basis TKL trefoils is #_TKL−null_ = (*n*_*B*,max_ − 4) (*n*_*B*,max_ − 5) */*2, or *D*_TKL_-choose-2.
- The sum of the four groups of independent basis cycles is #_null_ = #_AL−null_ +#_AlKe−null_ +#_TAL−null_ + #_TKL−null_ = (*n*_*B*,max_ − 5) (*n*_*B*,max_ − 3) − 1.given for the nullity above.

By comparing the final values #_null_ = (*n*_*B*,max_ − 4)^2^ − 2 from part B 2 b b, and #_null_ = (*n*_*B*,max_ − 5) (*n*_*B*,max_ − 3) − 1 from part B 2 b c, we confirm that the independent null cycles constructed by symmetry account for the full rank of the null space computed from the link number and stoichiometric dimension, at any *n*_*B*,max_ These cycles are therefore a basis. ■

## Appendix C Integer flow ILP enumeration and properties

This appendix contains several count statistics from the list of 763 integer-flow solutions fi to the problem *J* = 𝕊fi, with *J* given by the net conversion (34) and 𝕊 the stoichiometric matrix for the full reaction graph of Fig. 2. The flows are produced as solutions to an integer-linear program (IPL) within the software platform MØD. They are generated in an ascending order of the number of distinct reactions in the supporting graph, and then of the sum of absolute counts on all reactions (unsigned), meaning that all solutions at a given value of these properties are exhaustively enumerated before the first solution is given at a higher value of either count.

An initial list of 800 solutions (the upper limit chosen arbitrarily) fi; *i* ∈ 1, …, 800 was generated. From these, 37 solutions were found to be stoichiometrically equivalent to earlier solutions in the list, because the ILP solver had produced partly-canceling currents on oppositely-directed unidirectional reactions. These duplicate solutions are eliminated from all results here, but the label *i* in fi was left in place from the raw list. Any solution fi used in the text is given explicitly in one of the tables.

### 1. Classification of irreducible and reducible integer flows for the sugar-phosphate example

#### a. Summary of solution types

- The list contains **763 stoichiometrically distinct flows**.
- All reducible flows have supporting graphs supp fi with exactly one null trefoil cycle, which may be either a TAL or a TKL trefoil.
- 19 subgraphs arise twice as supp fi for two distinct flows in the list. (By construction the two solutions sharing a supporting graph differ by an integer multiple of a trefoil flow.) Hence there are **744 distinct supporting graphs** from this list.
- Among the supporting graphs with null cycles, 7 distinct trefoils are found in the list. 2 are TAL trefoils and 5 are TKL trefoils.
- The list contains **85 stoichiometrically distinct reducible flows**. Of these 15 have supporting graphs with TAL trefoils and 70 have supporting graphs with TKL trefoils.
- The 85 distinct reducible flows include the 19 pairs mentioned above, with a supporting graph shared by the two flows in each pair. The list therefore contains **66 distinct supporting graphs for reducible flows**.
- For 12 out of the 85 reducible flows, the minimum-dissipation solution (solution to the mass-action rate law at *j*^‡^ = 1) on the same supporting graph differs from the ILP solution fi by an integer multiple of the trefoil.^32^ These are indicated with dots without (+) markers in Fig. 17. All 12 are included in the 19 pairs that duplicate supporting graphs.
- The remaining 73 reducible flows have minimum-dissipation solutions that differ from fi by some trefoil current *x/*3 for *x* ∈ {1, 2, 4, 5}. These are indicated with dots and (+) markers in Fig. 17.
- From the 744 distinct supporting graphs, there are 744 − 66 = 678 **distinct supporting graphs for irreducible flows**. All supporting graphs for reducible flows in this set can be written as unions of supporting graphs within the set of 678 for irreducible flows.
- Only the six supporting graphs for the 12 reducible flows on which the minimum-dissipation flow coincides with an earlier irreducible flow, because the ILP solution contains a “superfluous edge”, fail to determine distinct minimum-dissipation flows. Hence there are 744 − 6 = 678 + 60 = 738 **distinct minimum-dissipation flows** determined by supporting graphs from this list.

**FIG. 17:**
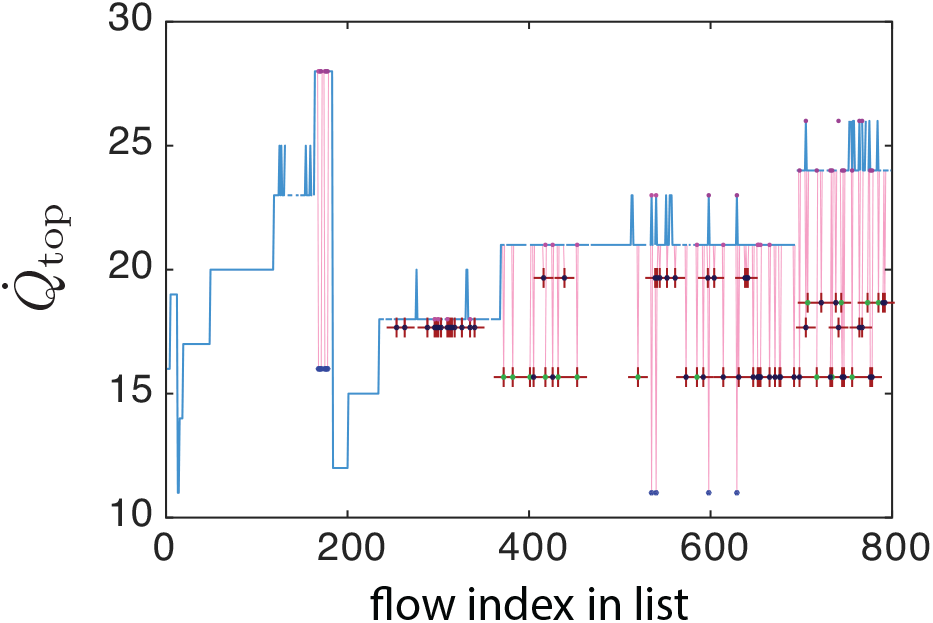
The topological dissipation 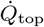 from Fig. 4 corresponding to Eq. (116) for integer solutions *fi* (blue), compared to the minimum dissipation 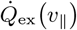 of Eq. (115) (pink) on the same supporting graph at uniform background *j*^*‡*^ ≡ 1. For irreducible flows the two are the same, and for reducible flows 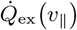 falls below 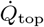. Each reducible flow contains a single null cycle, which is either a TAL (green dot) trefoil or TKL (blue dot) trefoil. Dark-red (+) mark the 73 flows in which the 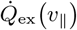 -minimizing trefoil current differs from the solution fi by 1/3 integer value. Dots without (+) markers are 12 flows at 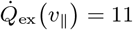 or 16, with trefoil currents that differ from fi by integer values. See App. C 1 for details. For all those the 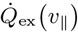 -minimizing solution matches an integer solution on a graph earlier in the list with one fewer reaction.

**FIG. 18:**
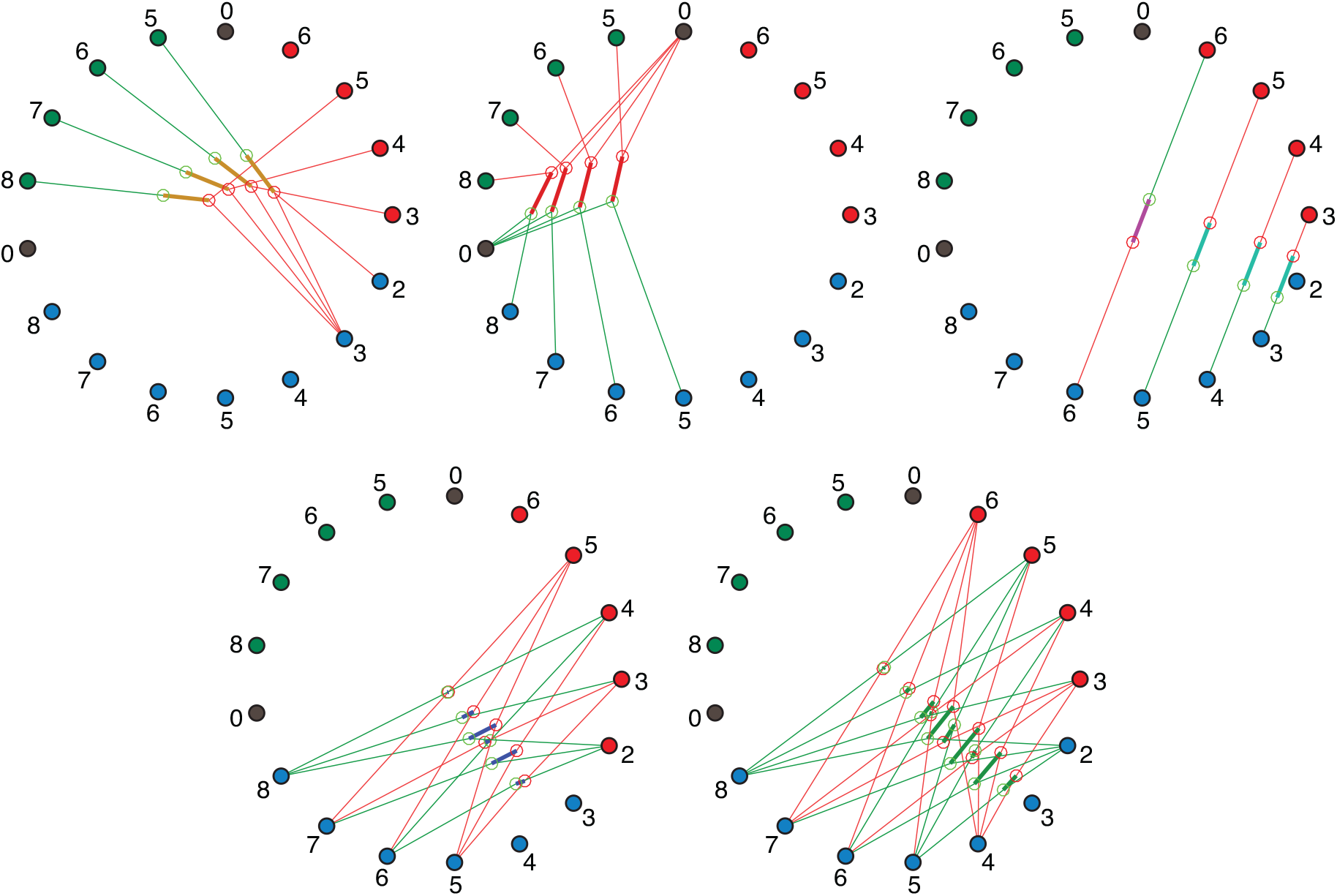
The edges from Fig. 2 grouped by the generating rule. Panel a) aldol/retro-aldol; b) phosphohydrolase; c) aldose-ketose conversions; d) transaldolase; e) transketolase

**FIG. 19:**
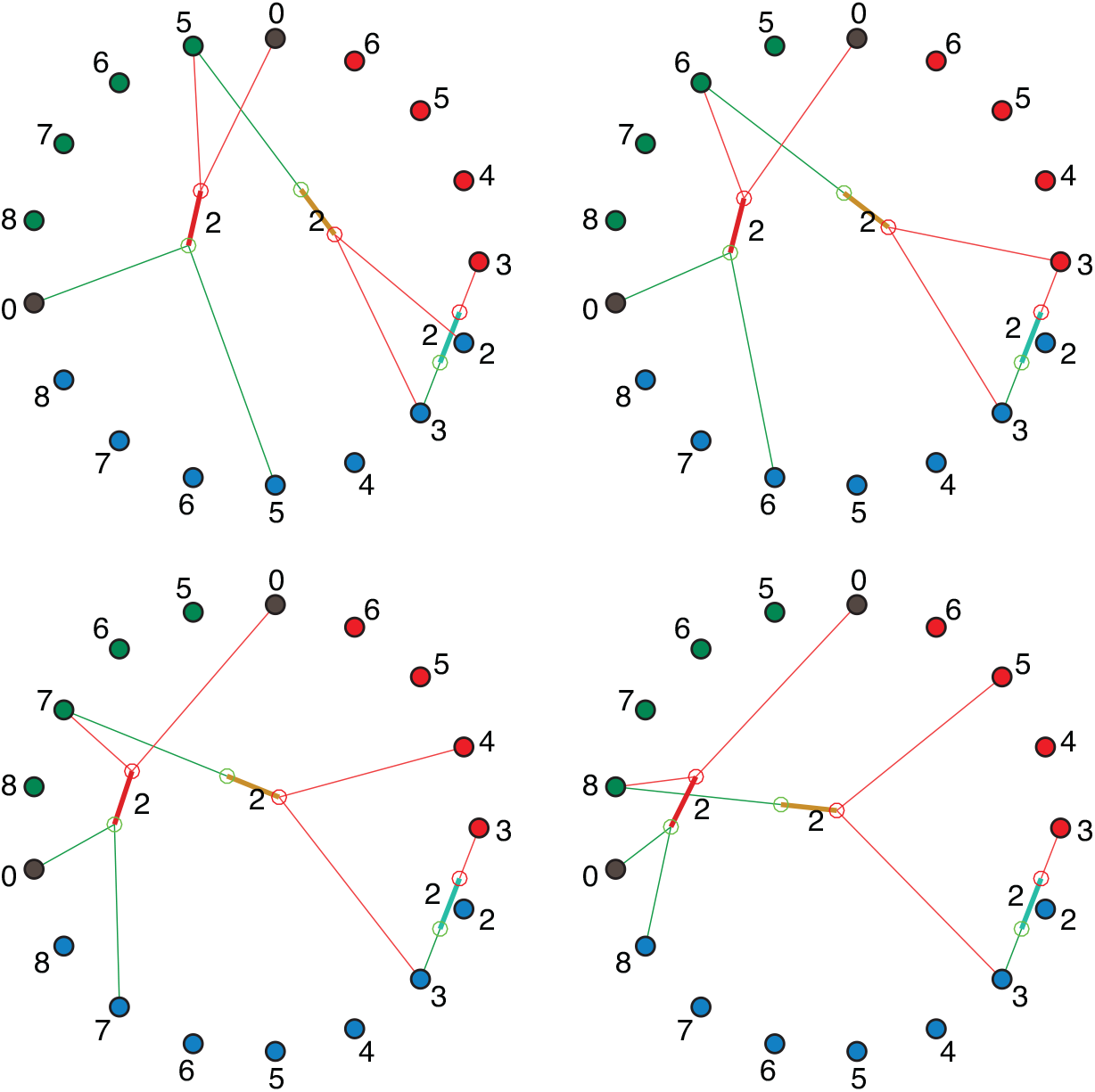
The four pure preludes with *B*_5_, *B*_6_, *B*_7_, and *B*_8_ bisphosphate intermediates.

**FIG. 20:**
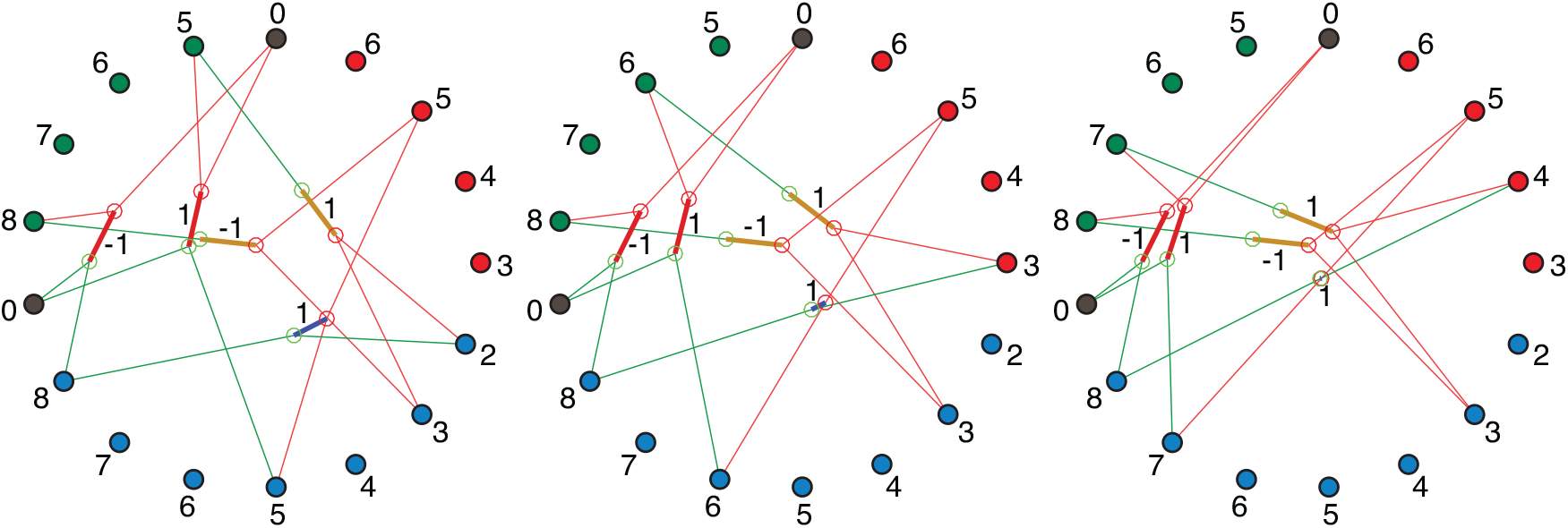
Three independent prelude differences, each inverted by a single TAL edge to form a null cycle.

**FIG. 21:**
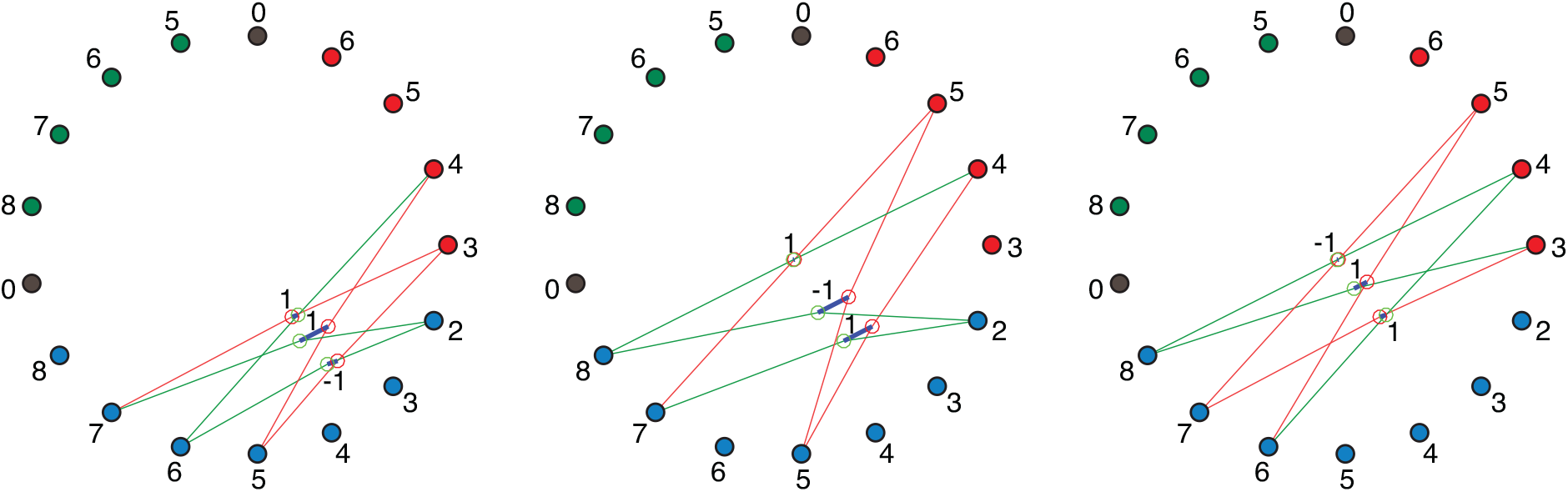
Three linearly independent TAL trefoils that form a basis for all null TAL cycles in the set.

**FIG. 22:**
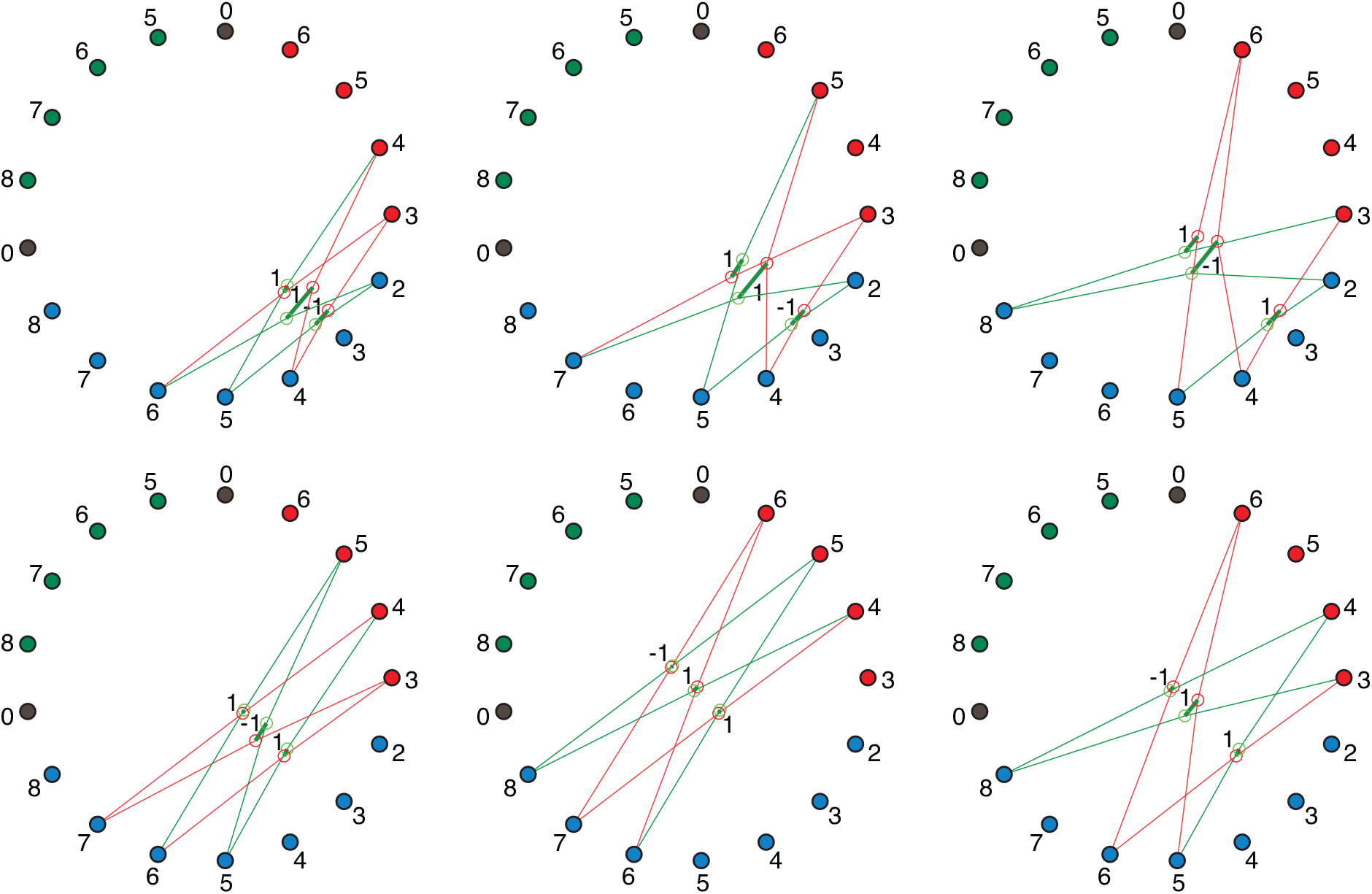
Six linearly independent TKL trefoils that form a basis for all null TAL cycles in the set.

**FIG. 23:**
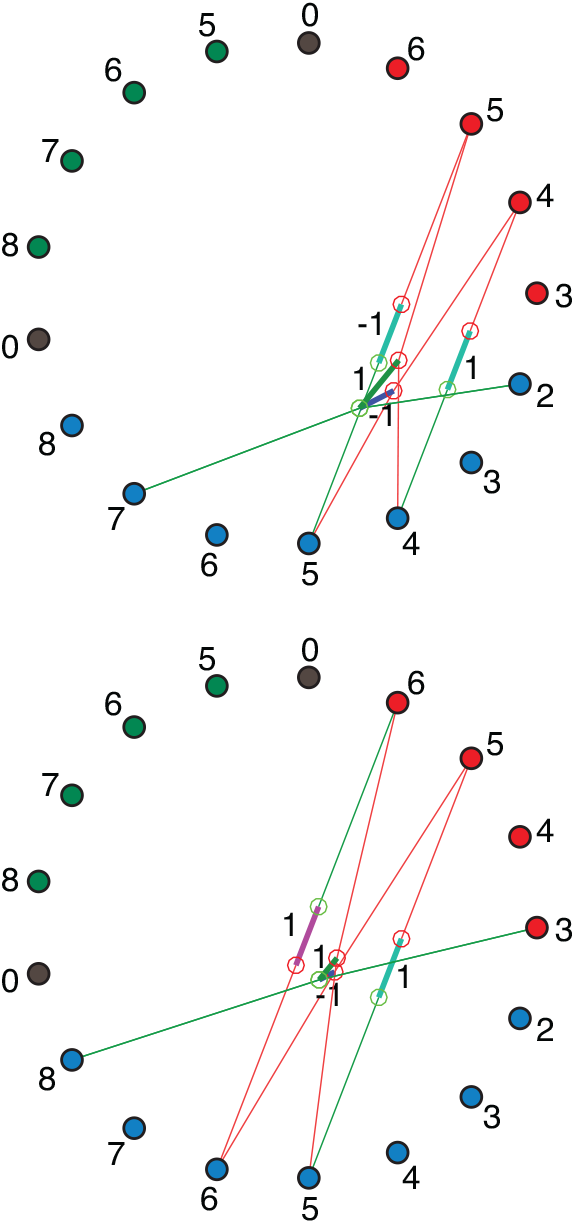
A basis for null cycles from opposed AlKe ⊕ KeAl acting at adjacent *A*_*n*_ and *K*_*n*+1_ cyclized with a composition TAL ° TKL from Table II.

**FIG. 24:**
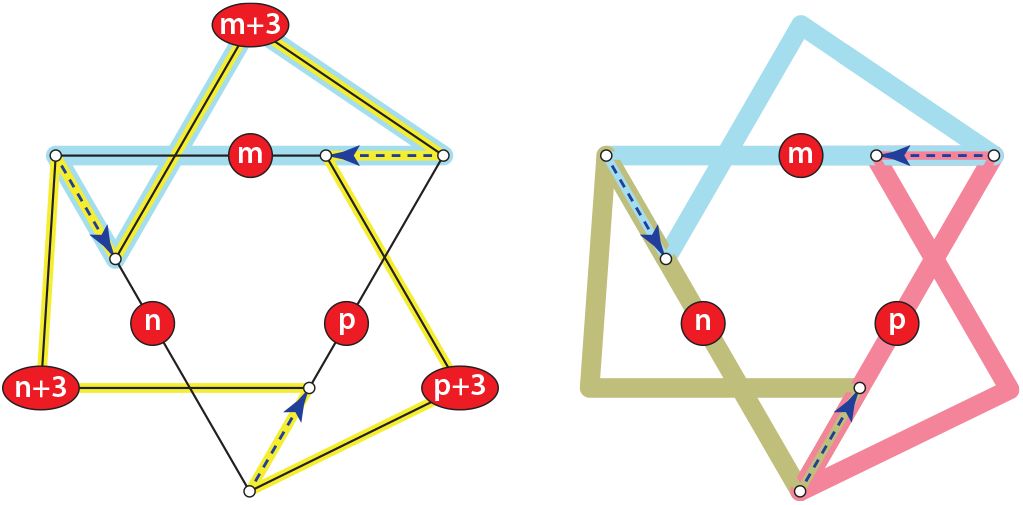
Hypergraph diagram of a TAL trefoil corresponding to the schema (38) Lefthand panel: Yellow circuit is the period-3 path of the C_3_ sugar end carried, bucket-brigade style, by ketoses. Cyan circuit is the period-2 path of the *m*-carbon backbone that is recycled by the two TAL edges in which it appears. Righthand panel: each period-2 circuit (colored bowties), identified with a backbone number *m, n*, or *p*, is a vertex on the simplex made by the aldoses connected through all TAL edges in Fig. 18d). Each reaction links two such vertices, so reactions are edges on that simplex. Note that each colored loop is the boundary of an oriented surface. For an odd number of oriented surfaces to have a boundary that cancels everywhere on their overlap, the surfaces must form a Möbius band (more directly visible in Fig. 14).

These statistics are summarized in Table. X.

## Appendix D Lattice-graph representation of rule symmetries

This appendix describes in detail what we have called the “algorithmic” structure of the lattice diagrams for integer-flow solutions shown in Fig. 10. It includes proofs of the minimality of 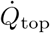 in the solutions f14 and f13 and the relation of minimality to self-seeding, and explains why reduction in the number of reactions in the supporting graph from 8 to 7 must increase the value of 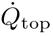 from the minimizing value.

### 1. Flow realizations and their representation by lattice diagrams

An integer flow solution *v* to *J* = 𝕊*v* for the conversion schema (34) determines only the concurrent fluxes on a subset of reactions in a graph 𝒢. For any flow *v*, multiple orderings of integer activations of the reactions can generally be chosen, with each reaction in the sequence acting on a state of the population produced by the immediately preceding reaction. In the Petri-net literature these are called *realizations* of the flow [34]. The realization chosen for a flow, together with the initial population state, dictates a specific sequence of states traversed in the state space.

A lattice diagram for a realization of a flow *v* as we have constructed them in Sec. III D, although not a map of literal translations in a state space, is isomorphic to a state-transition map up to translation in the lattice of states. It coincides with the smallest-population embedding in the state space with respect to all species, at which every reaction is *realizable* (the state departed contains the input complex to the reaction).

The lattice graph for a flow realization therefore is a concatenation of the reactions in Table IV and Table VI summing to the integer counts in *v*, which also comprises one or more simple loops, each of which returns to its starting lattice point when all steps are taken. The set of realization of a flow *v* corresponds to the set of possible concatenations satisfying these conditions.

### 2. Self-seeding and autocatalytic flow realizations

If a flow can be realized with a cycle passing through the origin in the lattice of complexes, then it can be realized from only the starting molecules 5 GAP + 2 H_2_O of the schema (34). Such a flow is *self-seeding*, in the terminology of [130, 131]. Otherwise the flow is *autocatalytic*, and one or more 1-complexes (*A*_*n*_) or (*K*_*n*_) are necessary “seed compounds” to realize the flow, as suggested in the general prelude-fugue partition of Eq. (35) and Eq. (36).

### 3. Algorithmicity and lower bounds for reactions and activities of minimal flows

The series of flows mapped in Fig. 10 of the main text, obtained as f14, f13, and f194 in the enumerated ILP list, are the first three members of an infinite series of such solutions with the common architecture that they perform the same conversion on two 1-complexes *A*_*n*_ and *A*_*n*+1_, followed by a return to the starting compound. These solutions use no TAL edges and only one AlKe edge each (beyond the flux 2 in the TIM reaction, common to all solutions), escaping redundancy and in that way providing minimal solutions to the problem of shuffling the relatively prime *A*_3_ inputs and *K*_5_ outputs.

The limited diversity of rules, inputs, and outputs isolates a shortest reaction sequence (shown as the fourth panel in Fig. 10) that is applied repeatedly in all these solutions. That sequence contributes a fixed number of reactions and a fixed summand in 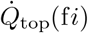 in all cases except the lowest two (f14 and f13, the first two panels in Fig. 10), which are shortened by passage through a fixed-point complex of the TKL symmetry.

#### a. An architecture of iterated loops and a reset

The figure shows that the absence of a common divisor of the *A*_3_ input and the *K*_5_ output necessitates the fusion *A*_*n*_ + *K*_3_ *⇀ K*_*n*+3_ + *A*_0_, leaving either an excess by C_1_ or a deficit by C_2_ after *K*_5_ is extracted (using a TKL edge to shift the C_2_). A composition of the subset of reactions (Ru5P_out ° TKL ° GAP_in ° PHL ° AL ° DHAP_in), using no AlKe conversions and supporting no trefoils, is an algorithmic “loop” that increments the counter *n* by 1 in its starting complex (*A*_*n*_). After two iterations of this loop, the C_2_ deficit in the complementary composition of reactions (Ru5P_out ° TKL ° GAP_in ° AlKe), which includes the minimal required additional AlKe instance, can then be used to reset the counter to its initial value.

#### b. A lower bound on the topological contribution 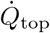 to dissipation

The first two flows shown in Fig. 10 use 8 distinct reactions, and are the unique solutions with the lowest value of 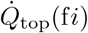 in Fig. 4. (The flow in the second panel is the carbon-rearrangement in the canonical Calvin cycle.) Although these are not flows using the fewest distinct reactions, the figure gives a graphic proof of why they are the two *unique* solutions with the smallest 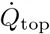 from Eq. (33). Both flows pass through the complex (*A*_3_, *K*_5_), which is a fixed point of the TKL symmetry and does not require an active TKL edge. It is the *only* such fixed-point complex in Fig. 9 that mediates input and output directly, and so supports the flow without requiring other edges to convert to GAP and Ru5P.

**Corollary:** The complex (*A*_3_, *K*_5_) can be exchanged with the complex (0, 0) by reversing the order of the links GAP_in and Ru5P_out, so the first two realizations in Fig. 10 are also self-seeding.

#### c. Reducing edges raises 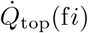

The only way to employ fewer edges – and 7 is the fewest edges attainable with these rules – with the requirement of 3 net AlKe conversions, is to pass all three conversions through the TIM reaction *A*_3_ *⇀ K*_3_ (as the canonical catabolic Pentose-Phosphate Pathway does). Doing so, however, yields two *A*_4_ species relatively prime with both *A*_3_ and *K*_5_, requiring further activations to re-arrange these carbons and resulting in larger 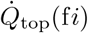 (16–19 vs. 11) in all solutions with fewer than 8 reactions in Fig. 4.

## Appendix E Thermodynamic landscapes

To assign values to relative dissipations or pathway resistances in the regime of linear response near equilibrium, we must identify a thermochemical landscape from which to derive values of the bidirectional equilibrium transition-state currents *j*^‡^. As noted in the main text, we may adopt an assumption of “barrierless” catalysis, in which the transition-state free energy equals the activity of the lower-activity complex of the reaction at the operating equilibrium, and the half-reaction rate constant from the higher-activity complex equals the equilibrium constant (up to a scale factor for rates common to all reactions). The relevance of topological indices such as _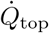_, computed about a uniform background *j*^‡^ ≡ 1, to properties of actual networks will be determined by the degree to which complex activities at physiological conditions approximate uniform complex activities and that rate constants are evolved to be near the diffusion limit of substrates to the active sites of catalysts.

### a. An automated pipeline from graph grammars to thermochemical landscapes

Our emphasis on rule-based modeling is meant to take advantage of high-powered computational platforms such as MØD to construct large graphs and to enumerate exhaustive solution lists in a high-throughput fashion. Therefore we have sought to obtain thermochemical data in a similar high-throughput manner that can be assembled in an automated pipeline from the input rule declaration to prediction of macro-parameters such as pathway resistances (117).

Our source for thermochemical data is the platform eQuilibrator 3.0 [62, 87], which uses sophisticated modern group-contribution algorithms to assign free energies of formation from molecule descriptors such as SMILES or INCHI representations, which are calibrated against database values from KEGG [132–134]. eQuilibrator 3.0 can take as input either stereochemical or non-stereochemical SMILES descriptors. The current version of MØD outputs only non-stereochemical SMILES. However, for the small network in this paper, almost all compounds are known and well-characterized metabolites, for which we can obtain the full stereochemical estimates by name searches within eQuilibrator. Therefore we have computed two thermochemical landscapes, which we term eQNS (for non-stereochemical SMILES), and eQS (for stereochemical values that closely approximate KEGG ground truth). This appendix lists the formation and reaction Δ𝒢^′0^ values and other thermochemical data used to produce results in the main text.

### 1. Standard-state free energies of formation and reaction

Table XI gives standard-state free energies of formation for the species in the model example. Any equilibrium background is chosen relative to these by setting reference concentrations of some collection of basis species.

Fig. 25 shows Δ_*f*_ 𝒢^′0^ values, with the regressions mentioned in the table used to fill the missing eQS values. In the table and the figure, for the eQS data, we have used the *β*-D-fructose values for both F6P and F16B, because this is the epimer used in both the Calvin cycle and the Pentose Phosphate pathway.

**FIG. 25:**
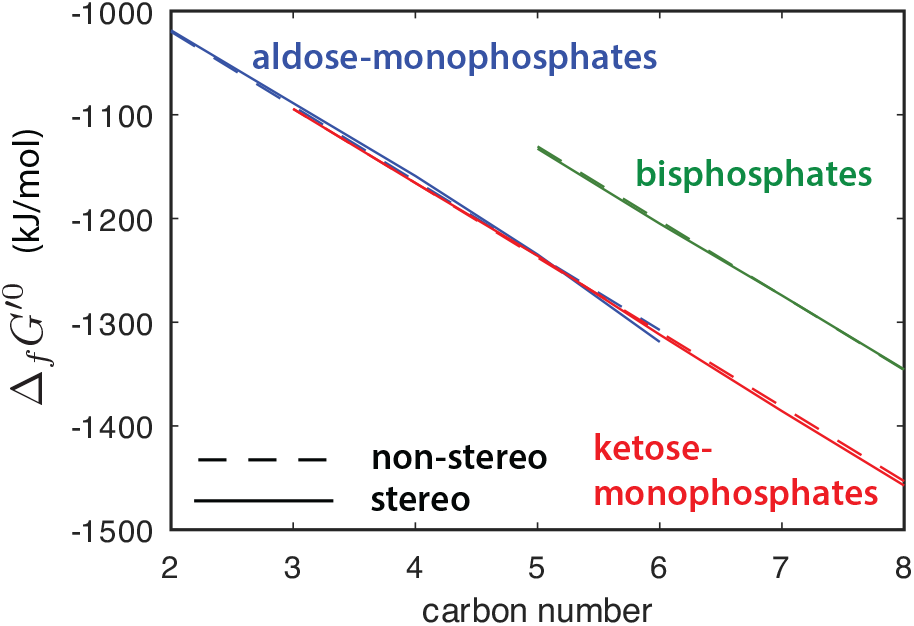
Values of Δ_*f*_ 𝒢^*′*0 (eQS)^ (solid) and Δ_*f*_ 𝒢^*′*0 (eQNS)^ (dashed) from Table XI for the sugar-phosphates. Aldose-monophosphates from eQS are fit with a 2nd-order polyfit to extend to GLP. Ketose-monophosphates are fit with a 1st-order polyfit (linear regression) to assign O8P. Bisphosphates are likewise fit with a linear regression to assign O18BP. The reasoning is that there is apparently stable curvature in the aldoses that is not scatter, whereas bisphosphates have only 3 points (giving no error for a 2nd-order fit), and the ketose-phosphates are not monotonic, with seemingly idiosyncratic variation dominated by Ru5P. The linear fits are tolerable agreement to the eQNS estimates, though we can see that the latter are notably flat for aldoses.

Fig. 26 shows that non-stereochemical group-contribution estimates result in Δ_*r*_𝒢^′0 (eQNS)^s that are nearly constant by reaction type. The eQS landscape diverges from the non-stereochemical estimates in many cases by larger errors than several of the free energy differences between reaction types.

**FIG. 26:**
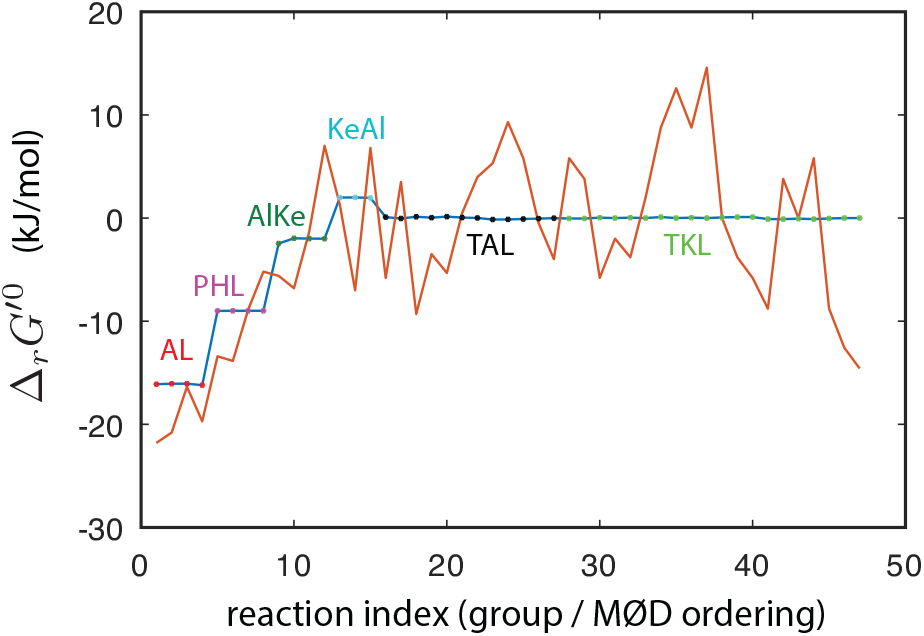
Δ_*r*_ 𝒢^*′*0 (eQNS)^ (blue) and Δ_*r*_ 𝒢^*′*0 (eQS)^ (red) from the stoichiometry and the formation free energies of Table XI. Starting from the reaction order assigned by MØD, the reactions are extracted by groups: AL, PHL, AlKe, KeAl, TAL, TKL, and plotted with the groups given this order. (Within each group, reactions follow the MØD order.) Dots on the eQNS plot mark each group.

#### a. Formation free energies from basis species

To produce a thermodynamic equilibrium reference state *n*, we must convert standard-state free energies of formation and reaction to free energies from basis species, for which we take the first four lines in Table XI: H_2_, CO_2_, H_2_O(*l*), and phosphate, which nominally we put in the form HPO4^2−^ to match the nominal charge state of phosphate groups on sugars.

The synthesis schema for a C_*n*_ sugar with *m* phosphate groups that we label *X* is

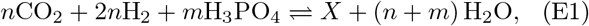

which defines the overall equilibrium constant in terms of the standard-state equilibrium constants of formation as

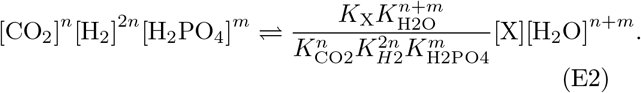

Equivalently, the standard-state free energy to form *X* from the basis species is

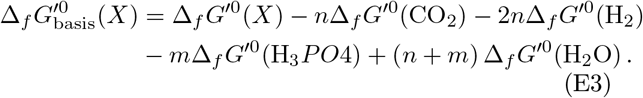

Dividing the free energies of formation from these species by the Carbon count gives an estimate of the heat of formation per CH_2_O group, plotted in Fig. 27.

**FIG. 27:**
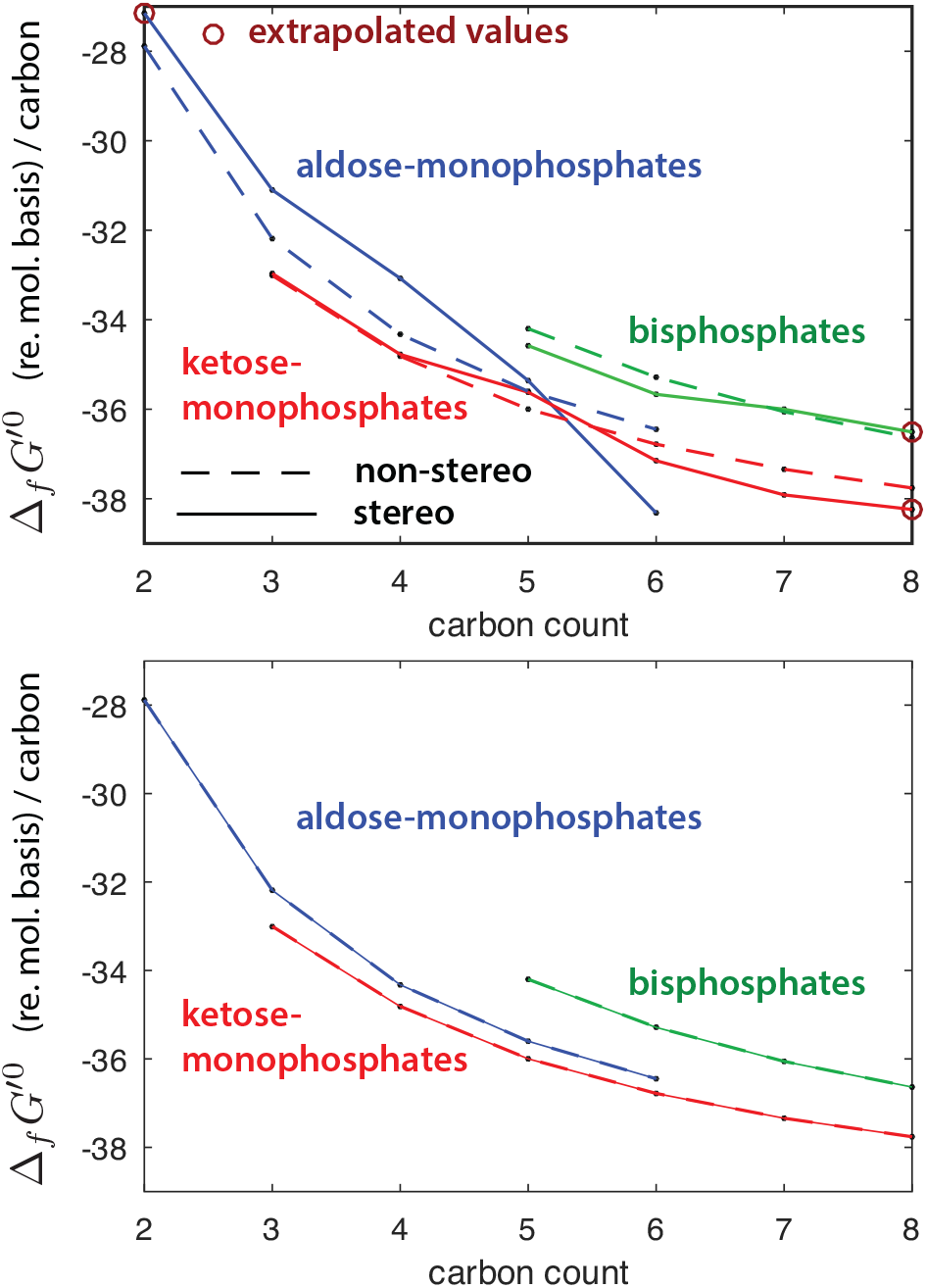
Top panel: Free energies of formation under standard conditions from basis species H_2_, CO_2_, H_2_O(*l*), and HPO4^2*−*^, also under standard conditions. Solid are eQS values and dashed are eQNS group-contribution estimates. The values with circles are the missing eQS values filled in with regressions (red entries in Table XI). Apart from the aldose-monophosphates, all differences are within 1kJ/mol/C. Bottom panel: the eQNS estimate shown alone to emphasize the smooth dependence on carbon number.

#### b. Physiologically pertinent landscapes from standard-state free energies

All reactions in the rule set of Table I conserve carbon and phosphate number. Therefore concentration and free energy profiles are determined by two scaling parameters. For these we choose the concentration ratio [F16BP] */* [F6P], which is controlled by the hydrolyzing potential [H3PO4] */* [H2O], and the “price per carbon”, which is controlled by [GAP] in relation to the Triose-Phosphate Isomerase reaction, which sets [DHAP] in terms of [GAP], and the Fructose-Bisphosphate Aldolase reaction, which sets [F16BP] from [DHAP] [GAP]. The target concentration for [GAP], at a given [H3PO4] */* [H2O], is then controlled by the second equilibrium product [CO2] [H2]^2^*/* [H2O].

Fig. 28 shows two examples of the equilibrium concentrations that result. Both are constructed to put [F16BP] */* [F6P] ≡ 1. The first equilibrium (upper panel) sets a physiological value [GAP] ≈ 50*µ*M for the eQS free-energy landscape, and a tenfold larger value for the eQNS free-energy landscape in order to produce [F16BP] values near the value of [GAP] in the eQNS landscape. The two produce a qualitatively similar decline of concentration with increasing carbon number.

**FIG. 28:**
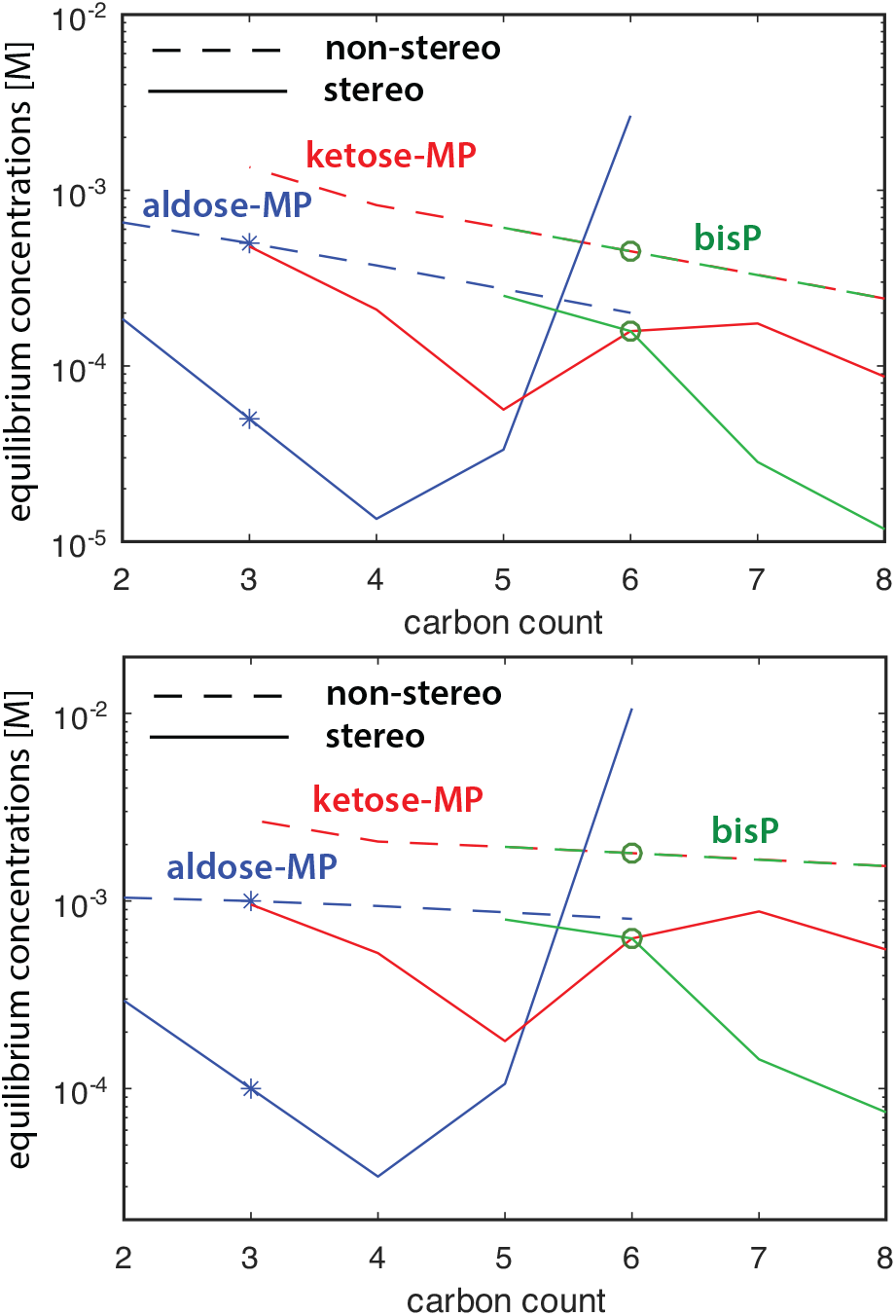
Two concentration landscapes at equilibrium, for both eQS (solid) and eQNS (dashed) Δ_*f*_ 𝒢^*′*0^ values, and hydrolyzing potential set to ensure [F16BP] */* [F6P] ≡ 1 (dark green circles). Upper panel: [GAP] = 50*µ*M (physiological) for eQS free energies, and [GAP] = 500*µ*M for eQNS free energies (blue asterisks), to set [GAP] ≈ [F6P]. These GAP concentrations result in broadly declining concentration with C count in either background. Lower panel: reference concentrations doubled to [GAP] = 100*µ*M for eQS and [GAP] = 1mM for eQNS, producing a nearly flat concentration profile for eQNS, and one without overall trend for eQS, though the latter has large idiosyncratic variation.

A second equilibrium concentration profile (lower panel) doubles the [GAP] values from the first example (to 100*µ*M and 1mM respectively for eQS and eQNS cases), to produce a more neutral profile with respect to carbon addition. The eQNS profile manifests the resulting near-zero carbon cost in nearly-equal concentrations for sugars of all length and the same phosphorylation type. The eQS profile varies by as much as an order of magnitude due to stereochemical formation energies. The furthest outlier from uniformity in the eQS land-scape is G6P, due to its very low free energy of formation (the lowest of the hexoses, and the reason glucose is the main storage and structural carbohydrate in biochemistry). Within the approximation error of the eQuilibrator 3.0 assignments, about 2.5 orders of magnitude separate G6P and E4P concentrations.

#### a. No-barrier equilibrium transition-state activities, and dissipation near linear response

The examples in the text are all computed (for simplicity) in the regime of linear response, where Δ_*r*_*G* values depart linearly from zero in the scale of the through-current. The zero-barrier limit of bidirectional transition-state currents that controls the linear force-flux relation is governed by the smaller activity on the inputs or outputs. Fig. 29 shows these activity products for all reactions in both the eQNS and eQS backgrounds, plotted in the same order as Fig. 26.

**FIG. 29:**
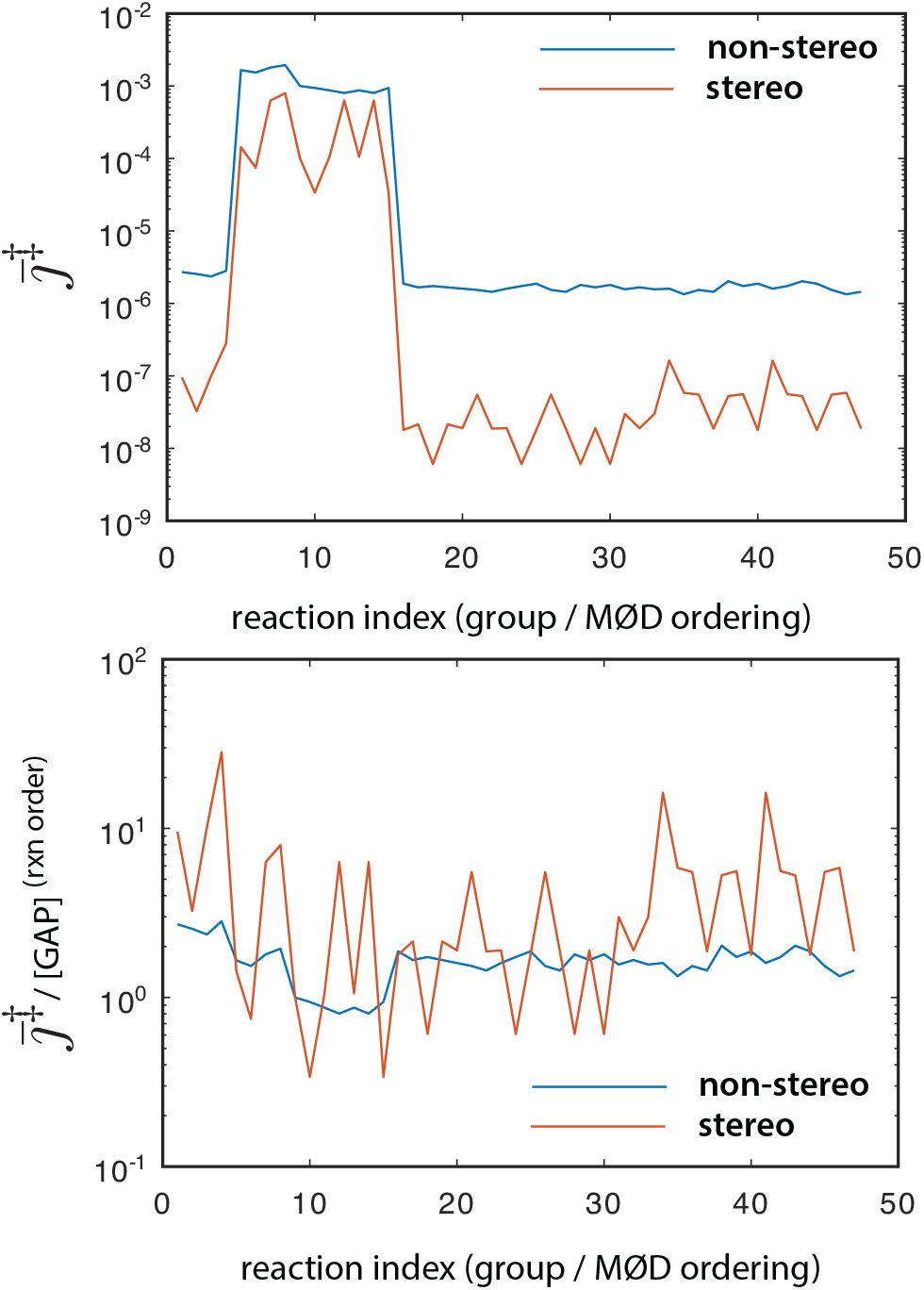
Zero-barrier transition-state currents *<u>j</u>*^*‡*^, defined by the minimum of the activities on the reaction input or output side, for both thermochemical landscapes, in the “uniform” equilibrium from the lower panel of Fig. 28. Upper panel: *<u>j</u>*^*‡*^ in absolute magnitude. Lower panel: *<u>j</u>*^*‡*^ normalized by [GAP] raised to the power of the order of sugar-inputs to the reaction (the reaction degree in organic species). For the eQNS estimator, these normalized activities are in the range ∼1–3. Their broad comparability for the eQS landscape measures the degree to which all sugar concentrations scale with [GAP] in this background.

Comparable concentration for all sugars very crudely approximates physiological conditions (factor of 10 variations from most metabolites to the highest or lowest concentrations) for sugar-phosphate chemistry [136, 137], and is compatible with evolutionary reversal [52] between the Calvin cycle and Pentose Phosphate versions, driven mainly by phosphorylation or dephosphorylation potentials and cofactors and under tight kinetic control [137, 138]. For the very uniform reaction free energies obtained with non-stereochemical SMILES, the corresponding model of equilibrium results in the nearly-constant values for all reactions of the same order shown in the lower panel of Fig. 28.

Fig. 30 shows the minimum dissipation values 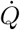(under the mass-action rate law) in the linear-response regime,normalized by [GAP]^2^, on each of the supporting graphs for the 763 valid integer solutions, in the no-barrier current background from Fig. 29. The values of realized dissipation can be compared to the “topological” contribution to dissipation 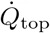 in Fig. 17 in the text.

**FIG. 30:**
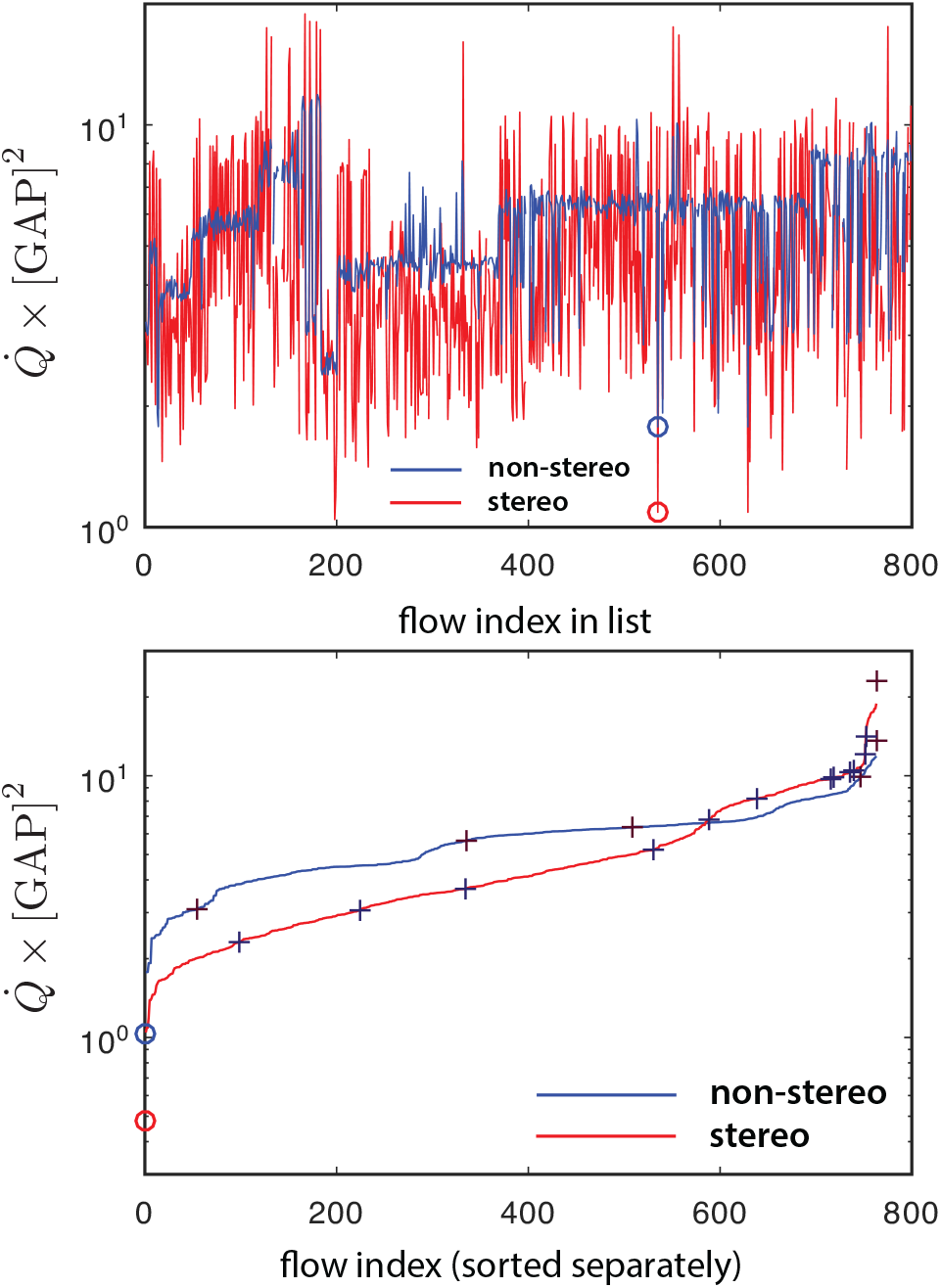
Dissipations normalized as 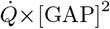 in the linear-response regime on the supporting graphs of the 763 ILP solutions from MØD, for the species-concentration profiles in Fig. 28 and the transition-state activity profiles in the lower panel of Fig. 29. eQNS profile in blue; eQS profile in red. Upper panel: reactions ordered by the original ILP list from Fig. 4. Circles mark the (reducible) flow f535; the supporting graph for which produces the minimal 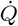 in the eQNS land-scape. Lower panel: reactions sorted independently in the two backgrounds in ascending order of 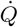. Circles mark the dissipation for the whole network from Fig. 2. The values in the two backgrounds are 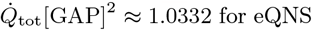 for eQNS, and 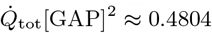 for eQS.

The values for the non-stereochemical SMILES estimates in Fig. 30 vary over a factor of about 5 because the pseudo-first order reactions have reduced impedance as shown in Fig. 29, versus a factor of about 2.5 for 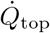. The stereochemically-estimated eQS values show more idiosyncratic variation about the expectation from 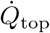 and vary over a factor of about 20.

### 2. Eigenvalues and their relation to low-impedance flows

The metric obtained from the Hamiltonian in Sec. IV E confers on the space of flow solutions a criterion of *orthogonality*, which mere stoichiometric linear independence of integer solutions does not provide. The eigenvectors of the metric then provide an orthogonal basis of flow components equal in dimension to ker 𝕊, to which integer flow solutions can be compared for similarity.

In the lower panel of Fig. 30, the values of 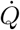 on the integer flow solutions from the ILP list are shown sorted. Interspersed with these are the dissipation rates of flows proportional to the eigenvectors of the metric. If we use an index *q* ∈ 1, …, dim ker 𝕊 to index these eigenvectors, and *α*_*q*_ the coordinates in the decomposition (95) of the main text, then *g*_°_*α*_*q*_ = *λ*_*q*_*α*_*q*_, and we normalize 1^*T*^ *α*_*q*_ = 1, ensuring that the resulting flow satisfies *J* = 𝕊*fα*_*q*_ for *J* the net conversion (34). The dissipation for the full network of Fig. 2 is marked with a circle. It is lower than the value on the supporting graphs of any of the ILP solutions, and lower yet than any of the eigenvectors of *g*_°_.

Among the lowest-impedance irreducible flows in the list are both f13, the canonical Calvin-Benson cycle from the top panel of Fig. 3, and f14, the other topologically minimal solution graphed in Fig. 5. The value 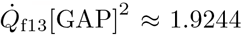, versus 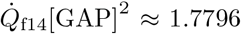 and 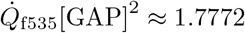, the reducible flow that is the minimal dissipation in the ILP list, and 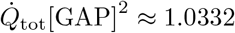 for the whole network.

It happens that for the uniform-concentration background in the lower panel of Fig. 28, f13 is well approximated by linear combinations of the few lowest eigenvectors of *g*_°_ from Sec. IV E, as suggested in Fig. 16. Two levels of approximation are exhibited in Fig. 31. The close approximation of an integer flow by a small number of low-lying eigenvectors of the metric implies that selection isolating that low-lying flow can be performed with minimal increase in dissipative cost to perform the conversion (34). The full topology of the network provides only limited parallelism that can be exploited by the mass-action solution to distribute flow and reduce impedance.

**FIG. 31:**
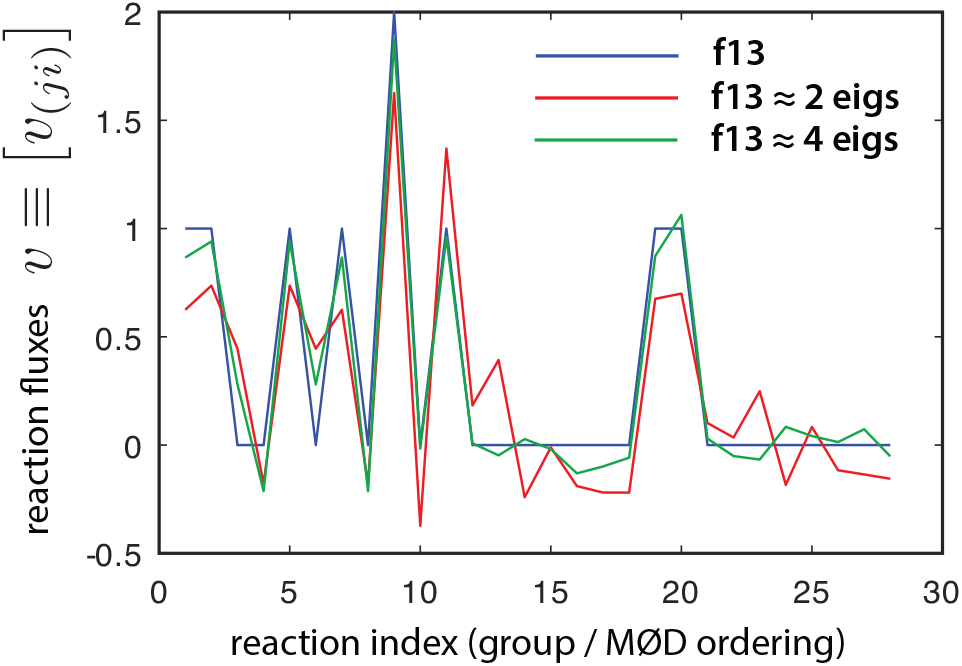
The integer flow solution (blue) for the canonical Calvin cycle (irreducible f13) from the top panel of Fig. 3, expanded in a basis of the eigenvectors of *g*_*°*_. The 28 reactions are listed in the order of Fig. 26. The velocity of each reaction is plotted as the ordinate. Red shows the contribution from eigenvectors 1 and 2; green shows the contribution from eigenvectors 1, 2, 3, and 10, the four largest projections in the L2 norm for eigenvectors (not the distorting trace norm 1^*T*^ *α*_*q*_ = 1).

## Appendix F Inversion of linear-response kernels within stoichiometric compatibility classes

In this appendix we derive the inversion of the kernel matrix from Eq. (18) within the projection of the stoichiometric subspace, which connects thermodynamic forces and fluxes to linear order in response.

We begin by introducing, for any stoichiometric matrix 𝕊, a projector Π_ss_ onto the stoichiometric subspace defined by

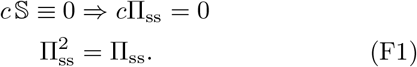

The kernel matrix 𝕊 diag [*j*^‡]^ 𝕊^*T*^ of Eq. (18) has a null space spanned by the conserved quantities *c* of Eq. (F1), and is regular in the stoichiometric subspace. Therefore we may introduce its inverse 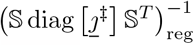 within the stoichiometric subspace, with the property

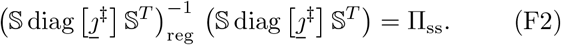

Introduce a notation *δn* ≡ ⟨*n*⟩ − *n* for the offset of *n* resulting from disequilibrium driving in Eq. (17). We must have that *δn* = Π_ss_ *δn*, because for sources that respect the conservation laws of the network (needed for a stationary state), *n* can be deformed only within the stoichiomet(ric sub space. We recall that to linear order *δn/n* ≈ *µ* − *µ*, the shift in chemical potential from equilibrium in the driven state. Then Eq. (17) and Eq. (31) give that

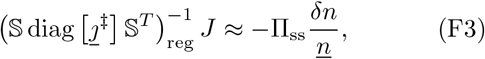

and the full shift *δn* must be given by

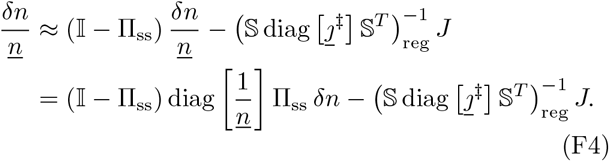

Note that if *n* is uniform then the first term on the right-hand side of Eq. (F4) vanishes; otherwise it gives a shift in the chemical potentials of some conserved quantities that cannot drive any *J*.

Multiplication of Eq. (F4) by diag [*n*] shows that the same terms appear on the left-hand and right-hand sides, so we must have

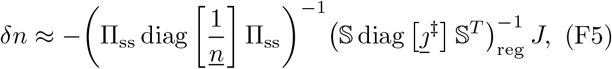

which is solved for *δn* in the stoichiometric subspace ensuring *c* S = 0 ⇒ *c δn* = 0.

The corresponding velocity profile evaluates as

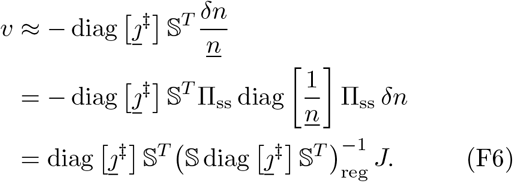

The first term on the right-hand side of Eq. (F4), while nonzero in general, does not contribute to the velocity solution from th e e xternal current once the inversion of the kernel 𝕊 diag [*<u>j</u>*^‡^] 𝕊^*T*^ is performed within the stoichiometric subspace. The quadratic approximation to the excess dissipation given in Eq. (116) then has the two forms

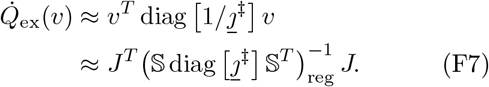

## Appendix G Tilting generators on the events, in closed and then in open graphs

This appendix provides supporting detail for the construction of the large-deviation function for currents in the form of a Lagrange-Hamilton action functional, on closed graphs and on their subgraphs with open boundaries. Some equations from the main text are repeated where doing so makes the presentation within the appendix more continuous and self-contained.

In this appendix we will treat the general case without assumptions of detailed balance in the rate constants, introducing a variety of extra steps in the analysis to separate formal, non-stationary terms in cumulant-generating functionals that do not affect ultimately observable properties, but are present in general as parts of the solution for stationary trajectories. Many of these complications do not arise for networks with detailed balance, and could go un-noticed; we emphasize then that the construction of interest is not limited to those simplifying cases, and can be carried out in the general case.

### 1. Tilting generators on closed graphs

We begin, as in the text, with the LDF conditioned on a single-time deviation from Sec. IV A 4 in the form of the action integral (55):

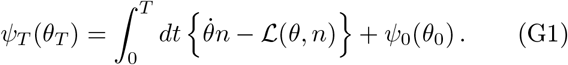

The generator tilted on events with a vector *η* of time-dependent trajectories, from Eq. (60), is written in sev-eral forms from the introductory Sec. II A as

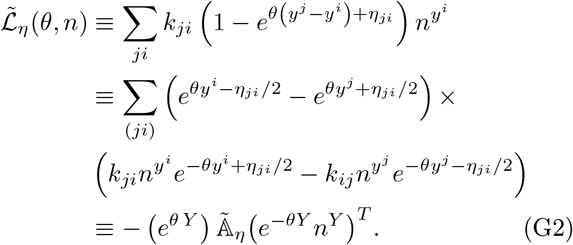

The velocities from Eq. (G2) are again

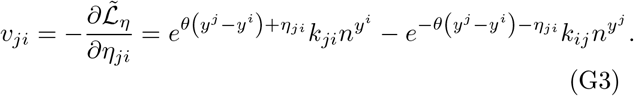

Repeating Eq. (62), we note that the potential part of *η* comes from the terms *θ* that are the Hamiltonian-conjugate momenta to counts *n*, as

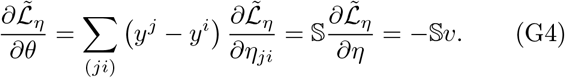

A CGF for the currents in variables (*θ, n*) on a closed graph, with tilting fields *η* in 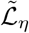, is then constructed from its Hamilton-Jacobi equation just as for the untilted case, giving the integral form

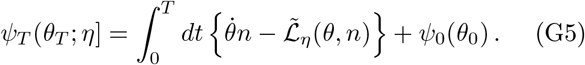

The stationary equations for *n* and *θ* on the solutions of which the integral (G5) is to be evaluated are

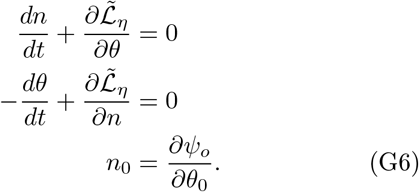

#### a. Identifying the saddle-point stationary path associated with a “nominal distribution” in the importance-sampling interpretation

As developed in [105], the field *n* that carries the saddle-point behavior of the large-deviation rate in this Hamilton-Jacobi theory has the interpretation of a saddle point for an importance distribution in importance sampling [139, 140]. The saddle point in the corresponding nominal distribution, which carries the interpretation of the underlying “state” deformed by the large deviation, is 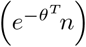, ^33^ for which the stationarity equation is

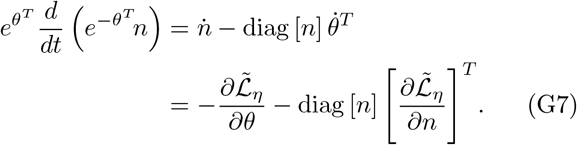

For many of the simple cases in which the trajectory equations are solvable in terms of mean-field dynamics, Eq. (G7) will have static solutions and will serve as an equation of constraint. This behavior reflects the anti-causal nature of the problem of inference [105] using the response field *θ* in the construction of Sec. IV A 4.

#### a. Matrix forms for the stoichiometric-process generator

Properties such as symmetry or stochasticity are more easily seen and more compactly denoted in matrix form, as we did in our discussion of balance classes in Sec. II C 2. Therefore define the matrix (*c*.*f*. fn. 23)

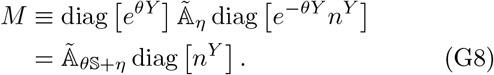

The two equations (G4) can be written in terms of *M* as

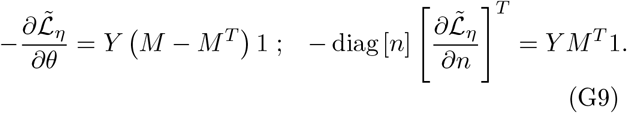

The gradient governing the behavior of the nominal distribution in Eq. (G7) is^34^

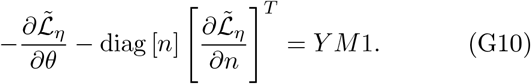

Antisymmetrization of *M* in the first equality in Eq. (G9) ensures that, for any values of *η, θ*, and *n* the gradient can be written in the form

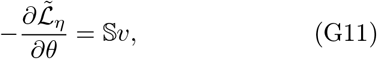

for some current vector *v*. It follows from Eq. (G11) that, for any *c* with *c* 𝕊= 0,

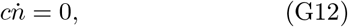

and *cn* is a conserved quantity under general deviations. The conserved quantities associated with transport and accumulation in the importance distribution are not changed from their values in the nominal distribution.

The relation (G12) entails a number of constraints on *n* at all times equal to the number of conservation laws. The first line of Eq. (G6) then, at steady state, provides *s* additional constraints acting jointly on *n* and *θ*.

#### b. Factoring spurious scale factors out of the CGF from fixed-point conditions

The conserved “phase-space” volume in the Hamilton-Jacobi theory for large deviations implies a duality between forward dynamics of nominal distributions and backward dynamics of the “response field” [100, 141] *e*^*θ*^ familiar from adjoint fluctuation theorems [142]. That duality is reflected in the symmetry of a pair of exponential-family coordinates, interchanged by a similarity-transform [105, 143] about the stationary state under the instantaneous generating parameters.

Introduce a dual exponential coordinate *ϕ* with the definition relative to the steady-state distribution *n* under the instantaneous parameters in ℒ at any instant:

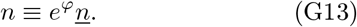

By Eq. (G12) *ϕ* respects *c* 𝕊= 0 ⇒ *c* (*n* − *n*) = 0 in all solutions.

To understand fixed points of trajectories in the CGF, we must first separate out dimensions of the tilting field *θ* that couple to conserved quantities *cn*, and which may have apparent dynamics in the CGF *ψ*_*T*_ but separate out into a total derivative that has no effect on the LDF *S*_*T*_. To do this, introduce a family of parameters 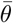, with only *s* degrees of freedom th at we may choose by requiring 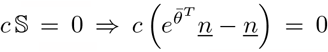 will provide a simple variable in terms of which to express gauge conditions for the field *η*. Now re-arrange Eq. (G8) and with it Eq. (G2) as

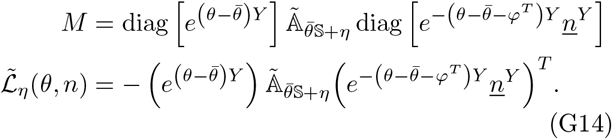

For any field configuration (*θ, ϕ*), assig(n a value to 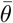 as a function of *θ* and *ϕ* with the condition 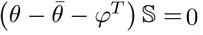, *i*.*e*. 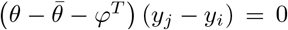 for any pair of complexes *j* and *i* connected by a reaction. These conditions fix the remaining *s* dimensions o f 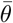 not set by the requirements 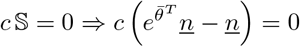.

The terms in 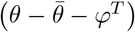 in the exponential matrices in *M* pass through 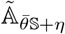 in each reaction term, so we may rewrite Eq. (G14) as

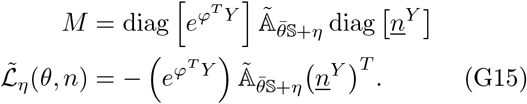

Thus *M* and 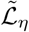 have as arguments only the fields 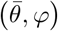 with 2*s* stoichiometric degrees of freedom. Any components of general *θ* ∝ *c* for conserved quantities *c* have canceled away.

**Remark:** A restricted but important subclass of cases are those with *θ* = *ϕ*, implying 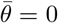. These are the tilts by *θ* that, in the absence of other forcing by *η*, reverse the current relative to mass-action flow for reactions with detailed-balance rate constants.

#### a. Dimension-counting of fixed-point conditions

The fixed-point condition for *n* given when 𝕊*v* = 0 in Eq. (G11), which can be written *Y (M* − *M*^*T*^)1 = 0, imposes *s* constraints on the 2*s* degrees of freedom in 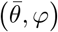. Take *ϕ* to be the independent variable in this constraint surface, and write the dependent field 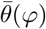.

To fix the remaining degrees of freedom in *ϕ*, look for a solution 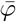 satisfying^35^

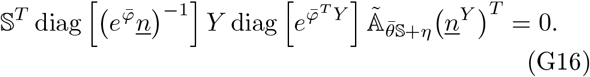

Eq. (G16) imposes *s* further constraints: that the vector acted upon by 𝕊^*T*^ is some linear combination of the conserved quantities of the stoichiometry.

#### c. Extraction of the nominal distribution in the stoichiometric compatibility class of the true dynamics

Eq. (G7) for the nominal distribution derives relative dynamics between *θ* and *ϕ* from a function depending only on 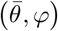 :

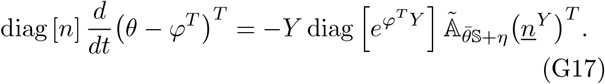

Because the conserved number components *cn* are non-dynamical, it is not possible in general to obtain a steadystate solution to Eq. (G17) for all components of *θ*. The most that can be required is 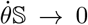 at steady state, so that a ny unbounded growth of *ψ*_*T*_ from components 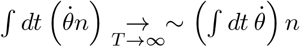 comes asymptotically from a total derivative that will subtract harmlessly away in the later Legendre transform (G46) to the LDF. (See [46] for a more didactic treatment in a simple example of these potentials multiplying conserved quantities in CGFs that include couplings to conserved number species.)

From the defining condition 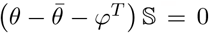 that projects a stoichiometric ray *θ* − *ϕ*^*T*^ to a representative point 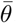, Eq. (G17) can be re-arranged to

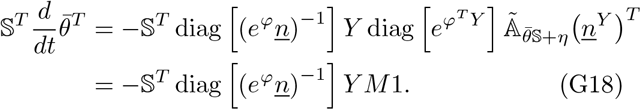

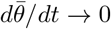 at the fixed point condition (G16). The corresponding condition in our original stationary-path equation from Eq. (G6) is

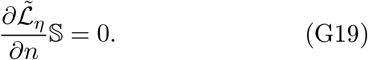

#### a.Nominal distribution within the stoichiometric compatibility class

Observe that the naïve nominal distribution evaluates to

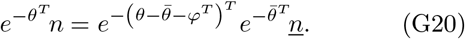

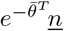 is the nominal distribution in the stoichiometric compatibility class of *<u>n</u>*. The logarithm of the weight term 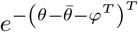 is proportional only to the conserved charges *c* of the species, and can be canceled out by a similar term in the likelihood function 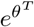. Hence we may factor the importance distribution into a likelihood and a nominal distribution entirely within the compatible stoichiometric subspace as

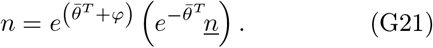

The log-likelihood 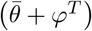 in Eq. (G21), like the chemical potential along an escape trajectory for the single-. time large-deviation function of Sec. IV A, will be the argument of the integral density for the functional LDF.

From Eq. (G21) we obtain a pair of closed equations:

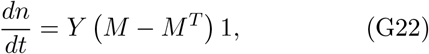

with Eq. (G18) for 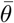 and Eq. (G15) giving the form of *M*. Although they are more complicated than the original dynamical equations(G6), and require the auxiliary condition 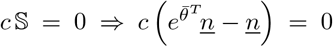, these are the stationary equations for a CGF acting only on the dynamically variable degrees of freedom in the system, and it is in terms of them that the LDF will assign deviation improbabilities.

We now turn to a sequence of constructions enabling us to solve for the Legendre-dual fields *η* or *θ* needed to produce target values of *v* or *n* by tilting.

#### d. Current reversal for steady states

On a closed graph 𝒢; the steady state solution to equations (G6) is necessarily a null flow: the velocities (G3) satisfy 𝕊*v* = 0.

**Lemma:** The tilting vector *θ*𝕊 + *η* that reverses the steady-state velocity is also a steady-state solution to equations (G6) at the same profile *n*.

**Proof:** First, note that since *η* is free at every reaction, a solution *θ*𝕊 + *η* always exists that reverses the steady-state flow as long as reactions in both directions exist. By Eq. (G4), the first line of Eq. (G6) is 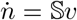, which is zero for both directions of a null flow. Denote by 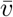 the steady-state velocity at *θ*𝕊 + *η* = 0. Some algebra, making use of the second line of Eq. (G2), shows that at the value of *θ*𝕊+*η* that reverses the steady-state velocity,

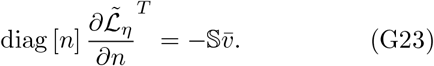

While no longer identically zero as it would be at *θ*𝕊+*η* = 0, the expression (G23) is still zero for 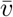 null, giving in the second line of Eq. (G6) that 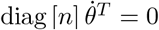. We may check also that 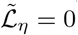 at this solution. ■

This result will be useful in relating the large-deviation function for flows through a subgraph driven by external sources, to the usual physical notions of dissipation within the subgraph at the the profiles *n* and currents *v* of non-equilibrium steady states.

#### e. Time-reversal of general solutions in graphs with detailed-balance rate constants

From the form (14) for the contribution from each complex pair (*ji*) connected by a reaction in a network with detailed-balance rate constants, the Liouvillian (G2) becomes

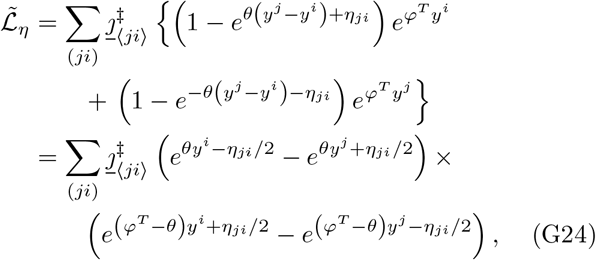

and the current equation (G3) derived from it becomes

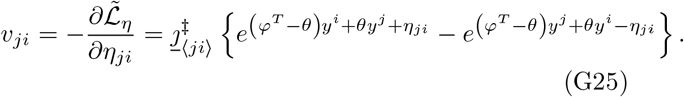

**Lemma:** In the detailed-balance case, the assignment *θ* = *ϕ*^*T*^, *η* = 0 is the velocity-reversing solution from *θ* = 0, *η* = 0, here not only for steady states but for general time-dependent trajectories.^36^

**Proof:** The *θ* = 0, *η* = 0 trajectory solves the classical mass-action rate equations at concentrations *e*^*ϕ*^*n*. As noted in the previous lemma, the second line in Eq. (G6) for 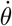, no longer identically zero, becomes the same as the first line for 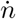 at reversed *v*. Therefore *θ* = *ϕ*^*T*^ is also a stationary solution. On both the classical and time-reversed solutions, 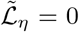 in Eq. (G24) as well. ■

**Corollary:** The assignment *θ* = *ϕ*^*T*^ implies vanishing of the time-derivative of the nominal distribution in Eq. (G7), even if *n* and *θ* are time-dependent. Thus the large-deviation likelihood reflects biased sampling of an unchanged nominal distribution, a general characteristic of *θ* = *ϕ*^*T*^ solutions on detailed-balance backgrounds [32]. While on a closed graph, the steady states are equilibria and are unique [22], so *ϕ*^*T*^ ≡ 0 and 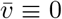, the above result also applies to steady states of subgraphs driven by sources, which will be its primary use here.

#### f. Variations of the CGF and functional Legendre transform to the LDF

The first-order expansion of the integral expression (G5), making use of the fact that *θ* and *n* satisfy the stationarity conditions (G6), is

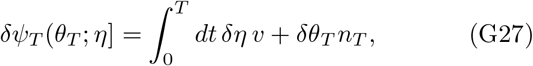

giving both the final-time and the variational derivatives

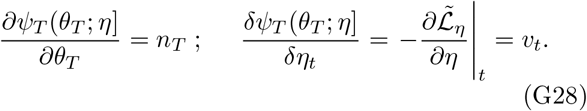

The second variational derivative is computed as

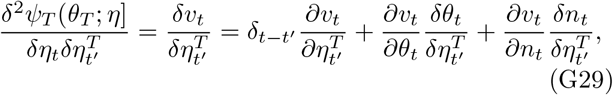

in which 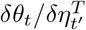 and 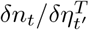 are the variations in the trajectory solution (G6) at any time *t* resulting from variation of *η* in 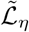 at a time *t*′.

The large-deviation functional dual to *ψ*_*T*_ (*θ*_*T*_ ; *η*] is the functional Legendre transform

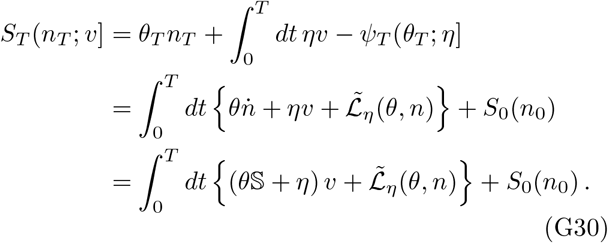

The form in the third line of Eq. (G30) uses Eq. (G6) to substitute 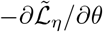 for 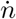, and then Eq. (G4) to evaluate 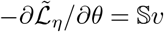. These results are transcribed as Eq. (68) and Eq. (69) in the main text.

**Remark: large deviations of dynamical quantities only**

In the third line of Eq. (G30), 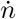 is replaced with 𝕊*v* by the first stationarity co1ndition (G5) anld Eq. (G11). In it, we can evaluate 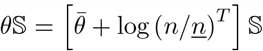. Then, with the cancellations connecting Eq. (G14) to Eq. (G15), 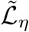 is also a function only of 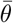 and log (*n/n*). The LDF is thus a function only of the nominal and tilted dynamically variable degrees of freedom in the system. As is common for manipulations of generating functionals with conservation laws (see [46]), the algebra remains simplest in the variables (*θ, n*), at the price of their possibly including spurious non-stationary degrees of freedom.

The first variational derivative of *S*_*T*_ follows from the construction of the Legendre transform as usual:

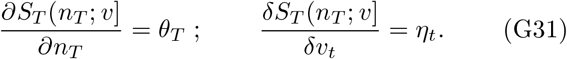

The relation between the second variational derivatives of the CGF and LDF generalizes that for the single-time construction of Sec. IV A 2, to

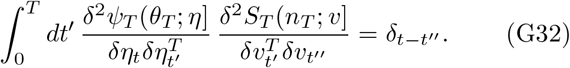

We therefore write as a matrix inversion, leaving the time indices as part of the matrix indexing, that

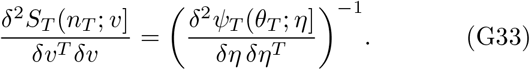

### 2. The LDF as a functional Bregman divergence

The large-deviation functional directly extends the Bregman-divergence form of the Legendre transform and its integral representation from the single-time construction in Sec. IV A 4. We may write the counterpart to the twice-integrated kernel in the local Fisher metric (58) both directly for the functional Hessian of *ψ*_*T*_, and in steady states as the integral of a time-local integrand of similar form in which the Hessian of the Hamiltonian 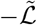 appears as the metric.

#### a. Simple starting case: steady-state solutions

We may wish to compute an improbability-cost for a shift in a steady state, which can be assigned to the boundary conditions causing the shift. In this case the large-deviation functional reduces to a simple time-integral of functions of partial derivatives. For convenience in the presentation of this case, we will not regard *n*_*T*_ as an additional argument of *S*_*T*_ distinct from its value in the shifted steady state. If we further absorb the stationary value of *θ*S into *η* by a gauge choice, to facilitate computation of the LDF as a Bregman divergence in its natural argument *η* of the CGF *ψ*_*T*_, the integral form (G30) evaluates to

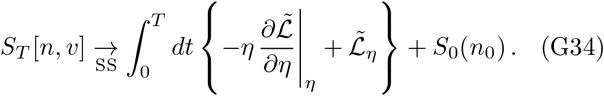

Although we write *η* as a single argument to the CGF, it is dual to both arguments *n* and *v* of 𝕊_*T*_. This is so because on a closed graph, steady-state solutions for *v* are constrained to 𝕊*v* = 0, and thus have the dimension of the space of null flows. The remaining dimensions of *η*, equal in number to the stoichiometric dimension *s*, are fixed in terms of the stoichiometric dimensions of *n* by the *θ*𝕊 = 0 gauge choice under Eq. (64).

In steady states the full time-dependence of the Hessian (G29) of *ψ*_*T*_ is not used, and only a time integral on one index appears, which evaluates to

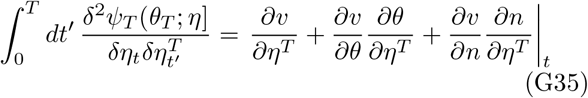

in which *∂θ/∂η*^*T*^ and *∂n/∂η*^*T*^ are now the ordinary responses of the stationary-point values to change in *η*.^37^

Denote any background current in the steady state of the generator by

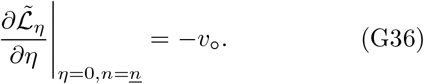

The flow-velocity variation that appears in the second derivative (G35) is given by

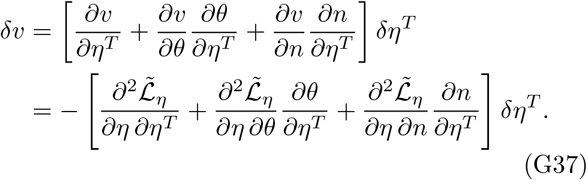

By taking the partial derivative of the argument in Eq. (G34), including the effect of *η* on the stationary solution (*θ, n*), we convert the Bregman form to the integral

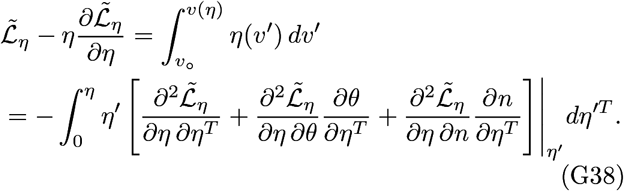

Under the stationarity equations (G6), the second and third terms in the square brackets vanish, and the second variational derivative becomes manifestly square and time-local, giving Eq. (75) in the main text. The *θ*-Hessian of the single-time CGF *ψ* in Eq. (58) has been replaced by the *η*-Hessian of the Hamiltonian 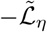 for steady-state tilting to drive null flows. The steady-state LDF (G34) is just the integral

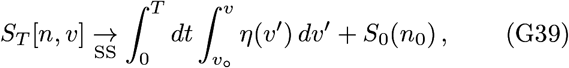

evaluated through Eq. (G38), and transcribed as Eq. (74) in the main text.

**Remark:** The expression (G38) is a sum over terms identified with the individual reactions. In the next section we will consider the partition of a graph 𝒢 into a subgraph 𝒢′ and its complement (𝒢′)^⊥^. Since every reaction in 𝒢 is either in 𝒢′ or in (𝒢′)^⊥^, the likelihood cost (G39) partitions into two summands corresponding to the restrictions of the Hessian in Eq. (G38) to 𝒢′ and (𝒢′)^⊥^.

#### b. Functional Bregman divergence: the general form

A more laborious but otherwise identical calculation can be carried through for general time-dependent *n, v*, and *η*. Taking the variational derivative of the argument in the final line of Eq. (G30), through its manifold of stationary solutions, we construct the equivalent form in terms of the time-dependent second variational derivative (68), as

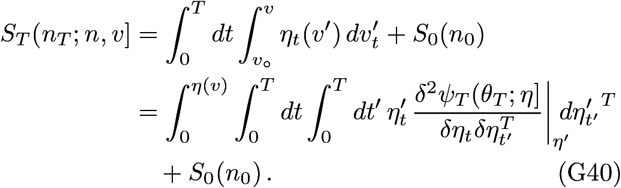

Eq. (G40) is the functional generalization of the integral representation of the single-time Bregman divergence in Eq. (58).

### 3. eparation of tilted generating functions between a subgraph and its environment

The observation that the LDF (G39) is a sum over reactions, and thus already additive under graph partitioning, suggests that the only additional work required to define a LDF for subgraphs with open boundaries is to identify the correct separating hyperplane to make stationary-path solutions conditionally independent between a graph and its complement, given the boundary conditions. This subsection provides that required construction.

Following the development from Sec. IV B 5, we consider flows 𝒢 supported by reactions in both 𝒢′ and (𝒢′)^⊥^. As in the text, denote by 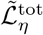 the generator on 𝒢, and by 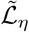 the restriction to 𝒢′ and 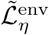 the restriction to (𝒢′)^⊥^. 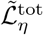is then the sum over the two subgraphs

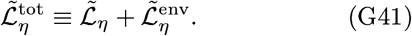

Gradients with *η* separate between the subgraphs, while gradients of both 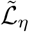 and 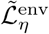 with *θ* and *n* produce terms on *∂*_𝒢_ 𝒢′.

Along any particular stationary trajectory, we may choose a column vector of species-fluxes *J*^ext^ and a row vector 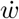 with the dimension of rates of chemical work per particle, nonzero only on the boundary species,^38^ that define a separating hyperplane between 𝒢′ and (𝒢′)^⊥^. Adding and subtracting these separating terms gives

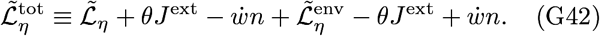

If we choose values for *J*^ext^ and 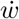such that the variation of 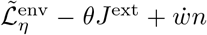 along the stationary trajectory vanishes, the variation of the remaining terms in Eq. (G42) modifies the equations of motion (G6) for (*θ, n*)| _𝒢′_ to

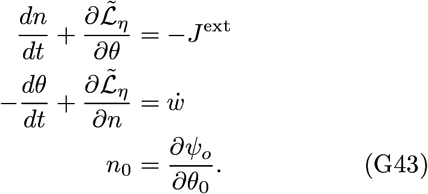

In Eq. (G43) and henceforth we drop the explicit notation for restrictions |_G_′, having already stated that the fields under consideration are evaluated on the subgraph.

The variational equations (G43) are those obtained from a CGF on the subgraph 𝒢′ of the form

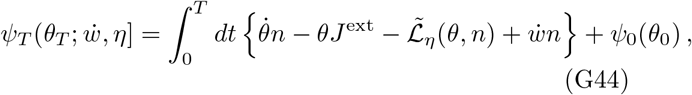

given in the text as Eq. (81). The first-order variation of 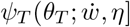 generalizes from Eq. (G27) as

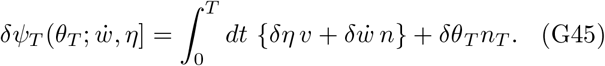

The LDF dual to Eq. (G44) is then

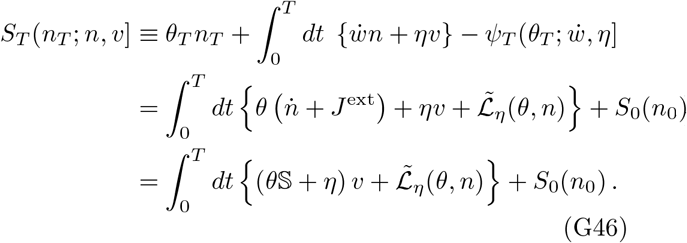

As in the computation of Eq. (G30), the stationarity equations replace 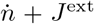 with 𝕊*v*. Eq. (G46) is just the restriction of the form (G30) to the reactions in 𝒢′.

Direct variation of the second line in the integral expression (G46), together with use of the stationary equations (G43), along any contour of boundary conditions 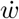 and internal tilts *η*, returns the result that the Legendre transform is constructed to yield:

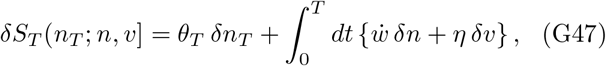

from which the first partial and variational derivatives follow:

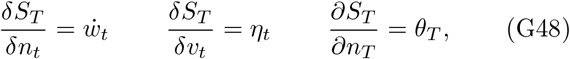

as well as the Hamilton-Jacobi equation

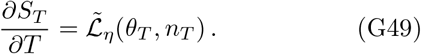

We repeat the observation in the text that the extension of the gauge invariance (64) for null flows in 𝒢, to flows in 𝒢′ fed by source currents *J*^ext^, becomes

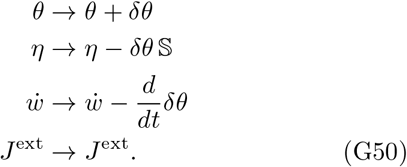

The transformation (G50) is manifestly a symmetry of the integral expressions for *ψ*_*T*_ and *S*_*T*_ and of the stationary equations (G43) for the trajectories in those integrals.

#### a. Relation of J^ext^ and 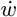 on the boundary of a driven subgraph

Again using Eq. (G4), the first line of Eq. (G43) becomes

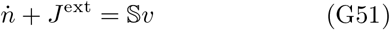

at any values of *θ, η*. From the form (G2), at 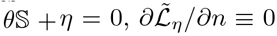, implying that in the second line of Eq. (G43), 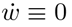 for classical driven steady states.

If we denote 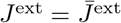 in a steady driven state at *θ𝕊*+ *η* = 0, we can extend the notation from Eq. (G23) of 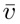 for closed graphs to the solution to 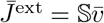, both from external sources and from any any breaking of detailed balance in the rate constants internal to the graph.

For *θ𝕊* + *η* the time-reversing solution, we then have 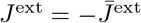 and 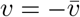. The evaluation Eq. (G23) for he gradient of 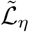 then gives the second line of Eq. (G43) the form

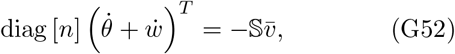

the negative of Eq. (G51). Recalling arguments in Sec. G 1 b that 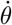 may be non-vanishing only in proportion to conserved quantities, to make (*n, v*) a current-reversed steady state, the boundary conditions must satisfy

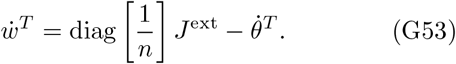

The in tegrand in the CGF (G44) becomes simply 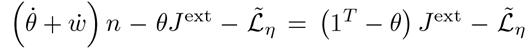 and is a function only of time-invariant variables. Not only the possibly non-stationary terms in 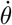 proportional to conserved quantities, but the entire term 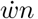, cancels in the Legendre transform to the LDF (G46).

Solutions with the same *J*^ext^ that are gauge-equivalent to *η* ≡ 0 will not generally preserve the profile *n* in networks without detailed balance. For those, the relation of *J*^ext^ to 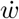 generalizes from Eq. (G53), by subtraction of the first two lines of Eq. (G43), to

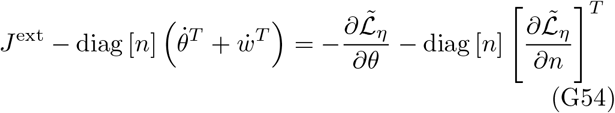

The right-hand side equals *Y M* 1 from Eq. (G10) and Eq. (G7), controlling the time-evolution of the nominal distribution, which vanishes identically for source-reversing solutions in networks with detailed balance.

### 4. Pythagorean decomposition of the Bregman divergence

We may now study the interaction of null flows within a subgraph, with conservative flows through it, and in particular express any conservative flow as a sum of a component with the same net conversion *J*^ext^ that is gauge equivalent to *η* ≡ 0 and a component that is null in the subgraph. The final overall tilt is decomposed into two stages: First, use 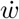 to arrive at a current *J*^ext^ in the absence of any tractions *η* internal to the subgraph. Then, hold that through-current *J*^ext^ fixed as *η* is introduced, passing only through configurations that differ from the *η* = 0 starting point by a null flow. A convenient use of the gauge symmetry in this decomposition is to hold *θ* fixed during the *δη* /= 0 transition, not at 0, but at the value *θ*(*J*^ext^) produced in the subgraph when 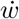 is tuned to yield current S*v* = *J*^ext^ at *η* = 0 in the subgraph.

To separate the transition from initial *v*_*°*_ and *n* of the untilted generator ℒ to some final profiles *v* and *n* into these two stages, we first partition the Legendre term in the integrand of the LDF (G46) as

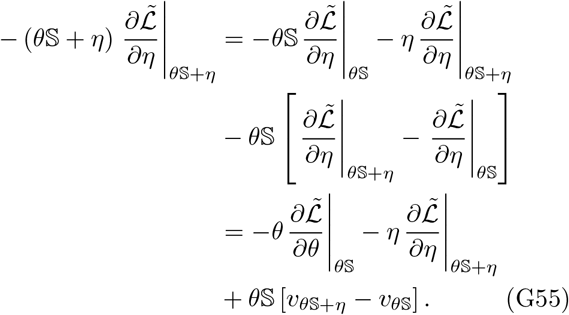

By choice to fix *J*^ext^ when introducing *η* in the square brackets on the left-hand side of Eq. (G55), the current difference [*v*_*θ𝕊*+*η*_ − *v*_*θ𝕊*_] is a null flow, meaning that 𝕊 [*v*_*θ𝕊*+*η*_ − *v*_*θ𝕊*_] ≡ 0.

The argument of the integral in the final line of Eq. (G46) is then first written as a Bregman divergence in standard form by recognizing that 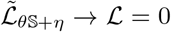 at *θ𝕊*+ *η* = 0, and then decomposed into a sum of two other Bregman divergences for the two stages of transformation using Eq. (G55), as

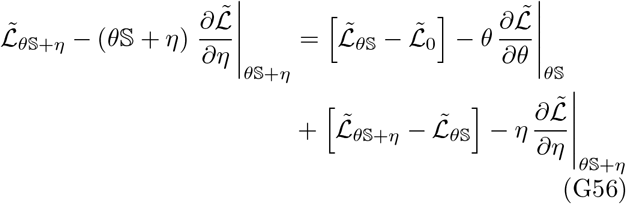

The absence of any cross-term, making the Bregman divergence on the left-hand side of Eq. (G56) a sum of two such divergences on the right-hand side, gives the required extended Pythagorean theorem. In the dual geometry induced by the Hessian of the CGF (G44), the configuration at *θ𝕊* is the nearest one in the manifold that can be induced by boundary values 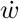 and *J*^ext^ alone, to the final profiles *n*_*θ𝕊*+*η*_ and *v*_*θ*S+*η*_. Any remaining divergence extends within the manifold of null flows at fixed *J*^ext^.

#### a. Integrated Bregman form in steady state

The integrand in the functional LDF for steady-state currents can be converted to the same double-integral form as the single-time LDF (58) for counts, with 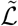 taking the place of *ψ* as the convex function from which a Fisher metric is defined. Recalling the final line of Eq. (G30), the different meaning of the two legs of the Pythagorean decomposition, as respectively potential-dependent and explicitly *η*-dependent, can be made manifest, and at the same time we may express all geometric quantities in terms of the non-diverging tilts within the stoichiometric subspace from App. G 1 b.

Using 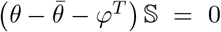 and the relation 𝕊*dv*^*′*^ = *dJ*^ext*′*^, the two legs of Eq. (G56) become

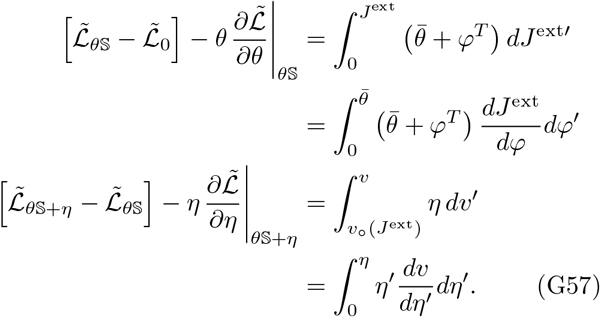

In the first line of Eq. (G57), 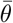 is a function of *ϕ*, which is identically zero for the case of detailed balance. In the second line, we use the fact that 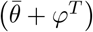 is a potential, and *dv* at every stage is an increment of a cyclic flux if the boundary flux *J*^ext^ is fixed. Cyclic fluxes have no inner product against the drops from a potential. So even though that potential will in fact be changing as we increment the cyclic flux, that change of potential makes no difference to the second Bregman divergence.

## Footnotes

1 We note in passing another work that, while not related to ours in representation or method, also studies the interaction of ther-mochemical characteristics with molecular combinatorics. Jinich *et al*. [41] make a computational library of small linear-chain molecules and explore the formation free energy and reaction opportunities for all distinct concatenations of oxidation states of the carbon centers, finding that biological metabolites are enriched among more stable and more soluble compounds relative to a uniformly sampled library. They omit bond-forming and breaking reactions and thus molecular re-arrangement, and consider phosphorylated (or aminated) metabolites equivalent to their alcohol counterparts in oxidation state, and in that sense pursue a different origin of combinatorics from the one we pursue with graph grammars.

2 More complex notations are available defining the hypergraph as a tuple of sets of species, complexes, and reactions or subsets of these [68]. Since reactions entail complexes and complexes entail their species members, the abbreviated notation suffices, and it provides a more compact subset definition for sub-graphs.

3 The notation is motivated by the action of a Taylor’s series for the exponential on functions of a continuous argument, though here the indices n take only discrete values. Keeping only the first two terms in such an expansion recovers the correct functional form for the diffusion approximation in the limits of large numbers n, and other useful transformations [29, 69].

4 The mass-action rate law is a form of *mean-field* approximation. Formally it is a leading-exponential approximation for expectations of factorial moments, and as such equivalent to the saddle-point approximation that defines large-deviation limits [43, 45]. These approximations are generally regarded as “un-regulated”, in the sense that they are not accompanied by perturbative small-parameter expansions that are assured to converge.

5 It must be noted, however, that the time rate of change in a steady state is also zero, so the mass-action law gives only the currents in the complex network. For a general ACK state, not necessarily the fixed point of the generator, Eq. (7) will still hold, but the ACK form will not be preserved except in special and much more restrictive conditions on the underlying network [71].

6 If we were describing species with concentrations rather than explicit integer molecule counts, these would be standard-state chemical potentials.

7 These may be more elaborately terms *balanced integer flows* or *balanced integer hyperflows*, with no change in meaning.

8 In mean-field approximation, these first moments are solved from the mass-action relations (7), and no higher-order moments are resolved. In an exact solution, the mean values may differ from mean-field values, and higher-order moments will generally also have values not derivable simply as functions of the first moments. See [39, 40] for approaches to such cases.

9 The definition here of *feasible processes* provides a graph-defined alternative to the notion of *chemical processes*, put forth in Wachtel *et al*. [1], which are defined only in terms of respect for conservation of elements (including separate treatment of electrons and protons) in the chemical formula.

10 One can view the molecular “turnstyles” and “escapements” [76, 77] that transduce free energy between processes that are not sto-ichiometrically coupled by chemical necessity as evolved mechanisms to impose high kinetic barriers along sequences of reactions coupled to a succession of overlapping inputs and outputs [78].

11 An example is the schema 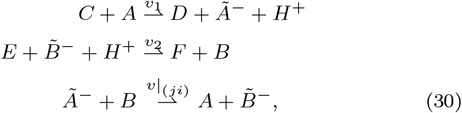 in which *A* and *B* are acids, 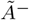 and 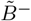 are their respective conjugate bases, perhaps with pKa changed by the reactions *v*_1_ and *v*_2_, which might be other redox reactions, and *H*^+^ is the proton. We think of chemical potential from *C* to *D* as driving the system through *v*_1_, and chemical potential from *E* to *F* as being received from the system through 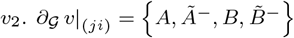. *∂*_*G*_ (*v*_1_ + *v*_2_) contains these species and also {*H*^+^, *C, D, E, F}*, in which *H*^+^ may receive fluxes from *v*_1_ that are delivered to *v*_2_.

12 It is derived in Sec. IV that this result in the linear-response regime is Onsager’s [81, 82] minimum dissipation principle.

13 If *n* = 3 then *A*_*n*_ ↔ GAP and we may cancel products in the schema (36) to yield 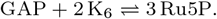 If *n* = 2 then K_*n*+3_ ↔ Ru5P and we may cancel reactants in schema (36) to yield 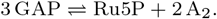

14 Cyclic permutations are the only possibility. The identity map on (*m, n, p*) is no reaction. Exchanges of two elements, fixing the third, are 2-cycles which are self-nullifying by TAL^2^ = *I*. One can verify that the two non-trivial cyclic permutations correspond to opposite orientations of the same cycle.

15 *A*_0_ has the stoichiometry of the realizable molecule HPO_3_. But in implementations where the free energy landscapes for molecules in solution are required, this molecule cannot take the charge states typical of all realizable *A*_*n*≥2_. Therefore the identification (39) is only a formal encapsulation of rule symmetries.

16 An original list of 800 solutions by the ILP solver produced 37 that duplicated flows earlier in the list, through un-trapped cancellations of current through two opposite unidirectional representations of the same reaction. These duplicates are not included in the plot. The index *i* from the original ILP list is shown, without renumbering.

17 Wachtel *et al*. [1] frame their definition of transduction by starting from pairs of *independent* chemical processes in the environment, so that each member has a defined forward direction under the classical Second Law of thermodynamics. Nothing in our definition of feasible flows and transduction makes specific reference to whether the transformation decomposes in the environment into independent smaller feasible transformations.

18 As a corollary, we see why solutions with mixed preludes can reach lower values of 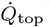 than those with “pure” preludes that use two aldol condensations of the same aldose input. Potentially greater genomic complexity to encode two specific enzymes is exchanged for lower dissipation in the operation of the cycle.

19 If *ψ* is twice differentiable the inverse exists everywhere; if *ψ* is once differentiable with discontinuous derivative, *n*(*θ*) may have gaps in its range; see [45] for treatment of these cases that we will pass over in the very brief summary presented here.

20 Its generalization to cases where *S* has multiple basins, requiring the treatment of instantons in a full saddle-point treatment, is explained in [32].

21 We may say that the role of energy change is to transduce like-lihood, from one large-deviation rate to another.

22 This simplification is in keeping with the interpretation of tilts in terms of buffered fictitious species. The general case with both symmetric and antisymmetric components of *η* adds the feature of modifying the heights of reaction transition states.

23 Indeed we could further compress the expression (60) for the generator to See Eq. (G8 in App. G 1.) 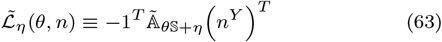

24 The derivation in App. G 3 a shows that, if there is any nonzero part of 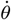 in a steady state, it is proportional to *c* with *c* S = 0 and does not contribute to the evaluation of either the CGF or LDF.

25 We assume here that the solutions to the stationarity equations cover the full space of currents. Cases in which the stationarity equations are incomplete for dynamical solutions [70, 70, 107– 109] create a typology of special cases, and we do not try to find general proofs of the conditions for completeness in this paper.

26 More generally, as detailed in App. G 4, *θ* and *n* pass through the trajectory defined above: starting from the velocity-reversing solution that preserves *n* from the mass-action solution, then altering purely null flows and *n* until *η* = 0 up to gauge equivalence.

27 For networks with detailed balance the forms of both legs of this decomposition are straightforward to interpret. For the more general case, equivalent expressions are given in Eq. (G57), which show that the geometric factors remain functions only of finite quantities defined within the stoichiometric subspace through the reference background *n*.

28 Another way to say this is that all information about the topology and through-current *J*^ext^ has already been incorporated in the choice of basis {*f*_*µ*_} for flows on the subgraph 𝒢^*′*^, and *h* ∝ 1 contains no further information about the ensemble.

29 The matrix 𝕊 diag[*j*^*‡*^ ] 𝕊^*T*^ has a null space spanned by the conserved quantities of the stoichiometry, so the matrix inversion to solve for *θ*^*T*^ = log (*n/n*) and thus *v*_∥_ from *J*^ext^ must be performed keeping *n* in the stoichiometric subspace of *n*, reviewed in App. F).

30 It has, however, been found to enable a novel non-oxidative glycolytic pathway in yeast [129].

31 The number of faces on the simplex in *D* dimensions is (*D* + 1)-choose-3. However, cycles around these faces are not all independent, due to cobordism relations. The number of faces always exceeds the number of independent flows by a factor (*D* + 1) */*3, with equality at *D* = 2.

32 These 12 solutions make up 6 pairs among the 19 pairs with duplicated graphs, accounting for 6 supporting graphs out of the 66 distinct graphs for reducible flows. Each of these 6 add a “superfluous edge” to one of the graphs for an irreducible flow, not activated in the minimum-dissipation solution, which recapitulates the irreducible flow.

33 The bias factor (called the likelihood function) for sampling is then just the tilt 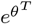.

34 The nominal distribution thus corresponds to a distribution in coherent-state coordinates, in place of the number-potential coordinates (*θ, n*). See [32, 46, 69] for discussion of properties in both sets, and of the anticausal dynamics of coherent-state response fields in the large-deviation function.

35 Proofs of existence or conditions for uniqueness of solutions to these nonlinear fixed-point equations are an exercise in convex analysis similar to those in [20]. We will assume here that such solutions exist in some neighborhoods of *η* about given fixed points *n* of the unperturbed system, and construct explicit examples to justify the assumption.

36 The assignment *θ* = *ϕ*^*T*^ also gives escape paths for more general complex-balanced rate constants, in which case the time-dependence of the escape ray is given by 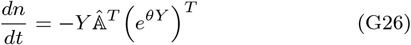 Comparing to Eq. (7), the EOM for free relaxation, where the term 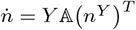, we see that escape rays behave as the time-reverses of relaxation trajectories under the transpose of 𝔸.

37 The full form including *θ*-dependence is written in Eq. (G35) for completeness, but in the gauge surface *θ* ≡ 0, terms from the variation of *θ* will of course be zero.

38 As we saw in Sec. G 1 c for *θ*, there may be terms in 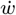 that drive components of 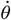 within 𝒢 ^*′*^ because they couple to conserved quantities, but which cancel in the Legendre transform to the LDF. The components of 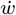 constraining the stoichiometric components of 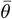 and *ϕ* are nonzero only on species in *∂*_𝒢_ 𝒢 ^*′*^.

